# Branch-Partitioned Selection Tests Reject Random Labels on Real Genes, Even Against a Null Simulated at the Depth of the Data

**DOI:** 10.64898/2026.07.20.739482

**Authors:** MinSeo Kim, Jae-Ho Shin

## Abstract

Comparative genomics often turns a trait into a hypothesis about selection by labelling the branches that carry it and testing them against the rest. We asked how such a test behaves when the labels carry no trait. In three panels of about 50 species, fungi, yeasts and lactobacilli, two of them registered in advance, randomly placed terminal labels on presumed-null single-copy genes were rejected by PAML’s two-ratio branch model in 65.9, 68.8 and 88.8% of genes, against a nominal 5%. Referring each real gene to replicates simulated under its own fitted null model did not remove the excess: with a generator matched to the depth of the real alignments, 65.0% (95% CI 53.5–75.3) and 83.8% (73.8–91.1) of 80 genes were still rejected. On the same genes the branch-site test, which asks only for positive selection on the labelled branches, rejected none after refitting, though on alignments simulated from its own fitted model it detected positive selection placed on all those branches in 95–100% of replicates, and replicates simulated under each gene’s fitted per-branch *ω* rejected 1 of 40 genes against 28 of 40 for a control carrying only the noise of that fit: *ω* plausibly varies among branches on real genes, so that a randomly chosen set of branches does differ from the rest, though the fitted pattern is an upper bound. Under correct specification at the depth of the data the same test rejected random terminal labels in 5.4 and 6.8% of replicates, and internal labels of matched summed length differed from them by +0.2 and -1.2 percentage points. On earlier simulations from a site-mixture model, moving the labels from terminal to internal branches raised the rejection rate 1.3-fold on genes with a moderate fitted *ω* and 6.2-fold where a fitted class reached *ω* ≥ 100, in interaction with the fitted model. A trait label should therefore be read against random labels of the same size rather than against χ^2^, and a reference simulated from the fitted model is not enough. The count question asked of the fungal panel is answerable: the within-genome ratio of plant-cell-wall-degrading enzymes to other carbohydrate-active enzymes is 0.473 times as large in ectomycorrhizal genomes as in free-living decomposers.

## INTRODUCTION

Ectomycorrhizal symbiosis has arisen repeatedly across the fungal tree, and in the lineages examined genomically the same signature has appeared repeatedly: the complement of enzymes that degrade plant cell walls is reduced relative to free-living decomposers (Kohler et al., 2015; Miyauchi et al., 2020). It is one of the clearest cases of convergent genome reduction.

What the observation means is less settled. The reduction is usually read as the loss of a capacity. These fungi, having secured carbon from a host, no longer decompose. Against that reading stands the argument that ectomycorrhizal fungi retain the capacity to oxidise soil organic matter and deploy it to mobilise nitrogen rather than to acquire carbon (Lindahl and Tunlid, 2015), a position whose authors are explicit that it is not a claim of facultative saprotrophy, and which transcriptomic and spectroscopic evidence of oxidative decomposition has since supported (Shah et al., 2016). The two readings predict different soil carbon dynamics, and the genomic evidence has been recruited by both.

The difficulty is that the evidence is a count. A gene may be present and never transcribed; a transcript may not be translated, secreted, or active on the polymer that matters. Counting establishes what a genome could in principle encode, but the step from there to what an organism does is not supplied by counting more genomes. That gap between gene content and functional capacity is the question this study set out to address, and we do not close it.

The natural bridge, and the one we attempted, runs through selection: copies that remain functional should be maintained by purifying selection, and copies en route to loss should show relaxed selection. RELAX (Wertheim et al., 2015) makes this comparison directly. The analyst partitions the tree into a test set and a reference set, and one parameter, *K*, describes whether selection on the test branches is relaxed or intensified relative to the reference. Because it needs no experimental data and turns a categorical trait such as a lifestyle directly into a hypothesis about selection, it is widely used, and where the trait is measured on extant organisms the test set consists of terminal branches and the reference set contains everything else.

That is the configuration whose behaviour we report, in RELAX and in two further implementations of the same comparison: the two-ratio branch model of codeml (Yang, 2007), which carries no *K* and whose fits on these data from two starting values agree to within 10^-4^ log-likelihood units, so that the headline measurements below come from it, and Contrast-FEL (Kosakovsky Pond et al., 2021), which works site by site. We applied the configuration to plant-cell-wall-degrading enzyme families across a large ectomycorrhizal panel and reported in version 1 of this preprint (Kim and Shin, 2026a) whether the surviving copies show maintenance or decay. This manuscript reports what we found when we tried to establish that the instrument was measuring what we thought.

We ask two questions of one dataset. The count question, how the number of these genes varies with lifestyle once shared ancestry is accounted for, is answerable. The selection question, in this configuration, is not, and the reason lies in what a branch-partitioned test asks. Its null hypothesis is that the labelled branches share *ω* with the rest. If *ω* varies among branches on real genes, that null is false for almost any set of branches, chosen for a trait or at random, and a test that detects the difference answers a question about variation in *ω* rather than about the trait. On real null genes in three panels, two of them registered in advance, randomly placed terminal labels were rejected by the two-ratio test in 65.9 to 88.8% of genes; a reference simulated from each gene’s fitted null model, even at the depth of the data, did not remove the excess, whereas one simulated under each gene’s fitted per-branch *ω* did. That the RELAX estimates also fail to reproduce across optimiser settings that should not matter is documented in a companion manuscript (Kim and Shin, 2026b).

We offer no new estimator and no correction factor, doubting that any constant could hold across designs when our own calibration moved with the number and placement of the labelled branches and with the fitted model. What we offer is the count result, with three corrections to our own lifestyle assignments found by auditing our data against the primary literature; a demonstration that random labels of the size of the trait set are rejected on real genes, so that a trait label must be read against such labels rather than against χ^2^; a characterisation, on data simulated with no difference between branch sets, of how the error rate moves with where and how many branches are labelled; and a re-examination of our own preprinted inference, which used none of these checks.

## MATERIALS AND METHODS

### M1. Genome panel, quality control and orthology

Fungal genome assemblies and predicted proteomes were obtained from the JGI MycoCosm portal and from NCBI, placing ectomycorrhizal (ECM) symbionts alongside free-living and other symbiotic comparators. BUSCO 6.1.0 in protein mode (Manni et al., 2021) against the fungi_odb10 lineage dataset left a 183-genome panel, and OrthoFinder 3.1.5 (Emms and Kelly, 2019) inferred orthogroups and a species tree placing 182 of the 183. Phylogenetic models and every phylogenetically corrected comparison use the 182 placed genomes; two descriptive comparisons in Results 1 use all 183 and are labelled as such (Supplementary S1). Lifestyle assignment and its audit are described in M6.

### M2. CAZyme annotation and the definition of PCWDE

Carbohydrate-active enzymes were annotated with run_dbcan 5.2.8 against the dbCAN HMM library (Zheng et al., 2023) at family and subfamily resolution, subfamily activities following Aspeborg et al. (2012). Plant-cell-wall-degrading enzymes (PCWDE) were defined following CAZy (Lombard et al., 2014) as the 23 glycoside hydrolase, polysaccharide lyase, carbohydrate esterase and auxiliary-activity families directed at plant cell wall polysaccharides, a protein counting once per family it carried (Supplementary S1). AA9 is counted but excluded from catalytic-residue scoring and from every selection analysis, for the reason given in M3, and the count comparison was repeated with it removed.

### M3. Catalytic-residue scoring

Four PCWDE families (GH5, GH6, GH7 and GH28) have a characterised reference enzyme carrying UniProt ACT_SITE annotations. Their members were aligned with MAFFT 7.526 (Katoh and Standley, 2013) together with that reference, and each copy was classified at the catalytic columns as intact, disrupted or fragmentary; the functional PCWDE count used in the comparative analyses is the number of catalytically intact copies in those four families per genome (Supplementary S1, S2.5). AA9 could not be scored: the N-terminal histidine of its His brace is the first residue of the mature chain, so in proteomes that retain signal peptides it does not occupy a homologous alignment column; a SignalP 6.0 check (Teufel et al., 2022) is reported in Supplementary S1.

### M4. Codon alignments, the identifier bridge, and the analysed entries

Coding sequences were retrieved from JGI MycoCosm for 51 of the 183 genomes, 21 of them ECM, so every codon-based analysis is restricted to that subset, an availability boundary rather than a sampling design. CDS were joined to the scored proteins on the gene model name and checked by translation, because JGI’s numeric identifiers pair unrelated records without any error (Supplementary S1).

Per-entry protein alignments of the retained copies were back-translated to codon alignments, an *entry* being one family or subfamily alignment tested as a unit; columns with more than 50% gaps were trimmed, and gene trees were inferred with FastTree 2.2.0 (-nt -gtr; Price et al., 2010). Of 28 entries attempted (twenty families, seven GH5 subfamilies and one subsampled GH5 alignment), 27 returned an interpretable statistic under both starting-point settings, and every count of analysed entries in this paper is that 27; the twenty-eighth, GH5_22, is reported in the companion manuscript (Kim and Shin, 2026b, Results 3). Only GH5, GH6, GH7 and GH28 were restricted to catalytically intact copies, the rest being filtered on length, so the length-filtered entries should be read as upper bounds on relaxation and the four reference-anchored entries as the most conservative (Supplementary S1).

### M5. Tests of selection, their reproducibility, and the null distributions

Selection regimes were tested with RELAX (Wertheim et al., 2015) in HyPhy 2.5.100, MP build (Kosakovsky Pond et al., 2020); RELAX.bf carries its own version string, 4.7. Terminal branches leading to ECM species were labelled {Test} (the foreground, in branch-model terms), leaving non-ECM tips and all internal branches as the reference set (the background). *K* and its likelihood-ratio *P* came from the RELAX JSON, *K* < 1 denoting relaxed and *K* > 1 intensified selection on the test branches; rejections are counted two-sided. Runs used --starting-points 5 unless stated otherwise, with --srv, --multiple-hits and --error-sink off at their defaults, the usage characterised here. Command lines and seeds are in Supplementary S3.

### Reproducibility and convergence diagnostics

We use *arm* throughout for one complete configuration of program, model, flag settings, labelling scheme and starting-point budget, run over a whole gene set. Each arm is read against its own baseline; where two arms are compared directly, the comparison is made on the genes interpretable in every arm compared, with that common set stated. A fit is *interpretable* when it returns a fitted *K* and a non-negative likelihood-ratio statistic; a fit terminating with a negative statistic after the five automatic restarts RELAX applies is assigned *P* = 1, cannot reject, and is excluded from any rate quoted on interpretable fits. Zero-byte outputs count as failures, and every paired comparison uses the intersection of genes succeeding in both arms, with its size reported.

### Null distributions

Three null sets were built from the same 51 genomes with coding sequences, so null and test share the alignment pipeline. Two are described below; the third, an earlier random labelling drawn from the whole panel rather than from the non-ectomycorrhizal species and fitted from one starting point rather than five, is null A, reported in Supplementary S5.8 only as a check.

**The selection analyses use the lifestyle assignment as it stood before the audit of M6**, under which *Morchella importuna*, *Penicillium swiecickii* and *Kurtia argillacea* are ectomycorrhizal, so the test set is 21 tips rather than 18; the count analyses use the corrected assignment. The nulls were built under the earlier assignment, and the difference is conservative (S3).

#### Null B, ECM labelling on single-copy orthologues

Every single-copy fungi_odb10 BUSCO gene (Manni et al., 2021) in all 51 CDS genomes went through the M4 alignment pipeline with the same 21 ECM tips labelled {Test}. Eighty-nine met the presence criterion, one (186574at4751) failing tree inference on a non-rectangular alignment; of 88 submitted, 82 completed, 70 of them with an interpretable statistic as defined above. No gene was dropped on the basis of its result.

#### Null C, ECM-free random labelling

The same alignments and gene trees were re-analysed with only labels changed, 21 tips per gene being drawn at random from the 30 non-ECM genomes, the deposited draws rather than the seed defining the analysis (S3). All 88 draws contained zero ECM tips and exactly 21 test tips; 81 completed, a different final *n* on the same alignments. A retired earlier version drew from all 51 genomes (S3).

Empirical *P* for relaxation was computed against each null as (number of null genes with *K* ≤ *K*_observed_, plus one) / (*n*_null_ + 1); with *n*_null_ between 75 and 82 the floor lies between 0.012 and 0.013, so across the 27 tested entries no family can pass Bonferroni correction, reversing version 1 of this preprint.

Tests of a proportion against a fixed value are exact binomial; interval estimates of a proportion are Wilson score intervals, except in the two registered panels, where the registered rule specifies exact (Clopper–Pearson) intervals (S3).

#### Dependence on test-set size

The nulls label 21 of 99 branches (fraction 0.212), whereas tested families span 0.047–0.255. On the same genes, alignments and trees, a *test-fraction series* of eight *steps* changed only the number of labelled tips, from 3 to 29 (fractions 0.030 to 0.293), drawn from non-ECM tips; 69 genes returned results at every step, and the fifth percentile of null *K* per step was bootstrapped over genes (S3).

#### Dependence on gene set

A further control differs from null C in gene set alone: forty non-plant-cell-wall-degrading CAZyme families, at alignment depths of 49 to 342, processed as null C and labelled as the real analysis, median test fraction 0.215 (S3).

#### A second implementation

To separate the test from HyPhy’s implementation, the question was repeated in codeml (PAML 4.10.10; Yang, 2007), fitting one-ratio against two-ratio with the labelled branches as foreground, likelihood-ratio test on one degree of freedom, each model fitted from *ω* = 0.5 and 1.5 with the higher likelihood retained. Four sets mirror the RELAX runs. These are null C (88 genes), the *K* = 1 replicates of the first (shallow, unscaled) simulated set labelled on tips (100) and the same alignments labelled on internal branches (100), and the divergence-matched *K* = 1 replicates labelled on tips (80). The matched set was fitted under terminal labelling only and so has no internal-branch counterpart in this program (S3).

#### A second model family within that program

Because the two-ratio model assumes one *ω* across sites, those sets were analysed again under Clade Model C against its null M2a_rel (Weadick and Chang, 2012), which it exceeds by exactly one parameter. An initial pass truncated the slowest fits at rates differing between observed and simulated sets, so all were re-run to completion; of the divergence-matched set’s 320 clade-model fits, two remained incomplete (S3).

##### A third implementation, asking the question site by site

Null C was analysed with Contrast-FEL (Kosakovsky Pond et al., 2021), which compares *ω* between branch sets site by site with no *K* parameter, at its defaults in HyPhy 2.5.100, which leave synonymous rate variation on (S3).

### Model-misspecification controls

Three misspecification mechanisms (unmodelled synonymous rate variation, Wisotsky et al., 2020; instantaneous multinucleotide substitution, Venkat et al., 2018; Lucaci et al., 2021; and alignment error absorbed by a rate class) were controlled by re-running null C with --srv Yes, --multiple-hits Double or --error-sink Yes, one at a time, each reported on the paired intersection (S3).

### Simulation under a known generating value

Alignments were simulated under the fitted model and re-analysed with the identical invocation. Each template gene supplied its own parameters, taken from its own RELAX null fit and its own alignment and {Test}-labelled tree, and sequences were evolved with pyvolve 1.1.0 (Spielman and Wilke, 2015) under MG94 (Muse and Gaut, 1994), the reference partition at the fitted *ω* classes and the test partition at *ω^K^*. Five template genes were simulated at *K* ∈ {0.5, 0.7, 1.0}. No random seed was passed to the simulator for this first set, so its replicates cannot be regenerated and are deposited individually (S3).

A second, divergence-matched set was built because the first was far shallower than the alignments it stood for. The first set took its branch lengths from the FastTree gene trees, which are in expected substitutions per nucleotide site, and the simulator applies a branch length per codon; the rescaling below corrects this along with any other difference in depth, because it is set by the realised identity rather than by the units. Branch-length scale was set per template by log-scale bisection until the simulated alignment matched that template’s observed mean pairwise identity, separately at each generating *K*; one template could not reach its target and was dropped. Only the scale is altered, so the generating *K* is preserved.

A third set extended the simulation to the null C gene set. Every gene with an interpretable RELAX null fit was classified by that fit, before simulation, into a *moderate* stratum (57 genes: all *ω* classes below 100, one above one), *no fitted heterogeneity* (seven) and a *boundary* stratum (17: a class at or above 100); three boundary genes at the optimiser ceiling were excluded, leaving 78. Four replicates per gene were generated at *K* = 1 under recorded seeds, with the generator of the first set and so at its shallow depth, and fitted with the two-ratio model under terminal labels and under internal labels on the 21 longest internal branches. The strata and the primary summary were fixed in a dated plan; the paired McNemar test was not (S3).

Every simulated alignment was fitted at --starting-points 1 and --starting-points 5 into separate output sets, the five-point arm being the one compared against the observed nulls. A third arm labelled internal branches rather than tips, reusing those alignments, which at *K* = 1 are independent of the labelling (S3).

### Two panels registered in advance

The predictions, decision rules, selection rules and analysis script for two further panels were fixed and their checksums recorded before any fit was run, and registered with the Open Science Framework (https://osf.io/cez4f) at 15:14 UTC on 30 September 2026, under embargo until 1 March 2027. The registered text is in Korean; an English record of every amendment and result is deposited with the data, and the rules are given in full in S3.

In each panel, 50 species (yeasts of the order Pichiales with the standard nuclear code, after Opulente et al., 2024, and the Codetta method of Shulgina and Eddy, 2021; lactobacilli after Duar et al., 2017, on the GTDB tree of Parks et al., 2026), 80 BUSCO genes (Manni et al., 2021) present in single copy in all 50 were sampled at random, aligned and treed as in M4, and given their own random terminal labelling of *k* non-trait tips, *k* being the number of trait species (19 and 18). Fits followed the fungal panel exactly. Replicates were simulated from each gene’s RELAX null fit as in the third simulated set: eight per gene at *K* = 1 for the comparison of labellings, 100 per gene and labelling for up to 30 genes as gene-specific references, 60 per gene as each real gene’s parametric bootstrap reference, and four per gene at *K* = 0.5 and 0.7 as positive controls. The four predictions were judged by an exact binomial test, an exact McNemar test, a cluster-bootstrap bound and an exact interval respectively (Results 2, Results 3 and Results 5); fits unfinished within 24 hours were counted as incomplete, not retried. Four sets of amendments were made after the lock, each before the fits it affects (S3). The registration specified that a failure of the fourth prediction would narrow the title towards an empirical null built from real genes; we did not, since that null has not been tested on real data either.

#### Exploratory follow-up

Five analyses were added after the fourth prediction had failed. Each plan, with its scripts, was fixed by checksum before any statistic it reports existed, with one exception stated in S3, and none changes a registered verdict. They were, in order: a parametric bootstrap from each gene’s one-ratio codeml fit, simulated with PAML 4.10.10 evolver, which reads branch lengths in codeml’s unit, a second generator named in the plan having been dropped before the full run began; codeml’s branch-site model A (Zhang et al., 2005) on the same alignments and labelled trees, from two starting values per model as planned and, after that result was known and because the alternative had ended below the null on 18 genes, from three more per model, one started from the other model’s optimum; the real gap pattern imposed on the replicates of the first analysis for the ten gappiest genes per panel, planned after both results were known; planned after the third result, replicates of 20 genes per panel simulated with pyvolve (Spielman and Wilke, 2015) under each gene’s per-branch *ω* pattern from MG94xREV fits in HyPhy 2.5.100 (Kosakovsky Pond et al., 2020) and, as a control carrying only estimation noise, under the same fit to a one-ratio replicate, each refitted with fresh random labels (S3); and, planned after every branch-site result was known, the power of the branch-site test on alignments simulated with pyvolve from the fitted branch-site null of ten genes per panel, with positive selection added on every labelled branch (S7.4). A worked example on a published RELAX study, planned and fixed by checksum before any run whose result is read, is described in Supplementary S8.8.

#### The labelling design on real and depth-matched data

Two further exploratory analyses, each planned and fixed by checksum before any of its fits existed, address the labelling design directly. In the first, on the replicates of the first follow-up analysis (20 per gene in both new panels), the label was placed on the random terminal set, on the longest terminal branches, on the longest internal branches, and on a set of internal branches whose summed generating length matched that of the random terminal set within 5%, each fitted with the two-ratio model and summarised with gene-clustered bootstrap intervals (S7.6). In the second, on the real null genes of all three panels, branches were divided into the randomly labelled terminals, the other reference terminals, the trait terminals and the internal branches; with the trait terminals and the internal branches given their own *ω*, a model giving the two reference classes separate *ω* was tested against one in which they share it, by a likelihood-ratio test on one degree of freedom, each model fitted from *ω* = 0.5 and 1.5 (S7.7).

### M6. Phylogenetic comparative analysis of gene-family reduction

Per-genome PCWDE counts were analysed against the 182-tip OrthoFinder species tree (Emms and Kelly, 2019) by phylogenetic generalised least squares (PGLS), fitted with caper::pgls (version 1.0.4; Freckleton et al., 2002; Orme et al., 2018) in R 4.5.3 over trees handled with ape (Paradis and Schliep, 2019), as log10(count + 1) ∼ is_ECM with Pagel’s λ by maximum likelihood (Pagel, 1999) and an ordinary least-squares baseline. Fold changes are the back-transformed slope, 10^β^, with normal-approximation intervals 10^β^ ^± 1.96^ ^SE^. The primary contrast is ECM versus free-living decomposers (SAP + WD), ECM versus all non-ECM genomes the secondary. The primary contrast was also referred to 1,000 relabellings of the same tree with λ held at its fitted value, drawn uniformly and by labelling the highest tips of a Brownian trait. A specificity control and a three-step decomposition separating composition from catalytic integrity are described in S3.

#### Lifestyle assignment and its audit

Lifestyle came from the ecology field of JGI MycoCosm metadata, not the genus-level FungalTraits assignment the preprint described. An audit against the primary literature (S3) corrected three. Morco1 (*Morchella importuna*) and Penswi1 (*Penicillium swiecickii*) were reassigned to saprotroph, and Kurarg1 (*Kurtia argillacea*) to ericoid mycorrhizal. The corrected assignment is primary, the uncorrected a sensitivity scenario. The selection analyses retain the pre-audit assignment, for the reason given in M5.

#### Optimiser handling

Two scenarios that failed under maximum-likelihood estimation of λ were recovered by profiling λ over a fixed grid, and refitting with BLAS threads pinned to one reproduced these fits (S3).

#### Data-availability sensitivity

Because 73 panel genomes are hosted on MycoCosm without an associated publication and may fall under the JGI data-use policy, the analysis was repeated on the published subset and both fits are reported. Under the corrected assignment the published subset retains 58 of the 119 ectomycorrhizal genomes and 34 of the 35 free-living decomposers.

## RESULTS 1 — THE PANEL AND THE COUNT

### The panel

Of 196 genomes identified from the mycorrhizal and comparison sets, 186 were retrievable; 183 met the 85% BUSCO completeness threshold, two *Suillus* assemblies fell below it, and one genome yielded no proteome. The 183 carry copy-number annotations, 182 are placed in the species tree and enter every phylogenetically corrected analysis, and codon alignments were recoverable for 51 species, 21 of them ectomycorrhizal under the assignment the selection analyses retain (18 after the audit below), a subset that is not a random sample of the 182 (Discussion 4). Every count below states which panel it refers to (Supplementary S1).

Three of the 122 ectomycorrhizal assignments in the 182-genome phylogenetic panel are corrected below, giving 119 ectomycorrhizal against 35 free-living decomposers in the primary contrast. Table 1 gives the composition of every guild under the corrected assignment, which is also the assignment all three panels of Figure 1 are drawn under; mean copy numbers per guild, a raw comparison on the 183-genome panel and the definition of the family set are in Supplementary S2.7.

**Figure 1.**
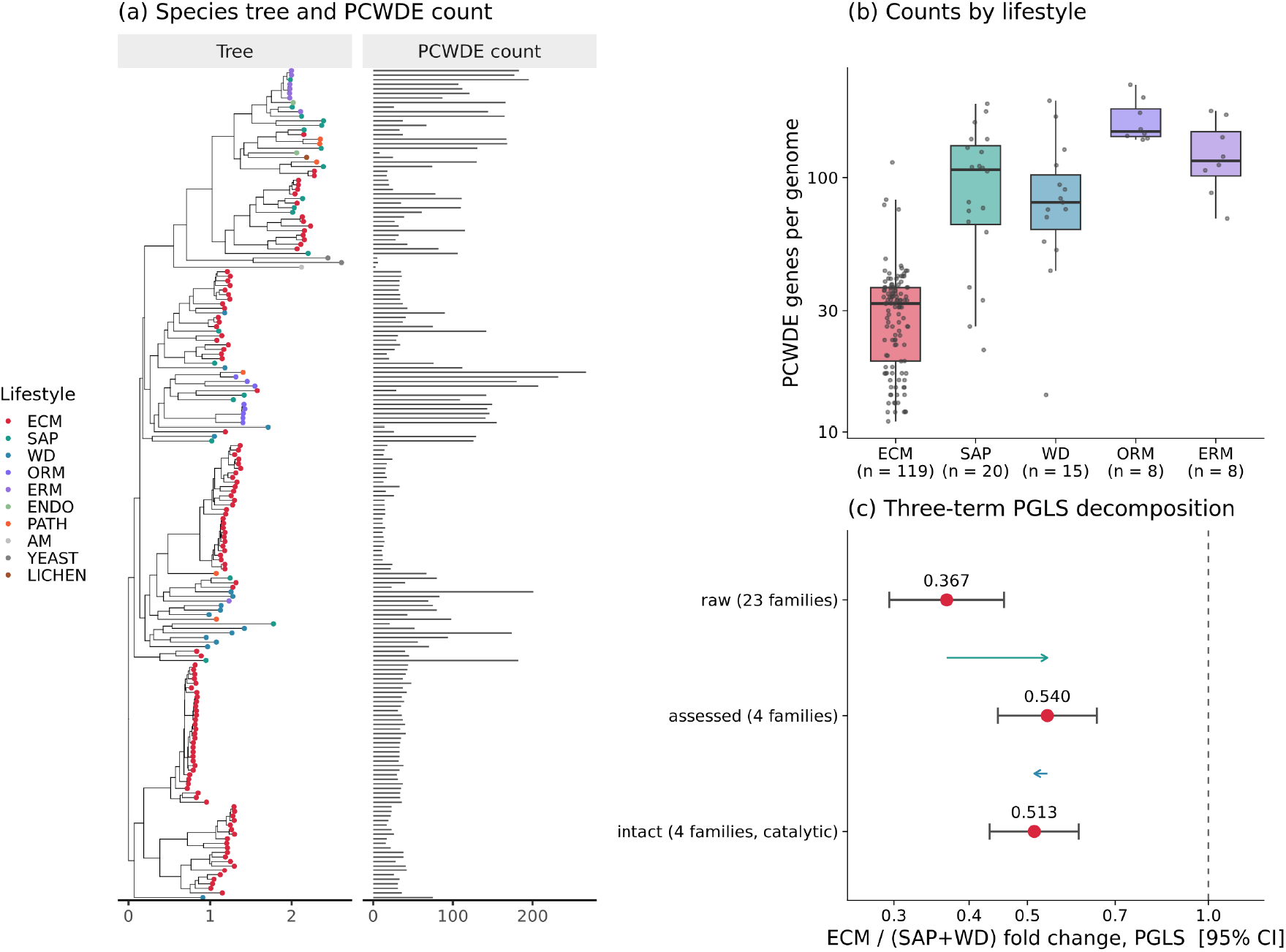
Gene content across the genome panel, and the three-step PGLS decomposition. All panels show observed genomes, not simulated data, and all are drawn under the corrected lifestyle assignment of Results 1 (ectomycorrhizal 119, saprotroph 20, ericoid 8), not the assignment before that audit. (a) The 182-tip species tree, tips coloured by lifestyle, with a bar per genome giving its total plant-cell-wall-degrading enzyme (PCWDE) count across the 23 families. The tree’s scale axis is in amino-acid substitutions per site; the adjacent axis is a gene count. (b) PCWDE genes per genome for the five lifestyles represented by eight or more genomes, log scale; boxes are median and interquartile range, points single genomes; *n* on the axis (ECM 119, SAP 20, WD 15, ORM 8, ERM 8). The remaining twelve genomes of the 182 (six pathogens, two endophytes, two yeasts, one lichenised and one arbuscular mycorrhizal) are omitted as too few to plot as distributions and are tabulated in Table 1, where the pathogen mean of 149.5 is the second highest of any guild. (c) Three phylogenetic generalised least-squares fits over the same tips, each plotted as the ectomycorrhizal (119) to free-living decomposer (saprotrophs plus wood decayers, 35) fold change with 95% confidence interval, on a log axis; the dashed line marks 1.0, no difference. Raw counts across 23 families, 0.367 (0.295–0.457, *P* = 1.3 × 10^-15^); all copies in the four assessed families, 0.540 (0.446–0.652, *P* = 2.1 × 10^-9^); catalytically intact copies within those same four families, 0.513 (0.433–0.609, *P* = 1.9 × 10^-12^). Arrows connect successive fits: the first step narrows the family set from 23 to four, the second restricts those four families to catalytically intact copies and moves the estimate in the opposite direction.

**Table 1.** Plant-cell-wall-degrading enzyme copy number by lifestyle across the 182-tip phylogenetic panel. Counts are summed over the twenty-three CAZy families designated as plant-cell-wall degrading (Results 1), one row per lifestyle, after the three assignments corrected in Results 1. Means are unweighted across genomes and are not phylogenetically corrected; the corrected contrast is in Results 1. The ordering does not follow a symbiotic against free-living division.

| **Lifestyle** | ***n*** | **Mean** | **Median** | **Range** |
|---|---|---|---|---|
| Orchid mycorrhizal | 8 | 169.1 | 152.0 | 141–232 |
| Pathogen | 6 | 149.5 | 148.5 | 67–267 |
| Ericoid mycorrhizal | 8 | 125.0 | 116.5 | 69–183 |
| Saprotroph | 20 | 99.7 | 107.5 | 21–195 |
| Wood decayer | 15 | 89.9 | 80.0 | 14–201 |
| Endophyte | 2 | 87.0 | 87.0 | 8–166 |
| **Ectomycorrhizal** | **119** | **30.3** | **32.0** | **11–115** |
| Lichenised | 1 | 25.0 | 25.0 | 25 |
| Yeast | 2 | 5.5 | 5.5 | 5–6 |
| Arbuscular mycorrhizal | 1 | 3.0 | 3.0 | 3 |

The arbuscular and lichenised rows rest on one genome each and the yeast and endophyte rows on two; the orchid and ericoid means are discussed in Supplementary S2.7.

### The count

Copy numbers for the twenty-three plant-cell-wall-degrading enzyme families were regressed on lifestyle across the 182-tip species tree by phylogenetic generalised least squares with Pagel’s λ estimated by maximum likelihood, setting ectomycorrhizal genomes (*n* = 119) against free-living decomposers, saprotrophs and wood decayers combined (*n* = 35).

Three genomes were reassigned before fitting. An audit against the primary literature of all 122 genomes originally labelled ectomycorrhizal found three that are not: *Morchella importuna*, *Penicillium swiecickii* and *Kurtia argillacea* (Methods M6); the reassignment is reported as a correction and the uncorrected assignment retained as a sensitivity scenario (0.410, below).

Ectomycorrhizal genomes carry 0.367 times as many of these genes as free-living decomposers (Figure 1c; 95% normal-approximation confidence interval on the log scale, 0.295–0.457; *t* = -8.94 on 152 degrees of freedom, *P* = 1.3 × 10^-15^; λ = 0.921). Version 1 of this preprint (Kim and Shin, 2026a), before the three reassignments, gave 0.410 (0.323–0.521; *P* = 1.6 × 10^-11^), which our re-run reproduces exactly; the corrected estimate is both further from unity and better supported. Restricting the comparison to public genomes, which halves the ectomycorrhizal sample to 58 against 34 comparators, gives 0.411 (0.316–0.535; *P* = 2.3 × 10^-9^).

Ectomycorrhizal genomes also carry 0.774 times as many of the non-plant-cell-wall carbohydrate-active families (0.682–0.877), and entering that complement as a covariate moves the headline estimate from 0.367 to 0.501, so roughly a third of the effect on the log scale is a general reduction. The within-genome ratio of plant-cell-wall to other carbohydrate-active families, which is free of that general reduction, is 0.473 times as large in ectomycorrhizal genomes (0.402–0.556; *P* = 4.4 × 10^-16^). **The claim the counts support is the ratio, not the raw depletion.** The headline estimate is unchanged by entering BUSCO completeness or log_10_ assembly N50 as a covariate, moves only away from unity when genomes of ambiguous lifestyle are excluded, and does not rest on λ (Supplementary S2).

An earlier version of this work (Kim and Shin, 2026a) attributed the gap between a 0.41-fold change in raw copy number and a 0.55-fold change in catalytically intact copies to intactness. Decomposing the comparison into three steps shows that the gap comes from the change from twenty-three families to four, and we withdraw the implication that surviving copies in ectomycorrhizal genomes are better preserved catalytically (Supplementary S2.8, S2.5).

### The count verdict against a null generated under its own labelling design

The prescription this paper ends with, that a statistic be referred to a distribution generated under the design that produced it, applies to the verdict we keep as much as to the one we withdraw. The ectomycorrhizal labelling is strongly clustered, the 119 labelled tips falling into eighteen wholly ectomycorrhizal clades, the largest holding 51 (Supplementary S4.5), so the 152 degrees of freedom the model reports are not 152 independent comparisons.

We therefore refitted the primary contrast under 1,000 relabellings of the same tree with λ at its fitted value, under two nulls: 119 tips drawn uniformly, and the 119 highest tips of a Brownian trait simulated on the tree, in the manner of a binary permulation (Saputra et al., 2021), which reaches the observed clustering and passes through it (Supplementary S2.2). The most extreme *t* statistic among the 2,000 relabellings was -3.72 against an observed -8.94, so the empirical *P* is at the floor of 1/1,001 under both nulls. The count verdict survives the test the selection verdict fails, against a null whose clustering spans the observed value; the claim rests on this comparison rather than on the model’s *t*-test.

## RESULTS 2 — RANDOMLY PLACED LABELS ARE REJECTED ON REAL GENES

### The fungal panel

Single-copy orthologues have no reason to be under relaxed selection specifically in ectomycorrhizal lineages, and run through the identical pipeline they measure how often the test rejects a presumed null in this design (Figure 2). On the 51-species coding subset, 88 such genes were labelled twice: on the 21 tips labelled ectomycorrhizal when the codon analyses were fixed (null B; Methods M5), and on 21 tips drawn at random, gene by gene, from the 30 non-ectomycorrhizal species (null C; a third, earlier labelling, null A, is reported in Supplementary S5.8 only as a check). RELAX rejected 51.2% of genes under the first labelling (42 of 82) and 35.8% under the second (29 of 81), about ten and seven times the nominal 5%. On the random labels the two-ratio branch model of codeml rejected 58 of 88 genes (65.9%), and Clade Model C, which lets one of three site classes differ between branch sets (Weadick and Chang, 2012), 40 of 88 (45.5%); Contrast-FEL (Kosakovsky Pond et al., 2021), which compares the branch sets site by site, rejected at 9.18% of 61,059 sites at its asymptotic stage. Whole-set rates keep genes without an interpretable statistic in the denominator as non-rejections (Methods M5); interpretable-only rates, the rates on the genes common to all three branch-level tests and the gene-level overlap between programs are in Supplementary S5.8 and S5.3.

**Figure 2.**
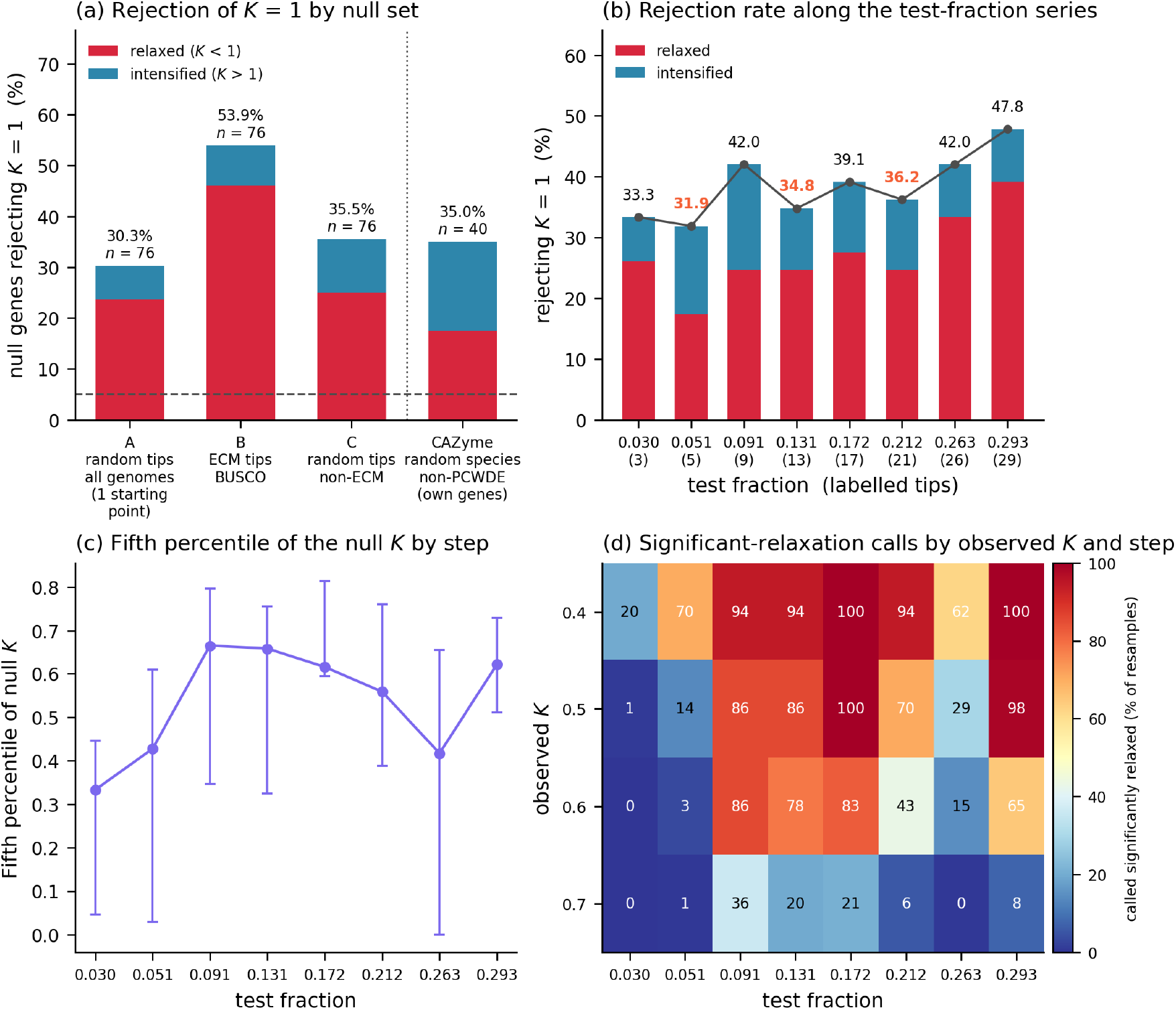
Every null set rejects *K* = 1 far above the nominal 5% (dashed line), and the rate depends on the labelling design. All panels use RELAX with terminal (tip) labelling; bars split rejections into relaxed (*K* < 1) and intensified (*K* > 1). (a) Percentage of null genes rejecting *K* = 1 in four sets: A, random tips from all genomes at one optimiser starting point; B, the pre-audit ectomycorrhizal tips on single-copy BUSCO orthologues (three of the twenty-one were later reassigned; Methods M5); C, random tips from non-ectomycorrhizal species, B and C at five starting points. A, B and C are counted on a common 76 genes interpretable in all three: 30.3, 53.9 and 35.5%. The CAZyme control, right of the dotted rule, is 40 non-plant-cell-wall-degrading families on their own genes and cannot be paired with them: 14 of 40, 35.0%. (b) Rejection rate along the test-fraction series, set C re-analysed changing only the number of labelled tips (3, 5, 9, 13, 17, 21, 26, 29), on the 69 genes interpretable at all eight steps: 31.9% at five tips to 47.8% at 29. The rank correlation with step size and its exact two-sided permutation *P* are ρ = 0.77 and *P* = 0.033; the three rates picked out in orange are the steps that fall below their smaller neighbour, at five, thirteen and twenty-one tips. (c) Fifth percentile of the null *K* distribution per step, the relaxed-side threshold, same 69 genes, spanning 0.33 to 0.66. Bars are bootstrap intervals over 5,000 gene resamples. In none of those resamples did the eight percentiles emerge in step order; we report that and set it aside, since the statistic resamples around an already-disordered point estimate and would return zero under a monotone truth as well (Supplementary S5.4). (d) For each of four observed values of *K*, the percentage of those 5,000 resamples in which the observation falls below the step’s fifth percentile and is therefore called a significant relaxation; steps as in (b) and (c). At *K* = 0.5 it runs from 1% at three labelled tips to 100% at seventeen, so the same observation is called significant essentially never or essentially always depending on which step supplies the null. The dependence is not confined to that value: the range across steps is 20 to 100% at *K* = 0.4, 0 to 86% at *K* = 0.6 and 0 to 36% at *K* = 0.7.

The ectomycorrhizal labelling was rejected more often than the random one on the 76 genes scored under both (McNemar exact *P* = 0.044), but the difference is not established as ectomycorrhizal-specific: three of its 21 tips are not ectomycorrhizal, and the random labelling is not structure-preserving (Saputra et al., 2021), though clustering does not predict rejection among the draws (Supplementary S5.1, S5.8).

Switching on, one at a time, the three RELAX options that change the fitted model (unmodelled synonymous rate variation, Wisotsky et al., 2020; instantaneous multinucleotide substitution, Venkat et al., 2018; Lucaci et al., 2021; and a rate class intended to absorb alignment error) did not return null C to the nominal rate: 35.2 to 32.4% (*n* = 71), 41.3 to 34.9% (*n* = 63) and 35.1 to 51.4% (*n* = 74) respectively, each against its own baseline on the genes that succeeded under both. Synonymous rate variation, the mechanism one would expect to account for an inflation of this size, moves the rate by under three percentage points (Supplementary S5.2).

### Two further panels, registered in advance

Every result so far comes from one panel of 51 fungi. Before fitting a single gene in two further panels we registered four predictions and their decision rules (Methods M5; https://osf.io/cez4f): 50 yeasts of the order Pichiales, where the trait is failure to grow on glycerol (Opulente et al., 2024), and 50 lactobacilli, where it is host adaptation against free-living and nomadic lifestyles (Duar et al., 2017), each with 80 single-copy null genes. The primary labelling in both is not the trait but a set of terminal labels drawn at random, gene by gene, from the non-trait species, 19 tips in yeast and 18 in lactobacilli, so that a genome-wide shift in selection on the trait lineages cannot masquerade as a false positive. All nine verdicts (the four predictions in each panel, and the third also in the fungal panel) are reported without correction for multiplicity, every count was reproduced by a second, independently written parser, and the amendments made after the lock are listed in Supplementary S3. Both panels are about half as deep as the fungal one, with a median total tree length under a one-ratio fit of 52.5 in yeast and 61.4 in lactobacilli against 116.3 (Supplementary S7.1).

The two-ratio test rejected the null at χ^2^_1_ on 55 of the 80 yeast genes, 68.8% (one-sided exact binomial *P* = 2.9 × 10^-52^), and on 71 of the 80 lactobacillus genes, 88.8% (95% Clopper–Pearson CI 79.7–94.7; *P* = 6.2 × 10^-82^), against 65.9% for the comparable null on the fungal panel (Figure 3a). The first prediction holds in both. The trait labels themselves, reported but not used for the verdict, were rejected less often than the random ones in lactobacilli, 33.8 against 88.8%, and only slightly less in yeast, 60.0% against 68.8%.

**Figure 3.**
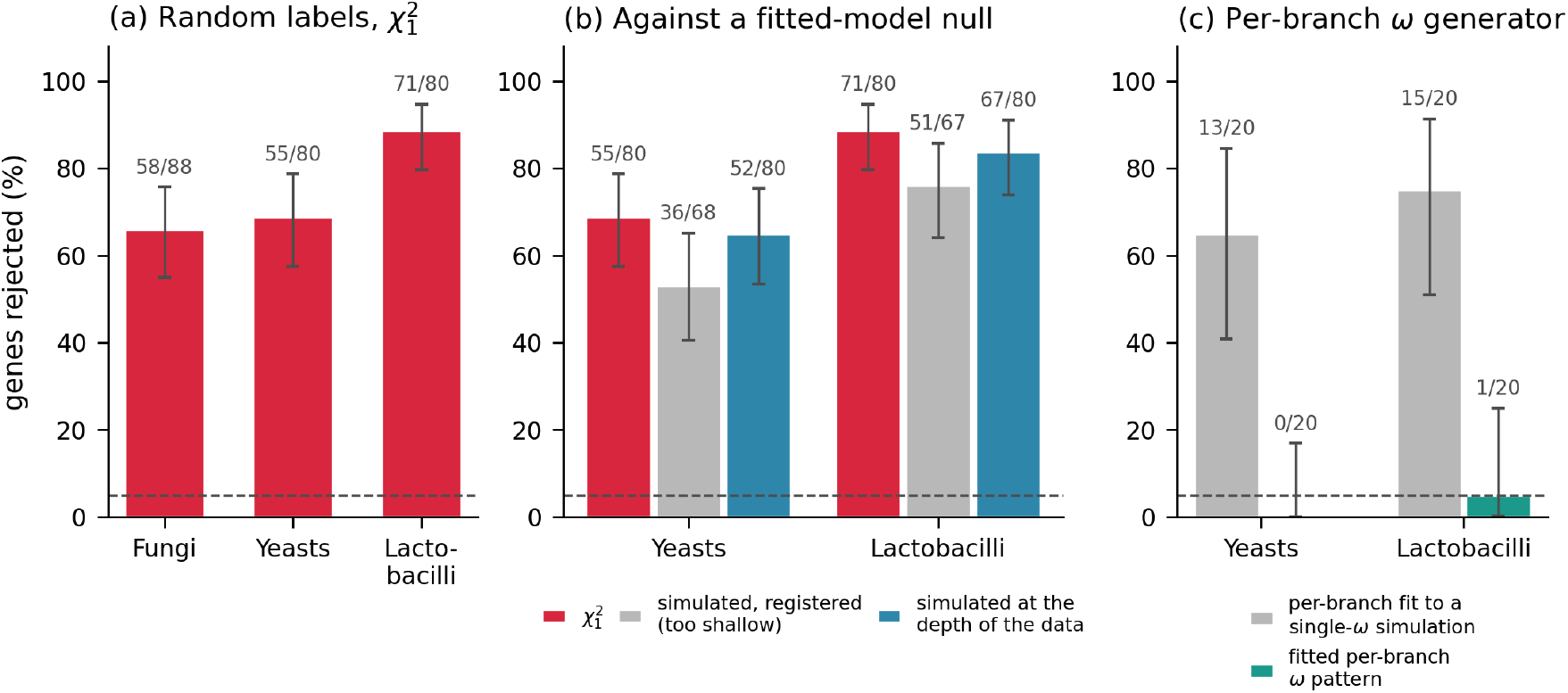
Randomly placed labels on real genes. Bars are the percentage of null genes rejected, with 95% Clopper–Pearson intervals; the numbers above the bars are genes rejected over genes scored. (a) The two-ratio branch model at χ^2^_1_ under terminal labels drawn at random from the non-trait species: 21 tips in the fungal panel (null C; 58 of 88 genes, 65.9%), 19 in yeasts (55 of 80, 68.8%) and 18 in lactobacilli (71 of 80, 88.8%). Dashed line, nominal 5%. (b) The yeast and lactobacillus genes of (a) referred instead to 60 replicates simulated from each gene’s own fitted null model: with the registered generator, whose replicates were far shallower than the data (36 of 68 and 51 of 67 genes), and with a generator at the depth of the data (52 of 80, 65.0%, and 67 of 80, 83.8%); red bars repeat the χ^2^_1_ rates of (a) (Results 3). Dashed line, the exact size of the reference, 3/61 or 4.9%. (c) Twenty genes per panel referred to 40 replicates simulated under a model with one *ω* per branch and refitted with fresh random labels: replicates built from a per-branch fit to a single-*ω* simulation, which carry the estimation noise of such a fit and no true variation (13 and 15 of 20 genes), and replicates built from each gene’s own fitted per-branch pattern (0 and 1 of 20). The fitted pattern is an upper bound on the variation it represents (Results 4). Dashed line, nominal 5%.

**On real genes, a randomly chosen set of terminal branches is therefore rejected by the two-ratio test in 65.9 to 88.8% of genes, in three panels and under a labelling that carries no trait.** The χ^2^ reference cannot be used at face value in this design, and a trait label cannot be read against it.

## RESULTS 3 — A NULL SIMULATED FROM THE FITTED MODEL DOES NOT REMOVE THE EXCESS

The standard remedy for a miscalibrated likelihood-ratio test is a parametric bootstrap: the statistic is referred to replicates simulated under the fitted null model on the same labelled tree, as Halabi et al. (2021) do for a test of this family. On simulated data such a reference returns the nominal rate by construction, since a replicate referred to others drawn from the same generator falls in their upper 5% about one time in twenty whatever the test does (Results 5); whether it calibrates a real alignment, which was not drawn from that model, simulation cannot show. The fourth registered prediction asked that question of the two new panels.

### As registered

The fourth prediction tests the remedy itself on data it was not drawn from, and it fails. Each real null gene, with its random terminal labels, was referred to 60 replicates simulated from its own fitted model on the same labelled tree, a reference whose exact size is 3/61, or 4.9%. In yeast 36 of 68 genes were still rejected, 52.9% (95% CI 40.4–65.2), against 69.1% at χ^2^_1_ on the same genes; in lactobacilli 51 of 67 were, 76.1% (64.1–85.7), against 86.6%. Neither interval contains 5%, and the conclusion is unchanged when the genes without a generator are counted as not rejected (Supplementary S7.3).

The replicates were also far shallower than the data. Fitted with the same one-ratio model, the simulated alignments have a median total tree length 0.16 times that of the real alignment in yeast and 0.13 times in lactobacilli, and the likelihood-ratio statistic is correspondingly small, with a median of 1.4 on the simulated replicates against 13.1 on the real genes in yeast and 1.5 against 29.6 in lactobacilli. The generator took the branch lengths of the nucleotide gene trees and the simulator read each as a length per codon, the defect that made the first simulated set of Results 5 shallow (Methods M5). As registered, the fourth prediction therefore cannot separate a failure of the remedy from a failure of its implementation; the exploratory follow-up below separates them.

### An exploratory follow-up: a generator at the depth of the data, and a branch-site test

Both analyses were designed after the fourth prediction had failed and are not part of the registration; their plans and scripts were fixed by checksum before any statistic they report existed (Methods M5). Two further checks followed them.

The first repeats the fourth prediction with a generator built for the test it calibrates. Each real gene was simulated under its own one-ratio fit, the null model of the very likelihood-ratio test being referred, with PAML’s own simulator, which reads branch lengths in the unit in which they are estimated; sixty replicates per gene were fitted as the real gene had been. The replicates now match the data: their one-ratio tree length is a median 1.05 times the real one in yeast and 1.06 times in lactobacilli, and their fitted *ω* 0.997 and 0.980 times. Every gene had all 60 replicates. Rejection rose rather than fell, to 52 of 80 yeast genes, 65.0% (95% CI 53.5–75.3), and 67 of 80 lactobacillus genes, 83.8% (73.8–91.1), against 55 and 71 of 80 at χ^2^_1_ (Figure 3b); on the 68 and 67 genes scored for the fourth prediction the rates are 64.7 and 82.1%, so the rise is not an effect of the larger denominator. A second planned generator was dropped before the full run began (Supplementary S7.4).

The second asks whether the excess belongs to tests of a difference in *ω* between branch sets or to branch tests in general. On the same genes and random terminal labels, codeml’s branch-site model A, which allows a class of sites with *ω* > 1 on the labelled branches only, rejected 1 of 80 yeast genes and none of 80 lactobacillus genes at χ^2^_1_ with two starting values per model, as planned. Because the alternative model, which contains the null, had ended below it on 14 yeast and 4 lactobacillus genes, every gene was then refitted from three more starting values per model, a change made after the first result was known; the one rejection disappeared, and no gene in either panel was rejected (95% CI 0–4.5% in each). An optimum missed by all five starts cannot be excluded, the test is conservative by design (Zhang et al., 2005), and its power was measured afterwards, on alignments simulated from ten of these genes per panel under each gene’s fitted branch-site null with positive selection added on every labelled branch: with *ω* = 3 or 8 at 10% of sites, or *ω* = 20 at 5%, it rejected 100 of 100 replicates in yeast in each case and 95, 96 and 97 of 100 in lactobacilli, against 5 and 3 of 100 with no effect added (Supplementary S7.4). These alignments come from the fitted model itself, with no other misspecification, so this power is likely optimistic for real data; within that limit the zero on 160 genes bounds the share of these genes carrying an effect of these sizes shared by all the labelled branches at about 2% (one-sided; 3.7 to 4.3% in either panel). Effects confined to some of the labelled branches were not simulated.

Neither generator imposed the gaps and ambiguous characters of the real alignments, which are few (a median of 1.5% of characters in yeast and 0.3% in lactobacilli). In a further exploratory check, the real gap pattern was imposed on the 60 replicates of the ten gappiest genes in each panel and the replicates were refitted; the same 14 of 20 genes were rejected, and no gene changed its decision (Supplementary S7.4).

**Together, a reference simulated at the right depth from the null model of the test itself does not remove the excess, and a test that asks only for positive selection on the labelled branches shows none, although on simulated data it detects positive selection of the sizes we placed on those branches.**

## RESULTS 4 — Ω VARIES AMONG BRANCHES ON REAL GENES

One reading of these results is that *ω* varies among lineages on real genes, so that a randomly chosen set of terminal branches does differ in *ω* from the rest and a test of that difference detects it; no null simulated with a single *ω* for both sets could describe such data. An exploratory analysis, planned after the gap check, tested this reading on 20 of the 80 genes in each panel (Methods M5). A model with one *ω* per branch improved the fit of every real alignment more than it improved the fit of each of five alignments simulated with a single *ω* (40 of 40 genes). Each real gene was then referred to 40 replicates simulated under its own fitted per-branch pattern and refitted with fresh random labels: 1 of 40 genes was rejected (one-sided 95% upper bound 11.3%), against 28 of 40 for replicates built the same way from a per-branch fit to a single-*ω* simulation, which carries the estimation noise of such a fit and no true variation (exact McNemar *P* = 1.1 × 10^-7^; Figure 3c; Supplementary S7.4).

The fitted pattern therefore suffices to account for the excess, but only as an upper bound. It carries estimation noise as well as any true variation; the rule fixed in advance pooled the two panels, and the lactobacillus panel alone would not meet it (1 of 20, bound 21.6%); and a per-branch *ω* cannot separate variation in selection among lineages from other departures it absorbs, such as variation in base composition or synonymous rate. Other properties of real alignments that neither generator reproduces, among them multinucleotide substitutions (Venkat et al., 2018) and alignment error, were not examined. None of these analyses tests the empirical null on real data. Positive controls, reported without a verdict, are in Supplementary S7.5.

In all 40 genes the per-branch fit gives internal branches a higher *ω* than terminal ones, a contrast that by itself sets a random terminal set apart from the rest. We do not identify its cause. Estimated *ω* depends on the span of time a branch covers (Mugal et al., 2014), and errors in the tree or the alignment, or saturation at synonymous sites on long branches, can each place substitutions differently on internal and terminal branches.

Giving those classes their own *ω* does not remove the excess. In an exploratory analysis planned after these results were known (Methods M5), internal branches, the terminals of the trait species and the remaining terminals were each given their own *ω*, and the randomly labelled terminals were tested against the other reference terminals alone. Internal branches had the higher class *ω* in 67 of 88 fungal genes and in all 80 genes of each new panel, and the random terminals were still rejected at χ^2^_1_ in 35 of 88 fungal genes (39.8%; 95% CI 29.5–50.8), 35 of 80 yeast genes (43.8%; 32.7–55.3) and 23 of 80 lactobacillus genes (28.8%; 19.2–40.0), against 65.9, 68.8 and 88.8% under the two-ratio model and a nominal 5% (Supplementary S7.7). The excess therefore persists among reference terminals; its source is not identified here.

## RESULTS 5 — THE LABELLING DESIGN ON DATA WHERE THE ANSWER IS KNOWN

### Simulated data with no difference between branch sets

To see what the test does when the answer is known, alignments were simulated under each template gene’s own fitted RELAX null model on its own labelled gene tree and re-analysed with the identical invocation (Methods M5). Five template genes gave a first set at the depth their gene trees implied, which proved far shallower than the data (median total tree length 12.1 against 116.3 on the null orthologues; the shallow set), the cause being in part a mismatch of units (Methods M5); a second set was rescaled per template until the simulated alignment matched that template’s mean pairwise identity (the divergence-matched set, four templates). A third set took the first generator to the 78 genes of null C with a usable fit, four replicates per gene at *K* = 1. The labelling contrasts below rest on the two-ratio model, whose fits on these data repeat; RELAX rejection rates carry optimiser variance on top of whatever the design contributes (Kim and Shin, 2026b, Results 1 and Results 2). The design and the RELAX results in full are in Supplementary S6.

Under tip labels at *K* = 1, RELAX rejected 22.3% of the shallow replicates (21 of 94; Supplementary Figure S1) and 17.5% of the divergence-matched ones (14 of 80). These alignments contain no alignment error, no rate variation across sites, no multinucleotide substitution and no branch-to-branch heterogeneity, and were generated under the model RELAX fits, so its excess cannot be attributed to any property of real sequence data. The two-ratio model, which assumes one *ω* across sites, rejected 10.0% of the shallow five-template replicates and 46.2% of the matched ones (37 of 80).

### Moving the label from terminal to internal branches

Moving the label changes the rate, and not by the same amount in both programs. On the shallow five-template replicates the two-ratio rate rose from 10.0 to 47.0%, 41 replicates being rejected under internal labels only against four the other way (*P* = 9.3 × 10^-9^), while the RELAX rate fell from 22.5 to 13.5% across the 89 replicates fitted under both. On the divergence-matched set the RELAX rate instead rose, from 17.5 to 35.0% (McNemar exact *P* = 0.016), and the side of *K* = 1 on which its rejections fell reversed, relaxation in 19 of 20 rejections on the shallow set against intensification in 10 of 14 on the matched one (Supplementary S6.6).

These five-template rates do not generalise. Over the 78 genes of null C (312 alignments, each fitted under both labellings) the direction replicated, 98 replicates being rejected under internal labels only against 45 under tip labels only (McNemar exact *P* = 1.1 × 10^-5^; 21.8% under tip labels, 68 of 312, against 38.8%), but the 4.7-fold separation of the five-template rates did not, although the five templates themselves reproduced their earlier behaviour, none of 20 tip-labelled replicates rejecting against 13 of 20 internal. The size of the contrast depended on the fitted *ω* configuration. On the 57 genes whose fitted classes are moderate, the stratum fixed as primary before the batch ran, the rates were 25.4 and 34.2%, 1.3-fold (58 against 38 discordant replicates, *P* = 0.052); on the 14 where a fitted class reaches *ω* ≥ 100 they were 10.7 and 66.1%, 6.2-fold, a stratum that holds every negative likelihood-ratio statistic and starting-point disagreement in either arm (Supplementary S6.10). The tip-labelled rate of 10.0% above is therefore a five-template value. In this set the internal labels were the 21 longest internal branches, so the contrast confounds position in the tree with branch length.

To separate the position of the labelled branches from their length, labels were placed on the replicates of the depth-matched generator of Results 3, simulated under each gene’s one-ratio fit, the null model of the test, so that the test is correctly specified (Methods M5; exploratory, its plan fixed before any of its fits existed). Random terminal labels were rejected in 5.4% of 4,800 replicates in yeast and 6.8% of 4,800 in lactobacilli (60 per gene) (gene-clustered 95% intervals 4.9 to 6.0% and 6.2 to 7.5%, the second reaching the upper limit of the range the plan calls nominal). On the first 20 replicates per gene (1,600 per panel), internal labels whose summed generating length matched that of the random terminal set differed from it by +0.2 percentage points in yeast (90% interval -1.2 to 1.7) and -1.2 in lactobacilli (-2.7 to 0.1), and the 19 or 18 longest internal branches by +0.7 in each (95% intervals -0.9 to 2.3 and -1.2 to 2.6); each interval excludes a difference of 5 percentage points, which on a nominal size of 5% still allows a type I error about twice that size. The longest terminal branches raised the rate by 2.0 percentage points in yeast (95% interval 0.3 to 3.8) and not detectably in lactobacilli (+0.5; -1.2 to 2.2). Because the two panels fall in different rows of the outcome table fixed in the plan, the plan allows no summarising statement, and we make none. The analysis concerns size under correct specification only and cannot apportion the contrasts above, on misspecified simulations or on real genes, between position and length (Supplementary S7.6).

In the two registered panels the second prediction repeated this contrast on alignments simulated at *K* = 1 under each gene’s fitted RELAX null model, eight replicates per gene fitted under both labellings, on the stratum registered as primary, the genes whose fitted *ω* distribution is moderate. These replicates come from the generator of the fourth prediction (Results 3) and so are a sixth to an eighth of the real depth. In yeast, across 416 replicates from 52 genes, 171 were rejected under internal labels only and 14 under terminal labels only (two-sided exact McNemar *P* = 1.7 × 10^-35^), and the gene-averaged rejection rate rose from 25.7 to 63.5%. In lactobacilli, across 328 replicates from 41 genes, the counts were 85 and 22 (*P* = 6.4 × 10^-10^), and the rate rose from 25.6 to 44.8%. The fungal panel, at four replicates per gene, had shown the same direction at 25.4 to 34.2% without reaching the registered level; in both new panels, at twice the replicates per gene, it reached it (Supplementary S7.2).

### A reference built under one labelling does not calibrate another

For 20 genes drawn at random from the moderate stratum, 100 replicates at *K* = 1 were fitted under both labellings, and each internal-labelled replicate was referred to that gene’s own 99 terminal-labelled replicates from the other alignments. Had the design not mattered, about 4% would have been rejected, the exact size of that reference. In the fungal panel 17.0% were (95% gene-level bootstrap CI 9.4–26.0), in yeast 33.8% (25.6–41.6) and in lactobacilli 13.9% (8.6– 19.7); the lower bound exceeds 5% in all three, so the third prediction holds in all three. A reference built under terminal labels is too lenient for an analysis under internal ones; the reverse referral, reported but not predicted, fell close to the nominal size, and a few genes carry most of the mean (Supplementary S7.2).

### The number of labelled branches

The number of labelled tips matters as well. On the fungal null genes and trees, with 3 to 29 labelled tips drawn from the non-ectomycorrhizal species, the RELAX rejection rate rose with the size of the labelled set, from 33.3% at three labelled tips to 47.8% at twenty-nine, with a minimum of 31.9% at five (Figure 2b; Spearman ρ = 0.77; exact two-sided permutation *P* = 0.033; 69 genes at every step), though not monotonically, and the fifth percentile of the null distribution of *K* spanned a twofold range across the steps, so that an observed *K* of 0.5 is called significant essentially always or essentially never depending on the step (Figure 2c, d; Supplementary S5.4). A control of forty enzyme families that are not plant-cell-wall-degrading, at depths of 49 to 342 sequences, agreed with null C but was too small to exclude a gene-set effect as large as this design effect (Supplementary S5.5).

### What simulation can show about the correction

Referring each simulated replicate to the others from the same generator returns the nominal rate by construction, 22.3 to 6.4% under RELAX on the shallow set and 17.5 to 6.2% on the matched set (Figure 4a), so the simulations cannot show that the correction works; they measure what it costs. On the matched set, detection at a true *K* of 0.5 fell from 71.9 to 46.9% and at 0.7 from 65.6 to 9.4% when the bootstrap threshold replaced the χ^2^_1_ threshold, because the ninety-fifth percentile of the null statistic there is 152.6 against a critical value of 3.841; on the shallow set the loss was slight (Figure 4b; Supplementary S5.6).

**Figure 4.**
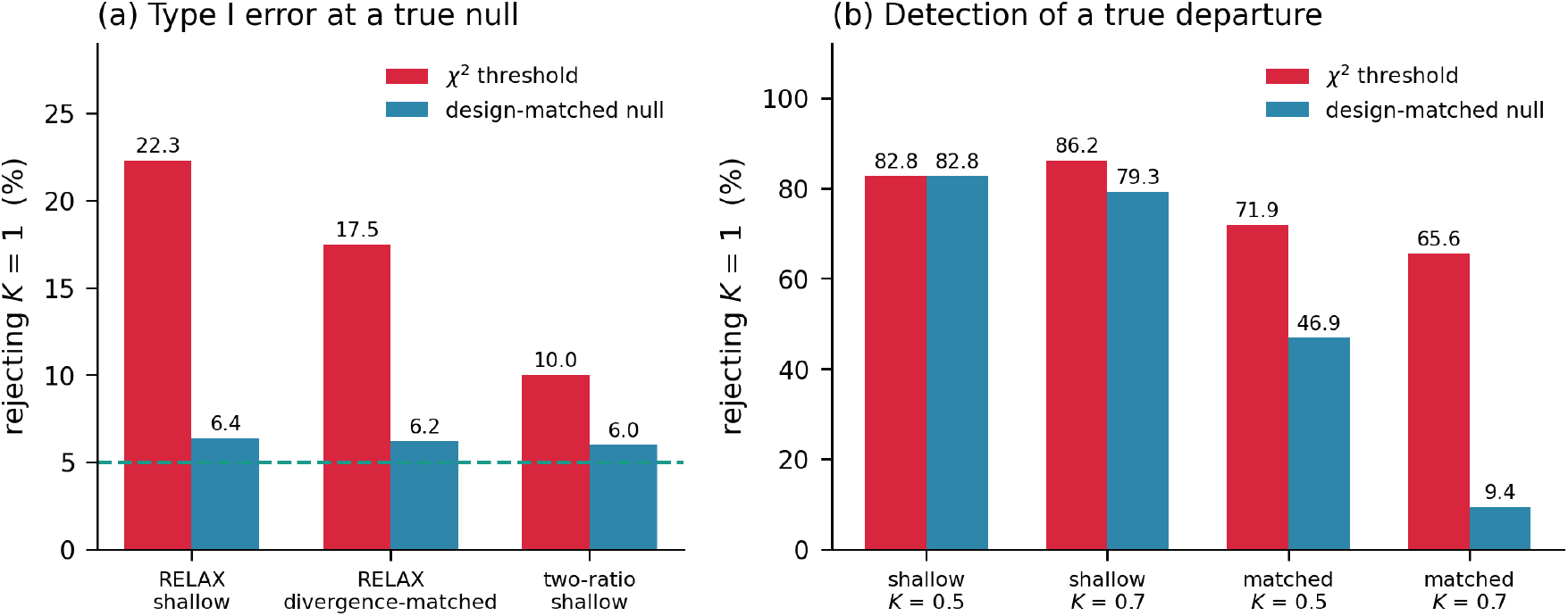
What it costs to refer the likelihood-ratio statistic to a null generated under the same labelling design depends on the divergence of the alignments. Red bars are the asymptotic χ^2^_1_ threshold and blue bars the threshold from the design-matched simulated reference distribution (parametric bootstrap), each replicate assessed against the remaining replicates so that none contributes to its own reference. (a) Percentage of true-null replicates rejecting *K* = 1 in three arms: RELAX on the shallow simulated set (*n* = 94), RELAX on the divergence-matched set (*n* = 80), and the two-ratio branch model on the shallow set (*n* = 100). The substitution takes 22.3 to 6.4%, 17.5 to 6.2% and 10.0 to 6.0%. Because each replicate is referred to others from the same generator, a corrected rate near the nominal 5% (dashed line) follows from the construction; it is shown for scale, not as evidence of calibration. (b) Percentage of replicates carrying a true departure that are detected, under the same substitution. On the shallow set detection is unchanged at *K* = 0.5, 82.8%, and falls from 86.2 to 79.3% at *K* = 0.7 (*n* = 29 at each value); on the divergence-matched set it falls from 71.9 to 46.9% at *K* = 0.5 and from 65.6 to 9.4% at *K* = 0.7 (*n* = 32 at each). The cost is arithmetic rather than a property of the labelling: the ninety-fifth percentile of the null statistic on the matched alignments is 152.6 against a χ^2^_1_ critical value of 3.841 (Results 5, Supplementary S5.6).

**The type I error rate cannot be quoted for the test alone; it depends on the labelling design and on the fitted model**

A study that builds one null and applies it across families of differing test fractions, as ours did, is not applying a calibration but an assumption. We reserve *type I error rate* for the rejection rate a design produces on a true null, and *the correction* for referring the statistic to a design-matched reference distribution.

## RESULTS 6 — THE ENZYME FAMILIES READ AGAINST RANDOM LABELS

Twenty-eight plant-cell-wall-degrading enzyme entries had codon alignments deep enough to test: twenty families, seven GH5 subfamilies and a subsampled GH5. Twenty-seven returned a fitted *K* under both starting-point settings.

Empirical *P* values were computed against the ectomycorrhizal-free null (null C), 81 genes, so no entry could have passed a Bonferroni-corrected threshold whatever the data had been (Discussion 2).

### What the enzyme-family results do not show

Version 1 of this preprint reported relaxed selection in GH6, GH7 and GH28, and argued that gene loss and relaxation were separable axes because GH5 showed no departure at family level. We withdraw all of it, on four grounds. The second and third, the irreproducibility of the fits and the simulation result, each defeat the whole claim, since neither depends on any particular family; the first and fourth say how the error was made.

First, the GH5 family-level result was an optimisation failure, not a signal. The fit in version 1 (Kim and Shin, 2026a) returned *K* = 1.0000 with a likelihood-ratio statistic of zero, and refitting from five starting points moved the log-likelihood by 722 units while returning a significant *P* value in the opposite interpretive direction. The earlier manuscript’s assertion that this was not a power problem was therefore false: it was not a departure-free fit but a fit that had not converged. Second, when the surviving candidates were refitted from more starting points, none of them stayed significant: at ten or twenty starting points each returned a *P* value above 0.05 against the asymptotic (χ^2^) reference distribution it had been read against all along. The last to fall moved by 0.27 in *K* at a cost of 3.3 log-likelihood units, a displacement the likelihood barely distinguishes; its earlier apparent stability rested on agreement between two settings only, and a third disagreed. Third, and independently of both, the simulation in Results 5 rejects the true null at three and a half to four and a half times the nominal rate in this design (17.5% on replicates matched to the divergence of our alignments and 22.3% on a shallower set), so a rejection rate of the size we reported is what this design produces on data containing no signal at all. The argument that the design produces relaxation specifically is withdrawn with it (Kim and Shin, 2026b, Results 1). Fourth, the entries were read against a null that did not match them. Our null carries a test fraction of 0.212 and the entries a median of 0.089, a difference Results 5 shows is not innocuous. Reading each entry against the step of the test-fraction series nearest its own test fraction, the design-matched reading we describe applied to our own data, leaves two of the 27 entries nominally significant rather than three; the entry that drops out is GH7, whose estimate is also the least well identified. The two that remain are GH6 (*P* = 0.049) and GH12 (*P* = 0.037). That pair is not the one the preprint’s three would suggest: refitted from five starting points, GH28 is significant against neither its nearest step nor the single shared null it was originally read against (*P* = 0.256 and 0.232), while GH12 was never among the claims withdrawn here. Neither survivor is rescued by this, since the starting-point refits of the second ground remove both (Supplementary S5.7).

### What survives

One statement survives the withdrawal, and it comes from the count axis rather than the selection axis: ectomycorrhizal genomes carry 0.367 times as many copies of these enzyme families as free-living decomposers, and the within-genome ratio on which the surviving claim rests is reduced 0.473-fold (Results 1).

## DISCUSSION 1 — TWO QUESTIONS, ONE ANSWER

Two questions about the same enzyme families were asked of one body of data, namely the same 183 genomes and annotation, the same 182-tip tree and the same 51-species coding subset. How many copies are there, and what selection are the surviving copies under? One returned an answer and the other did not, and that asymmetry is what this paper sets out to explain.

On the count axis, ectomycorrhizal genomes hold 0.367 times as many of these genes as free-living decomposers (*P* = 1.3 × 10^-15^), an estimate that survives every reassignment of lifestyle we could justify, halving the sample to publicly available genomes, and 2,000 relabellings of its own tree, including a thousand from a permulation-style null whose clustering spans the observed value (Results 1). On the selection axis, the test was answered before the trait was consulted. Randomly placed terminal labels on presumed-null genes were rejected by the two-ratio model in 65.9 to 88.8% of genes in three panels (Results 2); a reference simulated from each gene’s fitted null model did not remove the excess, even at the depth of the data (Results 3); a reference simulated under each gene’s fitted per-branch *ω* did (Results 4). Three properties are easily run together here. The *size* of the test, its rejection rate when the model it fits is the model that generated the data, was close to nominal at the depth of the data, 5.4 and 6.8% (Results 5). *Robustness*, the rate on data generated by a process the fitted model does not describe, moved with where the labels sat and with the fitted model, from 21.8 to 38.8% over 78 gene-fitted models when the labels moved from terminal to internal branches (Results 5). And *trait specificity*, whether a rejection reflects the trait rather than any set of branches, cannot be read from a rejection on real genes, where the null that the labelled branches share *ω* with the rest appears to be false for almost any set; it can be judged only against random labels of the same size. RELAX adds a failure of its own: its fits do not reproduce across optimiser settings, and 46 of 410 terminated with a negative likelihood-ratio statistic after the five automatic restarts it applies (Kim and Shin, 2026b, Results 1 and Results 2). The count is a property of the annotation, recoverable by counting; the selection-intensity parameter is a property of a multiparameter mixture model fitted to a likelihood surface on which relaxation and intensification are separated by a median of 0.95 likelihood units in the 36 of 81 null genes where RELAX’s own flat-surface check fired (Kim and Shin, 2026b, Results 2).

What the two failures of the selection axis share is a form, and it is the form that bears on the asymmetry. In each case the step terminated normally and returned a value of the kind we were looking for, and nothing in its output separated it from a step that had worked; the companion manuscript tabulates the instances. The count can be checked by counting again. The selection intensity cannot be checked by reading the output at all.

We therefore decline to convert our withdrawal into a claim. Our earlier work proposed that gene count and functional capacity come apart in ectomycorrhizal genomes, with copies lost and survivors under maintaining selection. We are not now asserting the opposite. The proposition is not disproved but untested, because the instrument does not distinguish the two answers at this scale. It survives here as a framing question rather than a finding. Discussion 2 sets out what a method must demonstrate about its calibration on branch-partitioned designs before such a claim can be accepted.

## DISCUSSION 2 — WHAT TO DO INSTEAD

The failures documented here arise from a widely used implementation at its default settings, on data of ordinary size, in a design that is common in comparative genomics. We state what we would now require of any branch-partitioned selection test, tying each recommendation to the observation that forces it.

### Run a random-label control on your own data before reading any result

Draw labels of the same size at random from the tips that do not carry the trait, on the genes under test or on single-copy genes with no reason to differ, and fit the same test. On our three panels the two-ratio test rejected 65.9, 68.8 and 88.8% of genes under such labels (Results 2). A rate that far above the nominal level means that, in that design, the test cannot tell the trait from an arbitrary set of branches, and the trait result should not be read. The control costs one further fit per gene and needs no simulation.

### A worked example on a published result

We applied the control to a published study of the epithelial sodium channel genes of up to 91 mammals, which tested marine lineages for relaxed selection with RELAX (Zahnow et al., 2025). The study’s central conclusion, reduced constraint on *SCNN1D* in marine mammals, rests mainly on tests with larger test sets, which have too few same-shaped alternatives on these trees to be read this way, and the example does not bear on it. Among its 23 deposited RELAX tests, four were significant and had a test set with more than one alternative of the same shape on the same tree, a single otter branch or the sirenian two-tip clade with its stem; we refitted each with the deposited alignment and tree and the published command (one argument renamed for HyPhy 2.5.101; the authors’ release is not stated) under three seeds, and moved each reproduced label in turn to every lineage of its shape among the reference branches (Supplementary S8.8). One test could not be run: the deposited alignment carries in-frame stop codons, and the edited alignment the published run used was not deposited. One did not reproduce because none of our runs reached the published optimum, which our runs therefore neither confirm nor contradict. In one, the published output shows that RELAX’s own check for a flat likelihood surface found a fit with the opposite direction of *K* that was 0.19 log-likelihood units higher; the program reported the first fit, the significance holds under either (*P* = 0.026 under the better one), and our runs also gave *K* < 1, so on these data the direction of *K* is weakly identified. The fourth, relaxation in the sirenian *SCNN1A* lineage, reproduced, and its statistic exceeded that of all 12 same-shaped alternatives (7.23 against at most 1.02), none of which RELAX called significant. But 11 of the 12 are shorter than it, and 13 labels cannot support a rank-based *P* value below 1/13, so the control is consistent with the published result but cannot, with so few alternatives, show it to be exceptional. On the way it found a direction of *K* that the data barely identify and a deposited alignment that does not load as deposited (Box 1).

### Ask the question the hypothesis needs

On the real genes of our three panels a difference in *ω* between branch sets was the usual finding whatever the labels, so a test for one is answered yes more often than not; on the same genes and random labels the branch-site test, which asks only for positive selection on the labelled branches, rejected at most one of 160 genes and none after refitting (Results 3). Where the hypothesis is positive selection on the trait lineages, test for that. On alignments simulated from ten of the same genes per panel it detected positive selection of the sizes we placed on the labelled branches in 95 to 100% of replicates and rejected 3 to 5% without it (Results 3); that is power on data drawn from the fitted model, likely optimistic for real alignments, against an effect shared by all the labelled branches, and the absence of an excess is not a demonstration that the test is calibrated on them.

### Before running an empirical null, compare its resolution against your multiple-testing threshold

With *n* null genes the smallest attainable empirical *P* value is 1/(*n* + 1). Our two nulls held 82 and 81 genes and we intended twenty-seven tests, which fixes the smallest attainable corrected value at roughly 0.33. No family-level claim could survive Bonferroni correction whatever the data showed. We did not compute this in advance; the check costs one line of arithmetic and should be reported in the methods.

### Do not substitute one implementation for another; generate a null under your own design

On the 81 null genes all the branch-level arms returned, the two-ratio branch model rejected 66.7% against 35.8% for RELAX, and Clade Model C 45.7%; a third program, testing the same null site by site, rejected at 9.18% of sites against a nominal 5%, and the three failed on different genes and under different labelling schemes (Supplementary S8.2).

A reference generated by the same procedure under the same design returns the nominal rate whenever the reference and the data come from one generator, which a simulation guarantees and an observed alignment does not (Results 3). On real null genes in two further panels a reference simulated from each gene’s fitted model still rejected 52.9 and 76.1% of genes, and in an exploratory follow-up a reference simulated at the depth of the data from the null model of the test itself rejected more, 65.0 and 83.8%, while a branch-site test of positive selection on the labelled branches showed no excess on the same genes (Results 3). Imposing the real gaps on the simulated alignments changed no gene’s decision (Results 3). What the real alignments carry and the simulations do not is plausibly variation in *ω* among branches, which makes a random set of branches genuinely differ from the rest: a reference simulated under each gene’s fitted per-branch *ω*, refitted with fresh random labels, rejected 1 of 40 genes where a control carrying only the noise of that fit rejected 28 (Results 4). That reference is an upper bound, and on every gene it gives internal branches a higher *ω* than terminal ones, so part of what sets a random terminal labelling apart may be its position in the tree alone. The empirical null we applied to the tested families, built from real genes under the same labelling, carries such variation by construction, but its calibration on real data remains untested.

Matching the design is therefore not sufficient (Results 3). Nor is it the same as permuting the labels. A permuted labelling is not structure-preserving, and a structure-preserving relabelling (Saputra et al., 2021), which we use on the count axis, varies the assignment while holding the design; the reference we recommend holds the assignment and reproduces the design, together with the variation in *ω* among branches that real genes carry (Supplementary S8.7).

The device itself is old: referring a likelihood-ratio test to a parametric bootstrap is standard, and Halabi et al. (2021) applied it to a test of this family, correcting type I error rates of 15 to 21% measured on simulated data. What we add is that on real genes random labels of the size of a trait set are rejected in three panels (Results 2); that a reference from the fitted model did not restore the nominal rate even when it reproduced both the design and the depth of the data, whereas one carrying each gene’s fitted variation in *ω* among branches did (Results 3, Results 4); and that the error rate moved with the number of labelled tips on real genes (31.9 to 47.8%) with the gene set fixed, and with their placement on simulated data, in two codebases that responded to the same change of design in opposite directions (Results 5). The short-branch effect of Hughes and Friedman (2008) and the whole-tree confound of Kowalczyk et al. (2021) measure different quantities and are complementary to ours (Supplementary S8.6).

We are not proposing a device of our own. Contrast-FEL ships with a label-permutation step on significant sites that took the site-level rejection rate from 9.18 to 0.478% on our null genes (Supplementary S5.3), evidence that referring the statistic to a reference generated from the data is practicable, though at its resolution the step cannot license a claim at a corrected threshold. The recommendation is that the two programs lacking any such step acquire one.

The correction buys size, not power, and in simulation only: on the divergence-matched replicates the loss is severe and on the shallower ones slight (Results 5, Figure 4b), though on our real null genes at the same test fraction the ninety-fifth percentile of the null statistic is 33.2, against 152.6 on those matched alignments, so the losses we measured are the worst rather than the typical ones. The cost is worth paying when a study intends to name particular genes and build on them, as ours did; it is not worth paying for an exploratory screen whose output is followed up experimentally. What is never defensible is the third case, which is where we were: running the uncorrected test, reporting the survivors as findings, and leaving the reader to assume the nominal rate applies. The choice is between correcting and not claiming, not between correcting and claiming anyway.

Two forms of the correction do not carry the same requirements. Referring the statistic to a percentile of the null distribution has no lower bound on the *P* value it can support, but an undersampled null can be less conservative than no correction at all (Supplementary S8.3). Reporting an empirical *P* value has a bound, 1/(*R* + 1) from *R* replicates, so *R* ≥ 1/α - 1: nineteen replicates for a single test at 0.05 and 539 for 27 tests at a Bonferroni-corrected level. Whether the reference must match the gene set as well as the design we cannot say (Supplementary S8.4). At a median of 64 minutes per RELAX fit on our panel of 51 taxa the 539 replicates come to about 570 processor-hours; ten thousand tests would demand about 200,000 replicates per labelling design, some 24 processor-years (Supplementary S8.5). The correction is thus affordable for a targeted analysis of tens of genes at a single design, and not at present at genome scale.

**Box 1. Reading a trait-labelled branch test.** (1) Before reading the trait result, fit the same test with every set of branches of the trait set’s shape on the same tree, or with random sets of its size, and report where the trait set ranks; report its rank among sets of comparable total branch length as well, since a set with fewer substitutions gives the test less to detect. (2) Compute the rank-based *P* value as (1 + the number of alternatives at or above the trait set)/(*N* + 1); it can reach 0.05 for a single test only with *N* ≥ 19 alternatives. With fewer, report the rank. (3) Fit every set, the trait set included, under the same several seeds or starting values and take the highest log-likelihood under each model; in RELAX output, compare the fits for both directions of *K* and read the direction from the better one. (4) Check the alignment for in-frame stop codons and the reading frame before fitting, and deposit the alignment actually fitted. (5) Where the hypothesis is positive selection on the trait lineages, test for that directly. (6) Before building an empirical null, check that its smallest attainable *P* value can survive the planned correction for multiple tests. Steps (1) to (4) are implemented for RELAX in the software labelrank (Data availability).

## DISCUSSION 3 — THE STATE OF THE BIOLOGICAL QUESTION

The argument that ectomycorrhizal fungi retain a capacity to oxidise soil organic matter to mobilise nitrogen rather than to acquire carbon (Lindahl and Tunlid, 2015; Pellitier and Zak, 2018) rests on the observation we reproduce here: these lineages carry markedly fewer plant-cell-wall-degrading enzyme genes than free-living decomposers (Kohler et al., 2015; Miyauchi et al., 2020). Our contribution to that debate is narrower than we intended.

The depletion is real and large, and robust to every sensitivity analysis and reassignment we could apply, though about a third of it on the log scale is a general reduction of carbohydrate-active enzymes and the within-genome share is the robust quantity (Results 1); but a gene count does not measure capacity, and bridging counts to capacity is what our selection analysis was meant to do and what we have withdrawn. In the one case we know of in which a retained plant-cell-wall-degrading enzyme has been characterised functionally in an ectomycorrhizal fungus, its role was symbiosis establishment rather than decomposition (Zhang et al., 2022): retention is evidence of function, but not of which function. Panels like ours also carry a pseudoreplication that a phylogenetic correction does not remove: the ectomycorrhizal habit has arisen in many independent lineages (Tedersoo and Smith, 2013), yet our 119 ectomycorrhizal genomes fall into eighteen clades on our own tree, where labelling the same number of tips at random returns a median of 83 such clades. We have bounded that only for our one count estimate, which lies outside a thousand relabellings that span the observed clustering (Results 1). Supplementary S4 sets out both positions, the characterised enzyme, that clustering measurement, and the direct measurement of activity that would settle the question. Our count result is consistent with organic nitrogen mining by a reduced toolkit and equally consistent with its absence.

## DISCUSSION 4 — LIMITATIONS

### The calibration was measured over a bounded range of alignment geometry

Our null distributions were built from single-copy orthologues present in all 51 species, so every null alignment carries 50 or 51 sequences and a test set of 21 of 99 branches, while the enzyme entries they calibrate span 43 to 902 sequences and test fractions from 0.047 to 0.255. Both axes are examined in Results 5: the labelled-set axis moves the rejection rate, and a second null of forty enzyme families extends the depth axis to 342 sequences without our detecting a dependence on depth. With forty families that test had little power, so the depth axis is better described as untested at useful power than as flat, and beyond 342 we cannot say whether the rate holds. Calibration figures here were measured between 49 and 342 sequences and should not be transferred beyond that range.

### The coding subset is not a random sample of the panel

Copy numbers are available for 183 genomes but codon alignments for 51, and the difference is driven by which assemblies have usable transcript-to-protein mappings rather than anything biological. The count analyses and the selection analyses therefore rest on overlapping but different samples, and we have not reweighted one to the other.

### Intactness is conditioned on survival

(Supplementary S2.5), and the decomposition in Results 1 inherits that condition; we present it as a correction of our own earlier inference rather than a result about enzyme evolution.

### The evidence concerns three implementations, and only one of them was judged beyond the fungal panel

Other branch-partitioned tests, aBSREL and BUSTED among them, were not run. The two-ratio results were repeated on 50 yeasts and 50 lactobacilli under rules registered in advance (Results 2), and the excess rejection of random terminal labels, the larger rejection under internal labels and the dependence of the reference on the design all recurred. RELAX was fitted in both new panels but, as registered, not used for a verdict, since its fits do not reproduce; Contrast-FEL rests on the fungal panel alone. Neither form of the reference is validated on real data: the parametric bootstrap failed on real genes with both generators we ran, and the empirical null we apply has not been tested against a known null on real alignments. The branch-site test was run only on the new panels, after their registered verdicts were known, and its absence of excess rests on fits that differed widely between starting values; its power was measured only on data simulated from its own fitted model, for ten genes per panel. The new panels share the fungal panel’s size of 50 or 51 sequences, so they extend the result across lineages and placements of the trait, not across alignment sizes.

### Our simulated rate for RELAX is not the published one

Its authors rejected 52 of 1,000 alignments simulated under the RELAX null model over a γ-proteobacterial phylogeny, an estimated type I error rate of 0.052 (Wertheim et al., 2015), where our shallow set rejected 22.3% and our divergence-matched set 17.5% (Results 5). We have not established the cause: the two simulations differ in tree, labelled set, parameter values and program version, and the comparison bears on how far a calibration measured on one design transfers to another rather than on whether either is correct.

### The null genes are not enzymes, and the control for that is a small one

Single-copy orthologues serve as a null because they have no reason to be relaxed in ectomycorrhizal lineages specifically, but they differ from carbohydrate-active enzyme families in copy number history, paralogy and alignment depth. The control that difference calls for, forty non-plant-cell-wall enzyme families labelled by the same procedure at the same test fraction, agrees with null C, but at forty families the interval around that agreement is wide enough that it does not exclude a gene-set effect as large as the design effect (Results 5).

### One smaller reservation

Several of our phylogenetic models return λ at a boundary; consistency with our own recommendation requires that we not treat those values as evidence, and we do not.

Finally, the panel remains dominated by one guild by construction, and the unpublished genomes within it constrain what can be redistributed alongside this analysis. The count result does not depend on them; the selection analyses do.

## FUNDING

This work was supported by Korea Basic Science Institute (National Research Facilities and Equipment Center, 2021R1A6C101A416) funded by the Ministry of Education, the Glocal 30 program of Kyungpook National University, funded by the Ministry of Education(MOE), and the ANCHOR program through the Daegu ANCHOR Center, funded by the Ministry of Education (MOE) and the Daegu Metropolitan City, Republic of Korea (2026-ANCHOR-03-001).

## ACKNOWLEDGEMENTS

A portion of these data were produced by the US Department of Energy Joint Genome Institute (https://ror.org/04xm1d337; operated under Contract No. DE-AC02-05CH11231) in collaboration with the user community. A portion of this research was performed under the Facilities Integrating Collaborations for User Science (FICUS) program (proposals: 10.46936/10.25585/60008430 and 10.46936/10.25585/60008431) and used resources at the DOE Joint Genome Institute (JGI) (https://ror.org/04xm1d337) and the National Energy Research Scientific Computing Center (NERSC) (https://ror.org/05v3mvq14), which are DOE Office of Science User Facilities operated under Contract No. DE-AC02-05CH11231, and we thank David Catcheside for permission to include the one use-restricted genome in the panel.

## AUTHOR CONTRIBUTIONS

M.K. designed the analyses, performed the genome annotation, phylogenetic and selection analyses, wrote the simulation and calibration code, and drafted the manuscript. J.-H.S. supervised the work and revised the manuscript. Both authors read and approved the final version, and both authors agreed to the withdrawal of the claims described in Results 6.

## COMPETING INTERESTS

The authors declare no competing interests.

## PREPRINT

This manuscript is version 3 of a preprint (doi:10.64898/2026.07.20.739482). Version 1 (Kim and Shin, 2026a) reported relaxed selection on GH6, GH7 and GH28; version 2 withdrew those claims; this version reports in full the analyses behind the withdrawal, whose grounds are set out in Results 6. The reproducibility failure referred to throughout is documented in a companion manuscript (Kim and Shin, 2026b, doi:10.64898/2026.09.20.751138), posted 25 September 2026.

## DATA AVAILABILITY

All data and code underlying this article are available from Zenodo at doi:10.5281/zenodo.24034526 (all versions: doi:10.5281/zenodo.22192658; the version described here is 1.9), the code comprising every analysis, simulation, verdict and deposit-assembly script and the HyPhy batch files. The 78-gene simulation panel regenerates from the seeds that the code on Zenodo sets for each replicate.

The software labelrank, which implements steps (1) to (4) of Box 1 for RELAX and reproduces the sirenian *SCNN1A* reading of Supplementary S8.8 from the same runs, is available under the MIT licence from Zenodo (version 0.5.0, doi:10.5281/zenodo.24035697) and from GitHub (https://github.com/sunsungkim04-sys/labelrank).

The registration of the two further panels is at https://osf.io/cez4f, under embargo until 1 March 2027; its four files, the registered text, the verdict script, its checker and a list of their checksums, match the checksums recorded in the deposit.

The fungal-panel data were assembled by scripts/build_deposit.py and are those of version 1.5 of the deposit, to which version 1.7 adds the per-replicate record of the divergence-matched simulations and version 1.8 the two exploratory analyses of the labelling design (S7.6, S7.7) and the material of Figure 3, and version 1.9 the power analysis of the branch-site test (S7.4) and the worked example of Discussion 2 (S8.8), whose input alignments are those of Zahnow et al. (2025; Source Data 6, doi:10.5281/zenodo.15255678) and are not redeposited; its MANIFEST pairs each canonical result file with the script that regenerates it. They comprise 157 codon alignments, deposited untrimmed and trimmed as 313 files (the subsampled GH5 alignment untrimmed only), restricted to the 35 published genomes among the 51 with coding sequence available: the trimmed set retains 9,620 sequences, and those belonging to the 16 genomes unpublished at the time of analysis are withheld. Trees carry no sequence and keep every tip, including those of the unpublished genomes: 158 gene trees and 118 labelled trees for the analysed entries and the orthologue null genes, the 88 null C labelled trees, 40 for the enzyme-family control and five for the internal labelling. The 304 simulated alignments that no seed can reproduce are deposited as well, held in three subdirectories by the set they belong to; the 312 of the 78-gene panel regenerate instead, as above. For the count analyses the deposit carries the per-genome enzyme count table, the 182-tip species tree, the lifestyle assignments before and after the annotation audit, and the fitting scripts, which together regenerate every estimate in Results 1. It further contains the 30 canonical summary result files behind the figures, tables and reported numbers of this manuscript, the eight figure audit tables recording every plotted value at full precision, and 171 analysis scripts.

**Two classes of artefact cannot be regenerated from a seed, and are deposited for that reason rather than for convenience.** For null C, the routine that drew the random test sets combined its seed constant with Python’s built-in hash() of the gene name, which is salted per interpreter process, so re-running that script does not reproduce the draws that were analysed; the 88 null C trees and the five internal-labelling trees define those analyses and are deposited individually. Draws for the enzyme-family control and for the test-fraction series key a CRC32 checksum of the family or gene name instead and do reproduce exactly from the seeds recorded in Supplementary S3; they are deposited nonetheless.

The simulated alignments are the stronger case. The generator passes no seed to the sequence simulator, which therefore drew from an unseeded stream, so neither we nor anyone else can reproduce the replicates that were analysed. All 304 are deposited individually, 160 shallow and 144 divergence-matched. The file name of each gives its template gene and generating *K*; the shallow replicates were simulated on the template gene tree without rescaling, and for the divergence-matched ones a per-replicate record gives the branch-length scale and the target and realised pairwise identity. They define Results 5 and the calibration of the empirical null, and the reproducibility rate reported in the companion manuscript (Results 1). We describe both defects rather than silently depositing the outputs, because they are instances of the class of problem this paper is about.

For the two panels registered in advance (Results 2), the data hold the registered text and its checksum, an English record of every amendment and result, the species and gene selections, including the pool of 55 lactobacilli from which 50 were drawn, and, for the 80 null genes of each panel, the alignments, the labelled trees and the full codeml and RELAX outputs. For every simulated replicate they hold the seed and the log-likelihoods on which the verdicts were computed, and they hold the output of the independently written parsers that reproduced every reported count (the parsers are on Zenodo). The commands in newpanel/README.md regenerate byte for byte the eight verdict files that carry the nine verdicts, using the analysis script on Zenodo unchanged for every verdict; the simulated alignments regenerate from the recorded seeds. The locked registration and the verdict files are in Korean, and the README gives a glossary. The yeast and lactobacillus genomes are not redistributed: they are available from Opulente *et al*. (2024; doi:10.25452/figshare.plus.c.6714042) and from NCBI RefSeq under the accessions listed in the data. The raw HyPhy outputs behind the reproducibility analyses of the companion manuscript are on Zenodo only.

For the exploratory follow-up (Results 3), the data hold the plans, the deviation record and the checksums written before each run, the seed of every simulated replicate, the codeml output of every fit, reduced as for the registered panels, the HyPhy output of the per-branch fits, the independent re-counts and the six result files, which are in English; the simulated alignments regenerate from the seeds, and the gapped alignments of the gap check from them and the real alignments.

Genome assemblies, predicted proteomes and coding sequences were obtained from the JGI MycoCosm portal and from NCBI and are not redistributed here. Portal identifiers for all 51 genomes are listed in Supplementary Table S9. Of these, 16 were unpublished at the time of analysis, meaning that their portal pages carried no associated publication. Consulting the portal’s own use-restriction flag on 31 August 2026 identified one of the sixteen, *Amanita* aff. *grandis* KIS10, as use-restricted; the other fifteen were not so flagged. That genome was sequenced under JGI proposal 1956 (award doi:10.46936/10.25585/60001059, "Acquisition of the sequestrate (truffle like) habit by basidiomycete macrofungi"), and it is used here with the permission of the proposal’s principal investigator, David Catcheside, who asked that the sequencing project be cited: the high-molecular-weight DNA extraction methodology it developed is described in Burgoyne *et al*. (2025), and the genomes it produced are described in Nilsen *et al*. (*Communications Biology*, pending). Those genomes are identified in Supplementary Table S9 and readers should obtain them from the portal directly. The count-based analyses do not depend on the unpublished genomes: restricting the primary contrast to public genomes alone gives 0.411 (0.316–0.535). The selection analyses do depend on them, since all sixteen carry coding sequence and lie inside the 51-genome codon panel, whose other 35 members are published.

Software versions and annotation database releases are given in Methods and Supplementary S3. No new sequence data were generated.

## Notes

https://github.com/sunsungkim04-sys/ecm-pcwde-erosion

https://doi.org/10.5281/zenodo.24034526

https://doi.org/10.5281/zenodo.24035697

### Competing Interest Statement

The authors have declared no competing interest.

### Summary of Updates

Title changed to reflect the new central question. The manuscript now asks how branch-partitioned selection tests behave when the labels carry no trait. New analyses: random terminal labels on presumed-null single-copy genes in three panels of about 50 species (fungi, and yeasts and lactobacilli registered in advance), tested with the PAML two-ratio branch model; references simulated from each gene's own fitted null model, including a generator matched to the depth of the real alignments; the branch-site test and its measured power; simulations under per-branch omega; and simulations under correct specification. The retraction of the relaxed-selection claims for GH6, GH7 and GH28 made in version 1 stands. The gene-count results are retained and the within-genome ratio is now also reported. The detailed account of RELAX reproducibility is now reported mainly in a companion preprint (doi:10.64898/2026.09.20.751138). Introduction, Methods, Results, Discussion, figures, tables and Supplementary Material revised accordingly. Data and code updated to Zenodo version 1.9 (doi:10.5281/zenodo.24034526). Author name display corrected.

