## Supplementary Material for "Branch-Partitioned Selection Tests Reject Random Labels on Real Genes, Even Against a Null Simulated at the Depth of the Data"

### Supplementary information

#### SUPPLEMENTARY S1 — PROCEDURAL DETAIL FOR METHODS M1–M4 (PANEL, ANNOTATION, CATALYTIC-RESIDUE SCORING, CODON ALIGNMENTS)

References cited in these supplementary sections are listed in the References of the main text.

*Supports M1 (Genome panel, quality control and orthology)*

**The panel as first described in Results 1.** Of 196 genomes identified from the mycorrhizal and comparison sets, 186 were retrievable. Proteome completeness was assessed with BUSCO against the fungal orthologue set; 183 met the 85% threshold, two *Suillus* assemblies fell below it, and one genome yielded no proteome. Orthology inference across the retained proteomes yielded 90,236 orthogroups with 92.7% of genes assigned.

Two panel sizes recur here and are not interchangeable. 183 genomes passed quality control and carry copy-number annotations. 182 are represented in the species tree and therefore in every phylogenetically corrected analysis; one genome could not be placed. Every count below states which panel it refers to, and no comparison reported here mixes them. A third and smaller panel underlies the selection tests. Coding sequences suitable for codon-aware alignment were recoverable for 51 species, 21 of them ectomycorrhizal. This subset is not a random sample of the 182, and Discussion 4 takes up the consequences.

Catalytic-residue assessment of AA9 requires mature-chain re-scoring after signal-peptide removal, which was not completed; it is therefore included in the copy-number analyses and excluded from the intactness analyses of Supplementary S2.5.

**The procedure as summarised in the main text, in full.** Fungal genome assemblies and predicted proteomes were obtained from the JGI MycoCosm portal and from NCBI, placing ectomycorrhizal (ECM) symbionts alongside free-living and other symbiotic comparators. Completeness was assessed with BUSCO 6.1.0 in protein mode (Manni et al., 2021) against the fungi_odb10 lineage dataset, leaving a 183-genome panel; orthogroups and a genome-scale species tree were inferred with OrthoFinder 3.1.5 (Emms and Kelly, 2019), which placed 182 of the 183 genomes. The retention threshold and the species-tree inference are given below.

Including the unplaced genome (Xylhyp1, a saprotroph carrying a large plant-cell-wall-degrading enzyme complement) in otherwise tree-based comparisons would change the comparator mean, so all phylogenetic models, and every count comparison that is phylogenetically corrected, use the 182 tree-placed genomes. Two descriptive, uncorrected comparisons in Results 1, the raw mean count ratio and its AA9 sensitivity, are reported on the 183-genome annotation panel and are labelled as such. Each figure and table states which panel it refers to. Lifestyle assignment and the literature audit that corrected three of them are described in M6.

Assembly and gene-model completeness was assessed with BUSCO in protein mode against the fungi_odb10 lineage dataset (Manni et al., 2021). Genomes reaching at least 85% complete BUSCOs were retained, giving the 183-genome panel used for annotation and for catalytic-residue scoring.

**The species tree.** OrthoFinder was run in its default multiple-sequence-alignment mode, which aligns the single-copy orthogroups with FAMSA, concatenates them, and infers the tree with FastTree (Price et al., 2010); the rooting is OrthoFinder's own, from gene-duplication events.

Branch lengths are therefore in amino-acid substitutions per site, and every phylogenetic model here takes that tree as given rather than re-estimating it.

*Supports M2 (CAZyme annotation and the definition of PCWDE)*

**The procedure as summarised in the main text, in full.** Carbohydrate-active enzymes were annotated in the QC-passing proteomes with run_dbcan 5.2.8 against the dbCAN HMM library (Zheng et al., 2023), and counts were tabulated per genome at family and subfamily resolution. Plant-cell-wall-degrading enzymes (PCWDE) were defined following CAZy (Lombard et al., 2014) as the 23 glycoside hydrolase, polysaccharide lyase, carbohydrate esterase and auxiliary-activity families directed at plant cell wall polysaccharides. A protein was counted once per family it carried, and families were summed per genome to give the raw PCWDE count. The invocation, its thresholds and the family list are given in Supplementary S1.

Subfamily resolution was recovered from the model names rather than by re-annotation, and subfamily activities follow Aspeborg et al. (2012) and CAZy (Supplementary S1).

AA9 (lytic polysaccharide monooxygenases) is counted but excluded from catalytic-residue scoring and from every selection analysis, for the reason given in M3. Because the counts and the integrity analyses therefore cover different families, the count-based comparison was repeated with AA9 removed from every genome; the result is reported in Results 1 as a difference of mean counts, not a phylogenetic model.

run_dbcan/dbCAN3 was invoked as run_dbcan CAZyme_annotation --mode protein --methods hmm at the tool's default HMMER thresholds. The HMM method was used alone rather than in combination with DIAMOND or dbCAN-sub, so that every call rests on one reproducible criterion.

The 23 PCWDE families: the glycoside hydrolases GH5, GH6, GH7, GH10, GH11, GH12, GH28, GH43, GH51, GH53, GH74 and GH93; the polysaccharide lyases PL1, PL3, PL4 and PL9; the carbohydrate esterases CE1, CE5, CE8, CE12, CE15 and CE16; and the auxiliary-activity family AA9.

**Subfamily parsing.** The dbCAN HMM library names its models at subfamily resolution where subfamilies exist (GH5_5, GH5_15). An earlier version of our pipeline applied a regular expression that stripped the numeric suffix and collapsed all GH5 subfamilies into one bin; re-parsing the model names to retain the suffix recovers subfamily assignment from the existing annotation, with no re-annotation required.

The AA9 sensitivity analysis reported in Results 1 is a difference of mean counts rather than a phylogenetic model. The annotation table underlying it does not carry AA9 as a separate column, so the PGLS version of the same sensitivity was fitted separately, on a rebuilt twenty-two-family table (Supplementary S2.4).

*Supports M3 (Catalytic-residue scoring)*

**The procedure as summarised in the main text, in full.** Four PCWDE families (GH5, GH6, GH7 and GH28) have a characterised reference enzyme carrying UniProt ACT_SITE annotations, and only those four could be scored per copy. All family members were aligned with MAFFT 7.526 (Katoh and Standley, 2013) together with that reference, and each copy was classified at the catalytic columns as intact, disrupted or fragmentary; the reference positions, the classification rule and the alanine-substituted controls that verified scorer sensitivity are in Supplementary S1. The proportions of intact copies are reported in Supplementary S2.5, and the functional PCWDE count used in the comparative analyses is the number of catalytically intact copies in GH5, GH6, GH7 and GH28 per genome.

AA9 could not be scored. The N-terminal histidine of its His brace is the first residue of the mature chain (reference Cel61B, Q7Z9M7), so in proteomes that retain signal peptides it does not occupy a homologous alignment column. This is a coordinate mismatch, not a biological finding; any intact rate our scorer returns for AA9 is an artefact of it, quoted in Supplementary S2.5 only to identify it as one. A SignalP 6.0 check (Teufel et al., 2022) is reported in Supplementary S1; the mature-chain re-scoring and its selection analysis were not completed for this manuscript.

Reference enzymes and their UniProt ACT_SITE positions, used to locate the catalytic alignment columns of each scored family:

| Family | Reference enzyme | UniProt | Catalytic positions |
|---|---|---|---|
| GH7 | *Trichoderma reesei* Cel7A | P62694 | Glu229, Glu234 |
| GH6 | Cel6A | P07987 | Asp245 |
| GH28 | polygalacturonase II | P26214 | Asp180, Asp201, Asp202 |
| GH5 | endoglucanase II | P07982 | Glu239, Glu350 |

**Classification.** The alignment columns corresponding to the reference catalytic positions were located, and every copy was classified as intact (the reference residue present at every catalytic column), disrupted (at least one catalytic column carrying a different residue) or fragment (at least one catalytic column gapped because the copy does not span it).

**Alanine controls.** Scorer sensitivity was verified with synthetic controls rather than assumed. For each family, a copy of the reference in which every catalytic residue had been mutated to alanine was added to the alignment; in all four families the unmodified reference scored intact and its alanine mutant scored disrupted, confirming that the scorer reads the intended columns.

The proportions reported in Supplementary S2.5 are per-genome means of the within-genome proportion, not proportions of pooled copies; both are given there, and both include incomplete gene models. Percentages reported per family are computed over all copies of that family in a lifestyle.

**An approach not adopted.** We first attempted to measure functional integrity by scoring each copy under a masked-residue protein language model. Three checks preceded genome-scale application. A validation set of enzymes with intact catalytic residues but confirmed loss of activity proved almost impossible to assemble; on the *Trichoderma reesei* cellobiohydrolase Cel7A, whose catalytic triad we take from the literature rather than from the two ACT_SITE positions UniProt annotates, knockouts of the three shifted the score by -9.2, -8.4 and -7.4 log-likelihood units while thirty random substitutions reached -10.8; and in a masked scan of all 513 positions those three ranked 9th, 11th and 29th, yet twenty positions exceeded the same confidence threshold. The instrument was discarded before application, and the catalytic-residue calls used instead.

**AA9 and SignalP.** In proteomes that retain signal peptides the AA9 His-brace histidine does not occupy a homologous alignment column, and the scorer reads a position that is not the catalytic one. We ran SignalP 6.0 (Teufel et al., 2022) on the AA9 sequences of the panel and confirmed that it recovers the annotated cleavage site of the reference (Cel61B, Q7Z9M7) exactly, but the mature-chain re-scoring and the corresponding selection analysis were not completed for this manuscript.

*Supports M4 (Codon alignments, the identifier bridge, and the analysed entries)*

**The procedure as summarised in the main text, in full.** Coding sequences were retrieved from JGI MycoCosm for 51 of the 183 genomes, 21 of them ECM, so every codon-based analysis is restricted to that subset while the count-based analyses use the full 182-tip panel. This is an availability boundary, not a sampling design, and the single largest structural limit on what the selection analyses can address.

Joining CDS to the proteins scored above fails silently. JGI labels proteome records by proteinId and CDS records by transcriptId, so a numeric join pairs unrelated genes or drops them while the pipeline still runs to completion and returns a codon alignment. We matched on the gene model name instead, checking the translated CDS against the proteome record (Supplementary S1).

Per-entry protein alignments of the retained copies were back-translated to codon alignments, an *entry* being one family or subfamily alignment tested as a unit; columns with more than 50% gaps were trimmed, and gene trees were inferred with FastTree 2.2.0 (-nt -gtr; Price et al., 2010); HMMER 3.4 underlies the profile searches above. Twenty-eight entries were attempted. Twenty were families, being the four reference-anchored ones and sixteen further plant-cell-wall families, together with seven GH5 subfamilies and one subsampled GH5 alignment (admission thresholds and the GH5 subsampling check in Supplementary S1). Twenty-seven returned an interpretable statistic under both starting-point settings, and every count of analysed entries in this paper is that 27; the twenty-eighth, GH5_22, produced no output under the five-starting-point setting and what happened to it is reported in the companion manuscript (Results 3). The family-level GH5 entry is not independent of the seven GH5 subfamily entries and is retained because the contrast between them is itself a result.

Copy filtering across those entries is asymmetric. Only GH5, GH6, GH7 and GH28 have characterised catalytic references, so only those four were restricted to catalytically intact copies before codon alignment; the rest were filtered on length. A length filter removes obvious gene-model fragments but not catalytically disrupted full-length copies, which are expected to evolve under weaker constraint, so the length-filtered entries should be read as upper bounds on relaxation and the four reference-anchored entries as the most conservative of the 27.

**The proteinId–transcriptId bridge.** JGI labels proteome records by proteinId but CDS records by transcriptId; these are numerically distinct identifiers that are reassigned between releases, so joining the two files on the numeric identifier pairs unrelated genes or drops them. The only field the two files share is the gene model name, the fourth |-delimited field of the JGI header. We therefore matched on model name first and used identity between the translated CDS and the proteome record as a fallback, discarding any pair whose translation disagreed with the proteome sequence. We record this because the failure mode is silent: before the correction the pipeline ran to completion and produced a codon alignment either way, but recovery was low for species whose releases were mismatched, and one family's codon set contained no ECM sequence at all, so no test branch existed and the selection test could not run, without any error being raised.

**CDS retrieval.** Coding sequences were retrieved from JGI MycoCosm, restoring tape-archived GeneCatalog_CDS files where necessary.

**Entry admission.** An entry was admitted if the 51-genome CDS set yielded at least 20 sequences of which at least five came from ECM genomes.

**GH5 subsampling.** GH5 is large enough that we analysed both the complete tree and a balanced subsample of 80 ECM and 80 non-ECM sequences (random seed 42). Both are carried through as separate entries, GH5 and GH5.sub, and both appear in the companion manuscript's table of paired entries (Table 1); the complete tree is what the main text refers to as the family-level GH5 result.

**Length filtering of non-reference-anchored entries.** For entries with no ACT_SITE-annotated reference, copies were filtered on length, excluding proteins shorter than half the median length of their family.

#### SUPPLEMENTARY S2 — THE COUNT VERDICT IN DETAIL (SUPPORTS RESULTS 1)

##### *S2.1 Decomposition into three comparisons: direct contrasts*

The three fits of the decomposition in Results 1 share a tip set and their responses are correlated, so the overlap of their intervals is not a test of the difference between the estimates; we estimated each contrast directly as the coefficient of a ratio response. The family-composition contrast is +0.158 (SE 0.026; $P = 6.1 \times 10^{-9}$) and the catalytic-intactness contrast is -0.031 (SE 0.008; $P = 3.4 \times 10^{-4}$). This is consistent with the descriptive statistic: the mean proportion of assessed copies that is catalytically intact is 0.845 in ectomycorrhizal genomes, 0.900 in saprotrophs and 0.921 in wood decayers (Supplementary S2.5).

##### *S2.2 The two relabelling nulls*

Supports the statement that the count contrast lies outside 2,000 relabellings of its own tree.

The first null draws 119 tips uniformly. It does not reproduce the clustering of the real labelling and is not meant to. The mean pairwise patristic distance among labelled tips is 2.057 for the real set and no smaller than 2.118 in any of the thousand uniform draws, so the ectomycorrhizal labelling is more clustered than every one of them. The second null simulates a Brownian trait on the tree and labels the 119 tips with the highest values, which produces clustered sets in the manner of a binary permulation (Saputra et al., 2021); this one does reach the observed clustering and passes through it, its labelled sets spanning 1.830 to 2.346 with a central 95% of 1.918 to 2.346.

Under the uniform null the central 95% of the *t* statistic lay between -1.57 and +1.42, with the most extreme of the thousand draws at -3.72; under the clustered null, between -1.83 and +1.81, with the most extreme at -3.52. The observed value is -8.94. It is more extreme than every one of the 2,000 relabellings, so the empirical *P* is at the floor the procedure allows, 1/1,001 or 0.001, under both.

*S2.3 Is the reduction specific to plant cell walls*

A depletion of one enzyme class is only evidence about that class if the genomes are not depleted in enzymes generally. The annotation supplies the control directly, since every genome's total carbohydrate-active enzyme count was recorded alongside the twenty-three plant-cell-wall families. Fitting the same model to the complement answers the question the headline estimate cannot.

| **Response** | **Fold change [95% CI]** | ***P*** |
|---|---|---|
| Plant-cell-wall families, 23 | 0.367 [0.295–0.457] | $1.3 \times 10^{-15}$ |
| All other carbohydrate-active families | 0.774 [0.682–0.877] | $9.8 \times 10^{-5}$ |
| Ratio of the two within a genome | 0.473 [0.402–0.556] | $4.4 \times 10^{-16}$ |

Ectomycorrhizal genomes carry fewer carbohydrate-active enzymes of every kind, not only the plant-cell-wall families, and roughly a third of the headline effect on the log scale is that general reduction. Entering the complement as a covariate moves the estimate from 0.367 to 0.501 ($P = 4.9 \times 10^{-13}$). The specificity nonetheless survives. Within a genome, the ratio of plant-cell-wall to other carbohydrate-active families is 0.473 times as large in ectomycorrhizal genomes as in free-living decomposers, on a response that is internally normalised and therefore free of whatever drives the general reduction.

Two properties of the assemblies could produce a spurious reduction and do not. Ectomycorrhizal and free-living genomes do not differ detectably in BUSCO completeness (medians 98.0 and 98.8%; Mann–Whitney $P$ = 0.12), and adding completeness as a covariate leaves the estimate at 0.367. They do differ in assembly contiguity, the ectomycorrhizal median N50 being 319 kb against 618 kb ($P$ = 0.003), but adding $\log_{10}$ N50 as a covariate also leaves it at 0.367. The reduction is not an artefact of how well these genomes are assembled or annotated.

*S2.4 Robustness, and the treatment of λ*

Excluding the two genomes whose lifestyle is context-dependent, and then five desert truffles as well, moves the estimate monotonically to 0.357 and 0.349, away from unity in both cases. Removing the AA9 lytic polysaccharide monooxygenases, leaving twenty-two families, moves it from 0.367 to 0.399 (0.320–0.498; $P = 1.5 \times 10^{-13}$), well inside the confidence interval of either fit.

One caution applies to our own reporting. Several of these models return λ at or very near a boundary, 1.000 in the secondary contrast and 0.000 for the intactness ratio. Elsewhere we argue that a parameter returned exactly at a boundary should be treated with suspicion, and consistency requires that we apply that to λ, not only to $K$; we therefore do not offer these boundary values as evidence of robustness. The substantive claim does not rest on λ. The coefficient, its interval and its significance are stable across every reassignment scenario and both contrast definitions.

Optimiser failures also occurred here. Two cells of the sensitivity grid did not fit under the default configuration, and the failure was silent: the fitting function returned an empty result rather than raising an error. Those cells were recovered by profiling λ over a fixed grid (Methods M6; Supplementary S3).

*S2.5 Catalytic integrity of the surviving copies*

Catalytic competence was assessed by checking whether a family's canonical catalytic residues are present at the aligned positions of each copy. Four of the twenty-three families have catalytic machinery defined precisely enough for this: GH5, GH6, GH7 and GH28. AA9 was excluded because reliable assessment requires re-scoring the mature chain after signal-peptide removal (Teufel et al., 2022), which was not completed; our earlier attempt produced an intactness rate of 0.5% that is a coordinate-mismatch artefact. A protein-language-model scorer we first built for this purpose was discarded before application, because on a reference enzyme it did not separate knockouts of the catalytic residues from random substitutions (Supplementary S1).

Within those four families most surviving copies retain their catalytic residues in every lifestyle, and the proportions run counter to the direction our earlier account implied (Kim and Shin, 2026a): averaged over genomes, 0.845 of assessed copies are intact in ectomycorrhizal genomes, 0.900 in saprotrophs and 0.921 in wood decayers. Pooling copies instead of averaging genomes gives 0.837, 0.877 and 0.913, the same ordering. The phylogenetically corrected contrast is in Results 1.

Intactness is measurable only on copies that still exist. A lineage that has lost a family entirely contributes no assessed copies; one retaining a single degraded copy contributes one. Any statement about the condition of surviving copies is therefore conditional on survival, and biased in a direction we cannot determine. This applies to Results 1's intactness contrast as much as to this section, and is why we frame that contrast as a decomposition of our own earlier claim rather than a finding about enzyme biology.

*S2.6 The count verdict in full*

Supports Results 1. The passages condensed in the main text read in full as follows.

Three genomes were reassigned before fitting. Our lifestyle assignments had been taken from source-database metadata rather than from the trait database our methods claimed; an audit of all 122 genomes originally labelled ectomycorrhizal found three that are not, namely *Morchella importuna*, *Penicillium swiecickii*, and *Kurtia argillacea*, which is ericoid rather than ectomycorrhizal. The three labels were wrong on the primary literature, so the reassignment is reported as a correction; the uncorrected assignment is retained as a sensitivity scenario (0.410, Results 1).

Ectomycorrhizal genomes also carry fewer carbohydrate-active enzymes of every kind, 0.774 times as many of the non-plant-cell-wall families (0.682–0.877), and entering that complement as a covariate moves the headline estimate from 0.367 to 0.501, so roughly a third of the effect on the log scale is a general reduction. Within a genome, however, the ratio of plant-cell-wall to other carbohydrate-active families is 0.473 times as large in ectomycorrhizal genomes (0.402–0.556; $P = 4.4 \times 10^{-16}$), on a response that is internally normalised and therefore free of whatever drives the general reduction. **The claim the counts support is the ratio, not the raw depletion**, and we report both because the difference between them is not small. The headline estimate is unchanged by entering BUSCO completeness or $\log_{10}$ assembly N50 as a covariate, moves only away from unity when genomes of ambiguous or context-dependent lifestyle are excluded, and does not rest on λ, which several of these fits return at a boundary we decline to read as evidence (S2.4). Two cells of the sensitivity grid failed silently under the default configuration, returning an empty result rather than an error, and were recovered by profiling λ over a fixed grid (Methods M6).

The earlier version of this work (Kim and Shin, 2026a) attributed the difference between its 0.41-fold change in raw copy number and its 0.55-fold change in catalytically intact copies to intactness; the decomposition in S2.8 shows that this attribution is incorrect, and the per-copy assessment of catalytic integrity on which the intact counts rest, with the survivorship condition it carries, is given in S2.5.

The prescription this paper ends with is that a statistic should be referred to a distribution generated under the design that produced it, and it would be inconsistent to apply that to the verdict we withdraw and not to the one we keep. The ectomycorrhizal labelling of this panel is strongly clustered, the 119 labelled tips falling into eighteen clades, each wholly ectomycorrhizal and none nested inside another, the largest holding 51 (Supplementary S4.5), so the 152 degrees of freedom the model reports are not 152 independent comparisons.

We therefore refitted the primary contrast under 1,000 relabellings of the same tree, holding λ at its fitted value, under two nulls: one drawing 119 tips uniformly, which is less clustered than the real labelling in every one of its draws, and one labelling the 119 highest tips of a Brownian trait simulated on the tree, in the manner of a binary permulation (Saputra et al., 2021), which reaches the observed clustering and passes through it (Supplementary S2.2). The most extreme *t* statistic among the 2,000 relabellings was -3.72; the observed value is -8.94, so the empirical *P* is at the floor the procedure allows, 1/1,001, under both nulls. The count verdict therefore survives the test the selection verdict fails, and it survives it against a permulation-style null whose clustering spans the observed value, rather than one that assumes the pseudoreplication away. We report the model's *t*-test *P* in Results 1 because it is what the model returns, but the claim rests on this comparison.

*S2.7 Panel composition, raw means and the family set*

Mean copy number per lifestyle is given in Table 1, and the ordering does not follow a simple symbiotic versus free-living division: the orchid and ericoid mycorrhizal guilds both average more of these genes than either free-living guild. Those are unweighted means over eight genomes each, with no phylogenetic correction and no test, and at that sample size we did not fit the contrast. They show only that the depletion is not a simple property of being a root symbiont, and we draw no inference about those symbioses.

Compared with saprotrophs and wood decayers combined, ectomycorrhizal genomes reproduce the depletion previously reported for a smaller panel, on the 183-genome annotation panel and before the three corrections, mean 31.7 against 96.6 copies, a ratio of 0.328 (Mann–Whitney $P = 2.2 \times 10^{-12}$). This comparison is not phylogenetically corrected; the corrected estimate in Results 1 shows a smaller depletion.

Twenty-three carbohydrate-active enzyme families, in the CAZy nomenclature (Lombard et al., 2014), were designated as plant-cell-wall-degrading before any analysis. The AA9 lytic polysaccharide monooxygenases are counted; an earlier version of this work (Kim and Shin, 2026a) wrongly stated that they were excluded from the counts as well as from the selection analyses, whereas they are excluded only from the latter and from catalytic-residue scoring (Methods M3). Removing them changes the ratio of raw mean counts between ectomycorrhizal genomes and free-living decomposers on the 183-genome panel from 0.328 to 0.345 (Mann–Whitney $P = 2.2 \times 10^{-12}$ in both), and a phylogenetic fit on the remaining twenty-two families agrees (Supplementary S2.4).

*S2.8 Decomposition into three comparisons*

The previous version of this work (Kim and Shin, 2026a) presented a 0.41-fold change in raw copy number beside a 0.55-fold change in catalytically intact copies and attributed the difference to intactness. Decomposing the comparison into three steps shows that this attribution is incorrect.

| Response | Fold change [95% CI] | *P* |
|---|---|---|
| Raw count, 23 families | 0.367 [0.295–0.457] | $1.3 \times 10^{-15}$ |
| Raw count, 4 families | 0.540 [0.446–0.652] | $2.1 \times 10^{-9}$ |
| Intact count, 4 families | 0.513 [0.433–0.609] | $1.9 \times 10^{-12}$ |

Moving from twenty-three families to four shifts the estimate by +0.172; moving from all copies to catalytically intact copies within those four shifts it by -0.026, in the opposite direction, each contrast being significant in that direction when estimated directly (Supplementary S2.1). The apparent attenuation is attributable to the change in family set, and we withdraw the implication that surviving copies in ectomycorrhizal genomes are better preserved catalytically (Supplementary S2.5). The impression was an artefact of comparing different family sets.

#### SUPPLEMENTARY S3 — PROCEDURAL DETAIL FOR METHODS M5–M6 (SELECTION TESTS, REPRODUCIBILITY, NULL DISTRIBUTIONS, SIMULATION, PGLS)

*Supports M5 (RELAX invocation)*

The canonical invocation was

```
hyphy relax --alignment <codon.fasta> --tree <labelled.nwk>  --test Test --
starting-points <n> --output <out.json>
```

--starting-points was varied deliberately and is reported for every result; unless stated otherwise the batches used --starting-points 5. Every run in this study was a single-process invocation of the multiprocessor (MP) build. No run used HYPHYMPI, so the message-passing layer is excluded by construction rather than by argument.

The three options that materially change the fitted model are off by default and were therefore off in the main analysis: synonymous rate variation (--srv, default No), multinucleotide substitutions (--multiple-hits, default None), and the alignment-error absorbing rate class (--error-sink, default No). The first two correspond directly to the model-misspecification mechanisms of Wisotsky et al. (2020) and Venkat et al. (2018), and the third is directly relevant to alignments trimmed automatically, as ours are.

*Supports M5 (Null B)*

The pipeline was MAFFT protein alignment (Katoh and Standley, 2013), back-translation to codons, removal of columns with more than 50% gaps, and FastTree gene tree inference (Price et al., 2010). 186574at4751 has a codon alignment that is not rectangular, one sequence being two nucleotides shorter than the rest, so tree inference fails and no run is possible; it was excluded on that ground alone, before any result was seen. No gene was selected or dropped on the basis of its result, so the set is the complete qualifying collection by construction. These are strongly constrained loci whose $\omega$ dynamics are not those of CAZymes; the property the null requires is not that they behave like CAZymes but that they have no reason to be relaxed *specifically on ECM branches*.

*Supports M5 (Null C, and the retired version of that control)*

Draws were made by scripts/gbe_batch1_refit.py, seed constant 20260730. The realised draws are deposited as labelled Newick files, one per gene, and it is those files, rather than the seed, that define the analysis: the drawing routine combined the seed constant with Python's built-in hash() of the gene name, which is salted per interpreter process, so re-running the script does not regenerate the same draws. We state this because it is a defect of exactly the kind this paper is about, and because the deposited trees make the analysis auditable regardless. Draws for the test-fraction series use a stable checksum instead and do reproduce from their recorded seed. All 88 draws were verified to contain zero ECM tips and a test set of size exactly 21. Alignments and trees were not recomputed, so the only difference from null B is which tips carry the label.

An earlier version of this control drew the 21 tips from all 51 genomes. Because 21 of the 51 are ECM, every draw then contained 5–13 ECM tips (mean 8.56, 41%), and the control did not isolate the design contribution it was introduced to isolate. That version is retired; it is described here because its numbers appear in the preprint of this work.

*Supports M5 (Test-fraction series)*

**As summarised in the main text, in full. Dependence on test-set size.** The nulls label 21 of 99 branches (fraction 0.212); tested families span 0.047–0.255, median 0.089. On the same genes, alignments and trees only the labelled-tip count changed, across a *test-fraction series* of eight *steps* from 3 to 29 labelled tips (fractions 0.030 to 0.293), draws restricted to non-ECM tips. Sixty-nine genes returned results at all eight steps, the common set; the fifth percentile of null *K* per step was bootstrapped over 5,000 gene resamples.

Steps of 3, 5, 9, 13, 17, 26 and 29 tips were run by scripts/testfrac_ladder.py, seed 20260803, keyed by a CRC32 checksum of the gene name and step size so that the draws reproduce. The eighth step, 21 tips, is null C itself. Draws remain restricted to non-ECM tips. Twenty-nine rather than thirty was used as the upper step because the panel contains exactly 30 non-ECM tips, so a 30-tip draw would be the exact complement of the ECM set: a structured rather than a random test set, with no draw-to-draw variance. Such a set would confound test-set size with test-set composition at that step alone. The empirical fifth percentile of the null *K* distribution at each step was bootstrapped by resampling genes 5,000 times (seed 20260804). The step table, the exact permutation test of the trend and the bootstrap are all regenerated by a single script, scripts/testfrac_verdict8.py, which prints its own comparison against the deposited result file.

Test fractions are quoted from the labelled trees as constructed. RELAX collapses internal branches of zero length by default (--kill-zero-lengths Yes), so the realised denominator is occasionally one or two branches smaller than the nominal 99. We verified across all eight steps, the null C set and the CAZyme control that this never removes a labelled tip (the realised test-set size equals the designed size in every run) and that it moves the test fraction by at most 0.003.

*Supports M5 (CAZyme control)*

**As summarised in the main text, in full. Dependence on gene set.** A further control differs from null C in gene set alone. It uses forty non-plant-cell-wall-degrading CAZyme families from the dbCAN annotations, excluding the lignin- and cellulose-active AA classes, at alignment depths of 49 to 342. Processing follows null C and labelling follows the real analysis, so the test fraction emerges from the gene tree, median 0.215 against 0.212.

Built and run by scripts/cazyme_control_build.py and scripts/cazyme_control_relax.py. Forty families were taken from the existing dbCAN annotations, excluding the AA1/AA2/AA3 lignin- and cellulose-active classes on the ground that a family with a reason to be relaxed on ECM branches is not a null; what remains is predominantly biosynthetic glycosyltransferases together with fungal cell-wall and housekeeping glycoside hydrolases. Five families were excluded as too deep to fit in reasonable time (GH16, GT2, GH18, GH13 and CE4, at 406 to 1,114 sequences). Alignment trimming, tree inference, the lifestyle assignment and the RELAX invocation are those used for null C. Labelling used CAZYME_SEED 20260804, CRC32-keyed per family. The resulting test fractions have a median of 0.215 against 0.212 for null C, so the two sets are matched on the axis the series varies and separated on the axis it cannot reach.

*Supports M5 (codeml two-ratio comparison)*

The one-ratio model (model = 0) was compared against the two-ratio model (model = 2, the labelled branches as foreground) by a likelihood-ratio test on one degree of freedom; the two models are nested at the interior point $\omega_{fg} = \omega_{bg}$, so no boundary correction applies. Runs used

NSsites = 0, F3x4 codon frequencies, an estimated transition/transversion ratio, cleandata = 0 so that the alignments are the ones RELAX analysed rather than a gap-stripped variant, and branch lengths estimated from scratch. Following our own recommendation about starting points, every model was fitted twice from $\omega = 0.5$ and $\omega = 1.5$ and the higher likelihood retained; the two starts also provide a paired reproducibility measurement in a second program, parallel to the starting-point comparison reported for RELAX.

Alignments were converted to sequential PHYLIP and the {Test} annotation to PAML's #1 foreground marker; the conversion was checked per gene by requiring the number of foreground labels to equal the number in the source tree and the number of tree tips to equal the number of sequences, and every gene passed. Each fit ran in its own working directory because codeml writes auxiliary files to the current directory and concurrent runs in a shared directory would overwrite one another silently. Of the 1,472 two-ratio fits across the four sets, none failed; the 1,152 quoted below for the truncation accounting are the fits on the three original sets, the same alignments as the clade-model count.

*Supports M5 (Clade Model C, and the truncation recovery pass)*

The alternative adds one parameter, a separate $\omega$ for the foreground branch set within one site class, and we confirmed on pilot genes that the fitted parameter counts differed by exactly one before interpreting any statistic on one degree of freedom.

These fits are far more expensive, and an initial pass did not complete. Each was capped at four hours and 161 of the 1,152 exceeded the cap, concentrated in the longest alignments and overwhelmingly at the higher starting value: 139 of the 576 fits begun at $\omega = 1.5$ did not finish, against 22 of the 576 begun at $\omega = 0.5$. Because the truncation rate differed between the observed and the simulated sets (22 of 352, 69 of 400 and 70 of 400, or 6.3, 17.2 and 17.5%), the sets would otherwise have rested on different denominators, and truncation is not random with respect to the outcome, since it discards the slowly converging fits. Every truncated fit was therefore re-run under a twenty-four-hour cap. Five of the observed-set fits were recovered in an earlier pass, leaving a final batch of 156, which completed in 30.4 hours of wall time with the longest single fit taking 14.0 hours; every one of the 161 finished, so the truncation was resource contention rather than non-convergence and the three sets are complete at 88, 100 and 100 genes. Where both starting values completed, the higher likelihood was retained, the same best-likelihood rule applied throughout. Every figure derived from these fits is reported with its denominator.

*Supports M5 (Clade Model C settings)*

Clade Model C was fitted as model = 3, NSsites = 2 with three site classes, against its null M2a_rel as model = 0, NSsites = 22 (Weadick and Chang, 2012). The two differ by exactly one parameter, so the likelihood-ratio test is on one degree of freedom. The divergence-matched set received its own recovery pass rather than the one described above for the other three sets.

*Supports M5 (Contrast-FEL)*

**As summarised in the main text, in full. A third implementation, asking the question site by site.** Null C was analysed a third time with Contrast-FEL (Kosakovsky Pond et al., 2021), which compares $\omega$ between branch sets site by site with no $K$ parameter, on its alignments and labelled trees unchanged, in HyPhy 2.5.100 (MP) at its defaults, which leave synonymous rate variation on, whereas the RELAX runs had it off, so an excess here cannot be attributed to that omission. All 88 genes returned output, 61,059 sites in all.

Run by scripts/cfel_run.py and scripts/cfel_verdict.py. We record the HyPhy version because this analysis has had repeated changes to its optimiser settings across recent releases.

Contrast-FEL reports, for each site, an asymptotic *P* value, a Benjamini–Hochberg *q* value computed across sites within the gene, and, for sites significant under the asymptotic test, a label-permutation *P* value, which the program computes by default. Sites at which the permutation test was not run carry a sentinel of -1 in that column rather than a missing value, and are excluded from any denominator involving it. Because the unit of inference is a site rather than a gene, these rates are not placed on the same axis as the gene-level rejection rates of the two branch-level tests; both a site-level rate and a gene-level count of genes carrying at least one significant site are reported, and the latter is a summary rather than a calibrated quantity.

*Supports M5 (Model-misspecification controls)*

**As summarised in the main text, in full.** Three misspecification mechanisms with documented effects, namely unmodelled synonymous rate variation (Wisotsky et al., 2020), instantaneous multinucleotide substitution (Venkat et al., 2018; Lucaci et al., 2021) and alignment error absorbed by a rate class, were controlled by re-running null C with one flag changed at a time, --srv Yes, --multiple-hits Double, --error-sink Yes, each reported on the paired intersection. The counts of genes moving between rejection and non-rejection carry more weight than the difference in rejection rate, because each gene was fitted once per arm.

Each control was re-run on the null C gene set under otherwise identical settings (--starting-points 5, same alignments, same ECM-free labelled trees), changing one flag at a time (scripts/srv_control.py, scripts/mh_es_control.py). For each control we report, on the paired intersection, the rejection rate before and after, the transition counts between rejection and non-rejection in both directions, the median change in $K$ ($\Delta K$), and the counts of two pathology indicators: runs terminating with a negative likelihood-ratio statistic, and runs returning $K = 1.0000$ exactly. Because each gene was fitted once per arm, the difference between two rejection rates carries the run-to-run irreproducibility documented in the companion manuscript (Kim and Shin, 2026b); this is stated with every such comparison and is the reason the transition counts, which are not a difference of two aggregates, carry more weight than the rates.

Aggregation used a tolerant JSON reader (scripts/arms_recount_tolerant.py). RELAX writes the bare tokens inf/-inf into the per-branch evidence-ratio block, which is not valid JSON; a strict parser raises and, in a permissive aggregator, silently drops that gene from the paired intersection. The tolerant reader rewrites those tokens and reports the number of files it recovers, so that the denominator of every comparison is stated rather than assumed.

*Supports M5 (Simulation under a known generating value)*

Built and run by scripts/sim_positive_control.py and scripts/sim_relax_run.py; the divergence-matched rebuild by scripts/sim_divcal.py, scripts/sim_divcal_perK.py and scripts/sim_matched_gen.py.

**Generating parameters.** Each template supplied its three-class $\omega$ distribution and proportions from its RELAX null fit with $K$ fixed at 1, the transition/transversion ratio $\kappa$ from its GTR fit, codon-site count from its alignment, and tree from its {Test}-labelled topology. Sequences were evolved under MG94 with F1x4 equilibrium frequencies. For the five-template set, 20 replicates were generated at $K = 1$ and 6 at each of $K = 0.5$ and 0.7. For the 78-gene simulated set the per-replicate seed is derived from gene, $K$ and replicate number, and realised pairwise distance was printed at generation as a self-check.

The bisection search was repeated separately at each generating $K$, since $\omega_{\text{test}} = \omega_{\text{ref}}^{K}$ accelerates the labelled branches whenever $\omega_{\text{ref}} < 1$ and a scale fitted at $K = 1$ therefore overshoots elsewhere. One template could not be brought to its target at any scale within the search bounds and was dropped. Only the branch-length scale is altered by this procedure; the $\omega$ classes, their proportions and κ are those of the first construction, so the generating value of *K* is preserved. Realised identities are reported per replicate alongside their targets. Internal-branch labellings for the divergence-matched set reuse the same trees as the shallow internal arm, so the two differ in the alignments and not in the labelling (scripts/sim_relax_run2.py, which takes the tree path as an argument and reports the resolved source and label counts before launching).

The paired one- and five-starting-point fits serve two purposes: they supply the convergence check on data whose generating model is exactly the fitted model, and they measure the effect of the starting-point setting on type I error under a known null. This design answers three questions that the observational analysis cannot: whether the test rejects at its nominal rate when the null is exactly true ($K$ = 1); whether it recovers and detects genuine relaxation ($K$ = 0.5, 0.7); and whether simulated data pass the same convergence criteria that the real data fail. If they do, the criterion is falsifiable and the problem lies with the data; if they do not, the problem lies with the optimiser.

*Supports M5 (Proportion conventions and aggregation)*

Two conventions for proportions run through the paper and are not the same family. Tests of a proportion against a fixed value are exact binomial; interval estimates of a proportion are Wilson score intervals, except in the two registered panels of Results 2, where the registered rule specifies exact (Clopper–Pearson) intervals. We name both because at the sample sizes involved the choice is not cosmetic: on the forty-family control of Results 5 the Wilson interval is 22.1 to 50.5% where the Clopper–Pearson interval is 20.6 to 51.7%, and a reader recomputing from the counts alone would not recover our figures.

Aggregation of RELAX output used a tolerant JSON reader, since RELAX writes bare inf/-inf tokens that a strict parser rejects, silently dropping the gene.

*Supports M5 (Lifestyle assignment of the selection analyses)*

**The two lifestyle assignments are not the same in the two halves of this paper, and the asymmetry is deliberate.** The count analyses of Results 1 use the corrected assignment, in which *Morchella importuna*, *Penicillium swiecickii* and *Kurtia argillacea* are not ectomycorrhizal; every selection analysis uses the assignment as it stood before that audit, under which all three are labelled ectomycorrhizal, so the test set is 21 tips rather than 18. The nulls against which the entries are calibrated were themselves built under the earlier assignment, and relabelling one side of a paired comparison would change two things at once. Three of the 21 labelled tips are therefore genomes we have since reassigned; the direction is conservative, since two saprotrophs and one ericoid symbiont among the labelled tips make the real labelling more like a random one, and the audit itself is described under M6 below.

*Supports M5 (The third simulated set)*

A third set extended the simulation from five templates to the null C gene set. Every gene with an interpretable RELAX null fit was classified by that fit before simulation into three strata: 57 with all $\omega$ classes below 100 and one class above one (the *moderate* stratum), seven whose highest class was at one (*no fitted heterogeneity*), and 17 with a class at or above 100 (the *boundary* stratum, named for that class rather than for the optimiser bound). Three of the last group, whose fitted class sat at the optimiser ceiling of $10^{10}$, were excluded because the generating rate matrix cannot be constructed there, leaving 78. Four replicates per gene were generated at $K = 1$ under a recorded per-replicate seed, so this set, unlike the first two, regenerates from the pipeline. It used the generator of the first set, on the gene trees as inferred, and so shares its shallow depth. The identical alignments were fitted with the two-ratio model under terminal labels and under internal labels marking the 21 longest internal branches, the rule of the five-template internal arm. Stratum definitions and the gene-averaged summary, with the moderate stratum as primary, were fixed in a dated analysis plan before the batch ran; the paired McNemar test reported for this set was not specified in that plan.

*Supports M5 (Two panels registered in advance: selection, labels, decision rules and amendments)*

The predictions, decision rules, selection rules and analysis script for two further panels were fixed and their checksums recorded before any fit was run, and the registration was deposited with the Open Science Framework (https://osf.io/cez4f), registered at 15:14 UTC on 30 September 2026, before the first fit, and under embargo until 1 March 2027. The registered text is in Korean; an English record of every amendment and result is deposited with the data. The script was first run on the outputs of the fungal panel and reproduced every fungal figure it was to be compared with.

*Yeasts.* Candidate species were those of Opulente et al. (2024) whose nuclear genetic code, as inferred by Codetta (Shulgina and Eddy, 2021), matched the standard code exactly, that did not descend from the whole-genome duplication (the most recent common ancestor of *Saccharomyces cerevisiae* and *Vanderwaltozyma polyspora* on the authors' time tree), whose genome was at least 90% complete by BUSCO, and that had growth data; one strain per species was kept. The order with most candidates, Pichiales (140 species), was sampled at random to 50 under a recorded seed. The trait was the carbon source, among those scored for all 50 species, on which 20 to 50% of species failed to grow with the most parsimony transitions on the pruned time tree, which was glycerol (19 of 50 species unable to grow); growth was read from the authors' growth rates, a rate of zero being no growth.

*Lactobacilli.* Lifestyles were taken from the supplementary assignment table of Duar et al. (2017), with vertebrate- and insect-adapted species as the trait species and free-living and nomadic species as the non-trait species, names being mapped to the current genera (Zheng et al., 2020); species absent from that table were excluded. The clade was the smallest on the GTDB R10-RS226 bacterial (bac120) tree (Parks et al., 2026) holding at least ten species of each kind and at least as many non-trait as trait species, the last condition being needed to draw the random labels below. It held 55 assigned species, 20 host-adapted, of which 50 were drawn at random under a recorded seed, leaving 18 host-adapted and 32 non-trait species; one NCBI reference genome was used per species.

*Genes, labels and fits.* In each panel the BUSCO genes (Manni et al., 2021) present in single copy in all 50 species, scored on the translated coding sequences, were sampled at random to 80 null genes, aligned and treed as for the fungal panel (M4). Each gene received its own random terminal labelling, $k$ tips drawn from the non-trait species under a seed keyed to the gene, with $k$ equal to the number of trait species (19 in yeast, 18 in lactobacilli). Fits followed the fungal panel exactly: the two-ratio and one-ratio models at two starting values of $\omega$ each, the larger log-likelihood of each being kept, and RELAX in HyPhy 2.5.100 at five starting points with the thread count fixed at one. Simulated replicates were generated from each gene's RELAX null fit as in the third simulated set of M5, eight per gene at $K = 1$ for the paired comparison of labellings, internal labels marking the $k$ longest internal branches; 100 per gene and labelling for 20 moderate-stratum and up to ten boundary-stratum genes drawn at random for the gene-specific reference distributions; 52 more per gene at $K = 1$ under terminal labels, bringing each gene's parametric bootstrap reference distribution to 60; and four per gene at each of $K = 0.5$ and 0.7 for the positive controls. The same gene-specific reference distributions were drawn for 20 moderate and ten boundary genes of the fungal panel's 78-gene simulated set with new seeds. That fungal-panel analysis was designed after an exploratory cross-design comparison, with reference distributions pooled over the 78 genes at four replicates each, had returned 4.8% in the moderate stratum; its replicates are new and independent of that exploration. Every seed was keyed to the panel, the gene and the replicate, and checked against every other seed in the study for collisions before use.

*Decision rules.* The first prediction was tested on the 80 genes under random terminal labels by a one-sided exact binomial test against 0.05. The paired comparison used a two-sided exact McNemar test at $\alpha = 0.05$ on the moderate stratum, and held only if internal labels rejected more often. A gene-specific reference distribution contained the 99 replicates of the other alignments, giving an exact size (type I error rate at the nominal $\alpha$) of 4%, and the gene-averaged rejection rate was bounded by a gene-level cluster bootstrap of 10,000 resamples; the prediction held if the lower 95% bound exceeded 5%. The real-gene reference distribution used an empirical $P$ of $(1 + n)/61$, $n$ being the number of the 60 replicates at or above the observed statistic, a gene being rejected at $P < 0.05$, and the prediction held if the exact two-sided 95% confidence interval of the rejection rate contained 5%. A fit not finished within 24 hours, or ended by an error, was not retried and was counted as incomplete against the number of fits planned before the run; a stage in which more than 10% were incomplete is flagged, and the verdict computed on the completed fits.

*Changes after registration.* Four sets of amendments were made after the lock, each before the fits it affects and each recorded with the rule it replaced; the last three were made after the first two yeast verdicts were known, though they concern only seeds and the selection of lactobacilli. The yeast data lacked the duplication flag, order assignment, binary growth calls and gene-to-coding-sequence links the selection rules assumed, and these were supplied as described under *Yeasts.* above; the BUSCO release named in the registration was replaced by 6.1.0, the earlier lineage dataset no longer being distributed. Two of these choices, the order in which strains were reduced to species and the species key, were made after the resulting trait and $k$ (sucrose and 24, or glycerol and 23) had been seen under an earlier key; applied as written, the final key gave glycerol and $k = 19$. No statistic had been computed at that point. The registration also specified that a failure of the fourth prediction would narrow the title towards an empirical null built from real genes; we narrowed it instead, first to the failure of the simulated reference and, after the exploratory follow-up, to the rejection of random labels against a null simulated at the depth of the data, since the empirical null has not been tested on real data either. The random-number key for the positive controls was amended to include $K$, which would otherwise have reused the seeds of the null replicates. The lactobacillus source table, the definition of host adaptation and the tree were fixed when the published assignments proved to be given in a supplementary table rather than in the text. And the requirement that the lactobacillus clade hold at least as many non-trait as trait species was added when the first clade, of 41 species, made the random labelling impossible to draw; that clade was set aside before any label was placed.

*Supports M5 (Exploratory follow-up)*

**Exploratory follow-up.** Five analyses were added after the fourth prediction had failed; each plan, with its scripts, was fixed by checksum before any statistic it reports existed, and none changes a registered verdict. In the first, each real null gene was simulated under its own one-ratio codeml fit (branch lengths, $\kappa$, $\omega$ and the codon frequencies codeml used) with PAML 4.10.10 evolver, which takes branch lengths in codeml's unit, 60 replicates per gene with a recorded seed; a fixed-parameter codeml run reproduced each gene's one-ratio log-likelihood to within $2.3 \times 10^{-5}$ before any replicate was written, and the depth of the replicates was checked against the real alignments by rules fixed before the check. Every replicate was fitted with the same one- and two-ratio models as the real gene, and the real statistic was referred to them as in the fourth prediction. The plan also named a second generator, the RELAX null model with branch lengths in the correct unit; it was dropped before the full run began, because those branch lengths are themselves inflated by the fitted large-$\omega$ classes. In the second, codeml's branch-site model A (Zhang et al., 2005) was fitted to the same alignments and labelled trees, with the null fixing the foreground $\omega$ class at one, from two starting values per model; after that result was known, and because the alternative had ended below the null on 18 genes, every gene was refitted from three more starting values per model, one of them started from the other model's optimum, and each model's log-likelihood was taken as the maximum over its five fits. The likelihood-ratio statistic was referred to $\chi^2_1$, and both the planned and the refitted results are reported. In the third, planned after both results were known, the 60 replicates of the first analysis for the ten genes per panel with the largest fraction of gap characters in the real alignment ('-' being the only character other than A, C, G and T) were given the real gap pattern: at every position where a taxon's real sequence has a gap the simulated character was replaced by the gap, and every other character was kept. No new replicate was simulated; tree and control files were byte copies of the replicate's, and each of the 4,800 fits completed. The rule for reading the result was fixed with the plan: the gap pattern was to be said to contribute if at least three fewer of the 20 genes were rejected against the gapped replicates than against the originals. The fourth, planned after the third result was known, asked whether variation in $\omega$ among branches accounts for the excess. Twenty of the 80 genes per panel were drawn with a recorded seed. Each real alignment, and five one-ratio replicates of the first analysis given the real gap pattern, were fitted in HyPhy 2.5.100 under MG94xREV with CF3x4 frequencies, once with one $\omega$ for the tree and once with one $\omega$ per branch, both from the same nucleotide pre-fit with its rates then held fixed and with every branch kept; the statistic was twice the difference in log-likelihood, and a gene was counted as departing if its real value exceeded all five. Two generators were built in pyvolve 1.1.0 with one GY model per branch: G3 from the per-branch fit of the real alignment, and G3n, the noise control, from the per-branch fit of the first gapped replicate, whose true variation among branches is zero. Each took the pattern of $\omega$ and branch length from the MG94 fit and the level from the codeml one-ratio fit of the same alignment, because the two models define both quantities differently; per-branch $\omega$ was capped at 100, and a branch longer than 20 substitutions per site in the MG94 fit, a run-away synonymous rate where $\omega$ is not identified, took its codeml length and $\omega$. Before any replicate beyond them was made, the first six genes per panel had to pass a check of depth (median replicate to reference ratio of one-ratio tree length and $\omega$ within 0.8–1.25 for each generator) and of the round trip of the per-branch pattern (median slope of refitted on source log $\omega$ within 0.7–1.3 for G3). Forty replicates per gene and generator were fitted with the one- and two-ratio models from two starting values each, every replicate under its own fresh random set of terminal labels of the gene's size, so that the reference does not copy any contrast on the branches the real statistic used. A gene was rejected if at most one replicate reached its statistic. The reading, fixed with the plan, required G3 to reject fewer genes than G3n (exact McNemar $P < 0.05$) with a one-sided 95% upper bound on the G3 rate of at most 15%, read as an upper bound because G3's per-branch values carry estimation noise as well as variation; it was to be read as 'noise suffices' instead if the bound on the G3n rate met the same 15%. The exception to fixing each plan before its statistics existed is here: in a dry run before the plan was fixed, one comparator value of the first statistic (on one gene) was seen, and the instrument check was revised once after it failed, on run-away branches in five genes, before any replicate beyond the check's was made; both are recorded in the plan. The program for the per-branch fits was changed from codeml's free-ratio model to HyPhy before the plan was fixed, because one free-ratio fit had not finished in more than 14 hours of CPU time. The fifth measured the power of the branch-site test on pyvolve-simulated replicates of ten genes per panel, with positive selection on every labelled branch; its plan and results are in Supplementary S7.4.

*Supports M6 (Tree pre-processing, lifestyle audit, thread pinning)*

**As summarised in the main text, in full.** Per-genome PCWDE counts were analysed against the 182-tip OrthoFinder species tree (Emms and Kelly, 2019), pre-processed as described below. Phylogenetic generalised least squares (PGLS) was fitted with caper::pgls (version 1.0.4; Freckleton et al., 2002; Orme et al., 2018) in R 4.5.3, over trees handled with ape (Paradis and Schliep, 2019), as log10(count + 1) ~ is_ECM, Pagel's λ by maximum likelihood (Pagel, 1999), with an ordinary least-squares baseline. Fold changes are the back-transformed slope, $10^{\beta}$, and the intervals reported with them $10^{\beta \pm 1.96\ \mathrm{SE}}$: a normal-approximation interval on the log scale rather than a *t* interval. The primary contrast is ECM versus free-living decomposers (SAP + WD), ECM versus all non-ECM genomes the secondary. The primary contrast was additionally referred to a null generated under its own labelling design: 1,000 relabellings of the same tree with λ held at its fitted value, drawn first uniformly and then by simulating a Brownian trait on the tree and labelling its highest tips, which spans and passes through the clustering of the real set where the uniform draw does not reach it. A specificity control fits the same model to the complement of the twenty-three families within the same annotation, every other carbohydrate-active family, and to the within-genome ratio of the two, and enters the complement, BUSCO completeness and $\log_{10}$ assembly N50 as covariates in turn. A three-step decomposition separates composition from catalytic integrity, using raw counts over the twenty-three families, raw counts for the four families with per-copy catalytic-residue calls (GH5, GH6, GH7, GH28), then intact copies of those four. Package versions, the permutation and specificity scripts, and the handling of tip-name ordering are given below.

**Software versions and label handling.** Trees were handled with ape 5.8.1 (Paradis and Schliep, 2019), phytools 2.5.2 and caper 1.0.4. The permutation null is scripts/pgls_permute.R, drawing 119 of the 154 tips in the primary contrast; the specificity control is scripts/pgls_specificity.R, whose first row reproduces the primary fit as a check. Labels are carried as tip names rather than as positions, since the tree and the data frame are not in the same order; a positional join would silently mislabel every genome.

**Which assignment the selection analyses use.** The selection analyses retain the pre-audit lifestyle assignment, under which three genomes since reassigned are still labelled ectomycorrhizal: three of the 21 labelled tips, 14% of the test set. This does not bear on the conclusions of Results 2 to 4 and Results 5, which concern the behaviour of the tests under labellings we generated ourselves and under simulation; but a reader comparing the two halves of the paper should know that the ectomycorrhizal set differs between them.

**Tree pre-processing.** The tree was pre-processed by dropping node labels (support values otherwise collide with tip labels in caper), resolving polytomies with ape::multi2di (Paradis and Schliep, 2019), and setting non-positive branch lengths to $1 \times 10^{-8}$.

**Lifestyle audit procedure.** All 122 genomes assigned to ECM were re-checked against the primary literature at genus level, and the seven genomes carrying the highest PCWDE complements were checked individually. Three assignments were wrong and were corrected:

Morco1 (*Morchella importuna*) and Penswi1 (*Penicillium swiecickii*) to saprotroph, and Kurarg1 (*Kurtia argillacea*, an ericoid mycorrhizal fungus of the Hymenochaetales, not the Atheliales) to ericoid mycorrhizal. The corrected assignment is the primary scenario for the count analyses, with the uncorrected fit reported as a sensitivity scenario so that the effect of the correction is visible rather than asserted.

**Optimiser recovery and thread pinning.** Two scenarios failed with optim ABNORMAL_TERMINATION_IN_LNSRCH under ML estimation of λ; *P* values below double precision are reported as $< 2.2 \times 10^{-16}$ with *t* and degrees of freedom. Failed fits were recovered by profiling λ over a fixed grid (0.05 to 1.00 in steps of 0.05) and retaining the maximum-likelihood grid point, and every reported fit records which path produced it. We report this because a manuscript whose subject is optimiser reliability cannot pass over an optimiser failure in its own comparative analysis. The linear-algebra library underlying these fits is multithreaded by default, and on a shared machine it spawned one thread per core. We therefore refitted the primary contrast and the three-step decomposition with the BLAS thread count pinned to one (OMP_NUM_THREADS, OPENBLAS_NUM_THREADS and MKL_NUM_THREADS all set to 1) and confirmed that the estimates, confidence intervals, *P* values and λ agree with the unpinned fit to four decimal places. We report this because the reproducibility failures documented in the companion manuscript (Results 1) raise the obvious question of whether our own comparative analysis is subject to them, and the answer here is that it is not: unlike the codon-model optimiser, this fit returns the same answer under both thread configurations.

**Data-availability sensitivity.** The comparative analysis was repeated on the subset of genomes that have an associated publication by scripts/pgls_published_only.R. The restriction removes 61 of the 119 ectomycorrhizal genomes and one of the 35 comparators, the *Penicillium* the annotation audit had reassigned from ectomycorrhizal to saprotroph, leaving 58 against 34.

#### SUPPLEMENTARY S4 — SUPPORTING DETAIL FOR DISCUSSION 3 (THE STATE OF THE BIOLOGICAL QUESTION)

##### *S4.1 How the nitrogen-mining position is usually paraphrased*

Supports the statement of what the argument about ectomycorrhizal decomposition actually claims.

The position is often paraphrased as "facultative" or "conditional" saprotrophy, including in our own earlier work. Its authors explicitly decline that framing: their claim is that decomposition serves nitrogen acquisition, not that these fungi feed saprotrophically. The distinction matters for what gene counts can bear on, because a reduced enzyme set is a weaker objection to nitrogen mining than it is to saprotrophic nutrition.

##### *S4.2 Why gene content is not capacity*

Supports the statement that counts cannot be bridged to capacity.

A copy that is present may not be expressed, expressed protein may not be secreted, and secreted enzyme may not act on the substrate that matters in soil. Conversely a reduced set may be sufficient: an organism that has retained the enzymes it uses and shed those it does not is not thereby less capable of the activity it retains. Bridging from counts to capacity was precisely what our selection analysis was supposed to do, and it is what we have withdrawn.

##### *S4.3 The evidence on both sides of the debate*

Supports the statement that both positions are thinner than the literature suggests.

The case that ectomycorrhizal fungi have lost decomposition capacity rests substantially on gene-count comparisons of the kind we report, and gene counts do not measure capacity. The case that they retain it rests on a smaller set of transcriptional and spectroscopic observations from a handful of species grown on organic-matter extracts (Shah et al., 2016). Neither body of evidence is at the scale of the claim it is used to support, and both are drawn from taxonomic samples shaped by which genomes were sequenced.

*S4.4 The characterised retained enzyme*

Supports the statement that retention is evidence of function but not of which function.

In the one case we know of in which a retained plant-cell-wall-degrading enzyme has been characterised functionally in an ectomycorrhizal fungus, its role was symbiosis establishment rather than decomposition: a symbiosis-induced GH28 polygalacturonase in *Laccaria bicolor* localises to the hyphal front and is required for Hartig net formation, and silencing it impairs colonisation (Zhang et al., 2022). A copy that survives, and is under selection, may therefore be maintained for host colonisation rather than for nutrition. This is a warning about the inference we and others have been drawing, not a resolution of it.

*S4.5 Order-level composition of the ectomycorrhizal panel*

Supports the statement that comparative analyses across our panel carry an unmeasured amount of pseudoreplication.

The ectomycorrhizal habit is distributed across many independently derived fungal lineages (Tedersoo and Smith, 2013), and the 119 genomes we label ectomycorrhizal, after the three reassignments described above, are drawn from across the panel, which sounds like broad phylogenetic replication. It is not. Measured on the species tree itself rather than on taxonomic rank, those 119 tips fall into only eighteen clades, each wholly ectomycorrhizal and none nested inside another: the largest holds 51 of them, the largest two 79, the largest four 91, and only seven are isolated tips. Drawing 119 of the 182 tips at random returns a median of 83 such blocks with a 95% range of 72 to 92, so the observed value lies far outside what an unstructured labelling produces (10,000 permutations, $P < 10^{-4}$). We prefer this measure to a count of taxonomic orders because it is computed from the tree the analysis actually uses; we could not reconstruct a reliable order assignment for most of the panel and therefore do not report one. We have not counted how many times the ectomycorrhizal habit arose within this panel, and we do not know how many of these genomes represent independent acquisitions. A phylogenetic correction addresses shared ancestry in the response variable rather than non-independence in the trait's origins.

*S4.6 The measurement that would settle the question*

Supports the statement that the decisive evidence will not come from more genomes.

The comparison that would settle it is not another count and not, on our evidence, another branch-partitioned selection test. It is a direct measurement of activity (expression, secretion or enzymology) on the retained copies, in taxa chosen to span the phylogenetic breadth of the habit rather than to be convenient. On the block structure above, that means sampling outside the two clades that hold two-thirds of our ectomycorrhizal genomes. That work is expensive and it is not comparative genomics, which is presumably why it has not been done at the scale of the debate. We think the honest position is that the genomic literature, ours very much included, has been asked to carry an inferential load it was never able to bear.

#### SUPPLEMENTARY S5 — CALIBRATION DETAIL, SUPPORTING RESULTS 2 AND RESULTS 5 (THE FUNGAL NULL SETS, THE NUMBER OF LABELLED BRANCHES AND THE COST OF THE CORRECTION)

*S5.1 Phylogenetic clustering of the labelled sets*

Supports Results 2 ("The fungal panel").

Measured on the deposited labelled trees, the pre-audit ectomycorrhizal set is more clustered than the random draws on the same trees, both in mean pairwise distance among labelled tips and in mean distance to the nearest labelled neighbour (paired Wilcoxon $P = 4 \times 10^{-5}$ and $3 \times 10^{-11}$, 88 genes). Among the random draws, neither measure is associated with whether the gene was rejected (Spearman $\rho = -0.004$ and $+0.015$; $P = 0.97$ and $0.89$), nor with the estimate of $K$; and the clustering of that labelling falls inside the range the random draws already span in 86 of 88 genes.

Assembly and annotation quality, among-lineage rate variation and base-composition differences remain open as explanations of the residual gap, and the shift in $K$ between the two labellings is itself not significant (paired Wilcoxon $P = 0.063$).

**Null A and the starting-point setting.** Null A is an earlier random labelling drawn from the whole panel, so it included ectomycorrhizal tips, and it was also fitted from a single optimiser starting point rather than five, so it differs from the other two in two respects at once. Two observations bear on earlier versions of these numbers. Our original ectomycorrhizal-free null was not label-free, every one of its 88 draws including ectomycorrhizal tips (mean 8.6 of 21); it rejects in 30.3% on the same 76 genes, below both nulls above as a diluted labelling should, and is shown in Figure 2a for that reason only. And null B run at one optimiser starting point rather than five rejects 57.0% (49 of 86) instead of 51.2%, a rejection rate on presumed-null genes that moves by six percentage points when a convergence setting changes, and the first indication of the irreproducibility documented in full in the companion manuscript. The unpaired difference between the whole-set rates of null B and null C (82 and 81 genes) is 15.4 points, smaller than the 18.4-point paired difference on the 76 genes present in both.

*S5.2 Direction of rejections under the three misspecification options*

Supports Results 2 ("The fungal panel"). All three arms were run on the ectomycorrhizal-free null set (null C), one flag at a time, and each is tabulated against its own baseline; the arms are not comparable with one another.

| **Option** | ***n*** | **Baseline** | **With option** |
|---|---|---|---|
| --srv Yes | 71 | 35.2% | 32.4% |
| --multiple-hits Double | 63 | 41.3% | 34.9% |
| --error-sink Yes | 74 | 35.1% | **51.4%** |

Modelling multinucleotide substitution (--multiple-hits Double) reduces the rejection rate but reorganises the rejection calls rather than removing them, moving the balance from 19 relaxed and 7 intensified to 6 and 16.

Adding the alignment-error rate class (--error-sink Yes) raised the rejection rate from 35.1 to 51.4% while reducing both indicators of numerical difficulty: terminations with a negative likelihood-ratio statistic fell from 13 to 5, and returns of $K = 1.0000$ exactly from 15 to 4. The diagnostic indicators and the error rate therefore moved in opposite directions.

*S5.3 Gene-level overlap between implementations, and site-level FDR*

Supports Results 2 ("The fungal panel") and Discussion 2. All figures are for the ectomycorrhizal-free null set.

Of the 81 genes analysed by both programs, 13 were rejected by RELAX only, 38 by the two-ratio test only and 16 by both. Under Clade Model C the corresponding partition is 16, 24 and 13. Accommodating variation among sites thus reduced the excess by roughly one third.

The Contrast-FEL site-level figure is not comparable to the gene-level rates: a gene counts as rejected under the gene-level tests as a whole, whereas a gene here contains several hundred sites and will contain a significant one by chance almost always. Every gene did (88 of 88 carried at least one site at $P \leq 0.05$), which is what a nominal site-level rate predicts and not evidence of anything. Controlling the false discovery rate across sites within a gene brings the gene-level figure to 26 of 88 at $q \leq 0.05$ and 61 of 88 at the program's default of 0.20.

Running both stages of the program's own procedure leaves 71 of 88 genes, 80.7%, carrying at least one surviving site. That figure is likewise a summary rather than a calibrated quantity, and it is not on the axis of the gene-level rates above: at the two-stage site-level rate of 0.478% and a median of 594 sites per gene, independent sites would put about 94% of genes above zero, so 80.7 is what a conservative site-level rate produces once sites within a gene are correlated. We report it because the comparison invites the question, not because the three programs can be ranked on it.

Two counts appear in the two-stage arithmetic and the difference between them is worth stating, because the program's own output invites a wrong denominator. The asymptotic test called 5,603 of 61,059 sites significant at $P \leq 0.05$, while 6,001 sites carry a computed permutation $P$; the remaining 55,058 carry the sentinel value of -1 and were never permuted. The 398-site difference is degenerate. Every one of those sites has an asymptotic $P$ of exactly 1.000 and a permutation $P$ of exactly 1.000, and none of them contributes a discovery at either stage. Setting them aside, the two figures share one denominator: of the 5,603 asymptotically significant sites, 5,311 were removed by the permutation and 292 survived, 94.8 and 5.2%. Referring the 292 survivors to the 6,001 permuted sites instead returns 4.9%, which is the figure the program's summary reports and which we do not use.

The permutation stage has a resolution worth stating, since our own recommendations require it to be checked. Across all 6,001 permuted sites the permutation $P$ takes exactly twenty distinct values and every one of them is $1/m$ for an integer $m$ between one and twenty; no other value occurs. The counts fall away from the coarse end: 3,121 sites at 1, 947 at 1/2, 499 at 1/3, 278 at 1/4, 187 at 1/5, and so on to 20 at 1/19 and 292 at 1/20. At most twenty permutations therefore inform any one site, the smallest attainable $P$ is 0.05, and the 292 sites the two-stage procedure confirms are exactly the sites that reach that bound. None lies below it, and none could. This does not affect the 0.478% figure, which is an end-to-end rate rather than a $P$ value, but it does mean the stage cannot support a claim at any threshold corrected for multiple testing. The permutation refers the statistic to a reference the data generate rather than to $\chi^2$, though by permuting the labels rather than by reproducing the design. The two stages take the site-level rate from 1.8 times the nominal level to a tenth of it; Contrast-FEL is the only one of the three programs that does not reject above its nominal level, though the rates are measured on different units and we do not rank them.

*S5.4 The test-fraction series, step by step*

Each tested alignment is an *entry* (Methods M4), and an entry's *test fraction* is the proportion of the branches in its tree that carry the test label. Entry test fractions span 0.047 to 0.255 with a median of 0.089, while the null carries 21 of 99 branches, a fraction of 0.212. We therefore re-analysed the same null genes and trees with 3 to 29 labelled tips, the eight steps of the test-fraction series (Methods M5); sixty-nine genes returned an interpretable result at all eight.

The rejection rate rose with the size of the labelled set, from 33.3% at three labelled tips to 47.8% at twenty-nine, with a minimum of 31.9% at five (Figure 2b; Spearman $\rho = 0.77$; exact two-sided permutation $P = 0.033$), though not cleanly: three steps fell below their smaller neighbour, among them the 21-tip step whose null every enzyme entry was originally read against. The fifth percentile of the null distribution of *K* spanned a twofold range across the steps with no consistent direction (Figure 2c), so an observed *K* of 0.5 is called significant essentially always or essentially never depending on the step (Figure 2d).

Supports Results 5 ("The number of labelled branches"). The panel contains thirty non-ectomycorrhizal species, so twenty-nine is the largest draw that is not the exact complement of the ectomycorrhizal set. Sixty-nine genes returned an interpretable result at all eight steps and are tabulated on that common set.

| **Labelled tips** | **Test fraction** | **Rejected** | **Rate** | **Relaxed** | **Intensified** |
|---|---|---|---|---|---|
| 3 | 0.030 | 23 | 33.3% | 18 | 5 |
| 5 | 0.051 | 22 | 31.9% | 12 | 10 |
| 9 | 0.091 | 29 | 42.0% | 17 | 12 |
| 13 | 0.131 | 24 | 34.8% | 17 | 7 |
| 17 | 0.172 | 27 | 39.1% | 19 | 8 |
| 21 | 0.212 | 25 | 36.2% | 17 | 8 |
| 26 | 0.263 | 29 | 42.0% | 23 | 6 |
| 29 | 0.293 | 33 | 47.8% | 27 | 6 |

The composition of the rejections shifts alongside the rate, from ten intensified against twelve relaxed at five tips to six against twenty-seven at twenty-nine.

*Fifth percentile of the null distribution of K*

Across the eight steps the observed values are 0.333, 0.427, 0.664, 0.657, 0.615, 0.558, 0.417 and 0.621: a twofold range with no consistent direction. Each is estimated from 69 genes and is correspondingly imprecise. Resampling genes 5,000 times gives intervals that are wide and heavily overlapping, from [0.047, 0.445] at three labelled tips to [0.510, 0.728] at twenty-nine.

No single reversal is supported by the intervals, and the statistic we first used for the purpose (the proportion of resamples in which the eight percentiles emerged in order, which was zero) cannot distinguish the two hypotheses, since it resamples around a point estimate that is already disordered and would return zero under a monotone truth as well.

*Probability of a significant call*

An observed $K$ of 0.5 is called a significant relaxation against the seventeen-tip null in 100% of resamples, against the twenty-nine-tip null in 98%, against the nine- and thirteen-tip nulls in 86 each, against the twenty-one-tip null in 70, against the twenty-six-tip null in 29, against the five-tip null in 14, and against the three-tip null in 1%. The same observation is called significant essentially always or essentially never depending on which step supplies the null.

At the smallest test sets $K$ is barely identifiable at all: at five labelled tips its ninety-fifth percentile among null genes is 4.0, against roughly 1.16 in the middle of the range.

Between the two highest steps, at test fractions 0.263 and 0.293, the probability of a significant call for an observed $K$ of 0.5 moves from 29 to 98%, so a nearest-step assignment is least stable where the steps are closest together.

*S5.5 The CAZyme control*

The series holds alignment depth fixed at 50 or 51 single-copy sequences, while the enzyme entries are multicopy families of 43 to 902. Forty CAZyme families that are not plant-cell-wall-degrading, labelled by the same procedure at depths of 49 to 342 sequences with a median test fraction of 0.215, reject at 35.0% (14 of 40) against null C's 35.8% (Fisher exact $P$ = 1.00), and rejection is not associated with depth (Spearman $\rho$ = 0.07, $P$ = 0.69). **The gene-set control could not detect an effect even as large as the design effect**: forty families put a 95% Wilson interval of 22.1 to 50.5% around that 35.0%, which contains the whole range the design axis spans, 31.9% to 47.8% (Discussion 4).

Supports the closing sentences of Results 5 ("The number of labelled branches").

The CAZyme control rejects $K = 1$ in 14 of 40 families, 35.0%, against 35.8% for the ectomycorrhizal-free null (Fisher exact $P = 1.00$); the median $K$ is 1.0000 in both; negative likelihood-ratio statistics occur in 12.5 against 18.5% ($P = 0.45$), and exact boundary values $K = 1.0000$ in 22.5 against 22.2% ($P = 1.00$). Within the CAZyme control neither rejection nor $K$ tracks alignment depth (Spearman $\rho = 0.07$, $P = 0.69$ and $\rho = 0.22$, $P = 0.18$ against the number of sequences; rejection rates of 37.5, 33.3 and 33.3% below 100, between 100 and 200, and above 200 sequences).

The absence of a depth association bounds that dependence loosely rather than excluding it, and the failure of the control to detect an effect as large as the design effect is a weaker statement than the absence of any such effect.

The two deepest families in this control, at 342 and 285 sequences, both terminated at the exact boundary $K = 1.0000$ with a likelihood-ratio statistic of zero, after 10 and 9.6 hours of fitting respectively.

The composition of those rejections differs without reaching significance. Null C splits 21 relaxed to 8 intensified and the enzyme control 7 to 7 (Fisher exact $P = 0.18$), so the two agree on how often the test rejects while it cannot be told whether they agree on which way; every other null built here was relaxation-dominated, and a control that is not would be a result rather than a reassurance.

*S5.6 Sampling error in the bootstrap threshold, and the reference set*

Referring the observed statistic to a null distribution generated under the same design, rather than to $\chi^2$, is a standard remedy for a miscalibrated likelihood-ratio test, and two constructions of that reference appear below. A *parametric bootstrap* refers the statistic to replicates simulated under the fitted null model on the same labelled tree, as do Halabi et al. (2021) for a test of this family.

An *empirical null* refers it instead to real genes carrying the same labelling but no expected signal; nulls B and C and the steps of the test-fraction series are of that kind, and it is the construction we apply to the tested families below. The first is available for any gene, the second only where the analyst is willing to call a gene set null; what separates either from prior use is set out in Discussion 2. Contrast-FEL already applies a related device by default, a label-permutation test on significant sites (Discussion 2; Supplementary S5.3).

The simulated replicates permit an evaluation, since the generating $K$ is known. They are described in Results 5; two sets are used, one generated on the gene trees as inferred (the shallow set) and one whose branch lengths were rescaled until the mean pairwise identity matched that of the observed alignments (the divergence-matched set). As a parametric bootstrap reference we used the 94 true-null replicates of the shallow set, assessing each against the remaining 93. The type I error rate fell from 22.3 to 6.4%; under the two-ratio test it fell from 10.0 to 6.0%, and on the divergence-matched replicates from 17.5 to 6.2% (Figure 4a). These corrected rates are not evidence that the correction works: a replicate referred to others drawn from the same generator falls in their upper 5% about one time in twenty whatever the test does. What the simulations can measure is the cost of the correction. Whether a null simulated under the fitted model also calibrates an observed alignment, which was not drawn from that model, they cannot show; we test it on real genes in Results 3.

What the correction costs is modest on the shallow replicates and severe on the matched ones (Figure 4b). On the shallow set, detection at a true $K$ of 0.7 falls from 86.2 to 79.3% when the bootstrap threshold replaces the $\chi^2_1$ threshold, while detection at $K = 0.5$ is unchanged at 82.8% (29 replicates at each value). On the matched set the same substitution takes detection at $K = 0.5$ from 71.9 to 46.9% and at $K = 0.7$ from 65.6 to 9.4% (32 replicates at each), because the ninety-fifth percentile of the null statistic on the matched alignments is 152.6, against a $\chi^2_1$ critical value of 3.841. That percentile is a property of these simulated alignments rather than of the labelling design (Discussion 2).

Supports Results 5 ("What simulation can show about the correction").

All figures in this subsection are for the shallow simulated set, whose null statistic is an order of magnitude smaller than the divergence-matched set's; the 152.6 quoted in the main text is the matched set's ninety-fifth percentile and is not comparable with the values here. Across resamples of the shallow set the ninety-fifth percentile of the null statistic ranged from 2.45 to 8.64 when estimated from ten replicates and from 6.63 to 7.43 when estimated from eighty, with median values of 6.1 and 7.4 respectively. Both medians exceed the $\chi^2_1$ critical value of 3.841, but a reference distribution built from ten replicates can fall below it, in which case the bootstrap threshold is less conservative than the $\chi^2_1$ threshold.

Pooling across the five template genes used for simulation was at least as well calibrated as matching each replicate to its own template. The enzyme-family control provides independent support for that observation, rejecting at 35.0% against 35.8% for null C.

On the shallow replicates the bootstrap and $\chi^2_1$ thresholds gave 24 of 29 detections at $K = 0.5$ alike, and 23 against 25 of 29 at $K = 0.7$; with 29 replicates the smallest resolvable difference is 3.4 percentage points, so the $K = 0.5$ comparison is a true absence of loss, while the $K = 0.7$ gap of two detections is 6.9 percentage points, two steps of resolution. We previously reported that agreement as a loss too small to resolve; it was a threshold too small to bite, on alignments an order of magnitude shallower than the ones the correction is meant for.

Test fractions among the enzyme entries range from 0.047 to 0.255 (median 0.089), whereas the null against which they were assessed carries 0.212.

*S5.7 Entry-level step assignment*

Applying the correction to the tested families leaves two of 27 entries nominally significant rather than three: GH7 drops out (*P* = 0.088 under the nearest-step null against 0.049 under the shared 21-tip null; Results 6), and with 75 to 81 genes per step the smallest empirical *P* lies between 0.012 and 0.013, above the Bonferroni-corrected level for 27 tests (Discussion 2).

Supports Results 6 ("What the enzyme-family results do not show").

Each entry is referred to the step of the test-fraction series whose test fraction is nearest its own. The assignment and both readings are given here so that the recalculation can be checked entry by entry; bold marks a *P* at or below 0.05. Two entries are significant against their nearest step and three against the 21-tip null every entry was originally read against, the difference being GH7.

| **Entry** | **Test fraction** | ***K* (five starting points)** | **Nearest step (labelled tips)** | ***P* nearest step** | ***P* against the 21-tip null** |
|---|---|---|---|---|---|
| GH7 | 0.047 | 0.4660 | 5 | 0.0875 | **0.0488** |
| GH5_5 | 0.051 | 1.0263 | 5 | 0.7125 | 0.7927 |
| CE15 | 0.054 | 1.2014 | 5 | 0.8375 | 0.9390 |
| GH93 | 0.056 | 0.8781 | 5 | 0.2500 | 0.2439 |
| GH74 | 0.057 | 0.8904 | 5 | 0.2750 | 0.2561 |
| GH5_7 | 0.067 | 1.0047 | 5 | 0.7000 | 0.7073 |
| PL3 | 0.068 | 0.6383 | 5 | 0.1375 | 0.0732 |
| GH10 | 0.068 | 1.0000 | 5 | 0.6375 | 0.6585 |
| GH43 | 0.070 | 1.0000 | 5 | 0.6375 | 0.6585 |
| CE8 | 0.072 | 1.3460 | 9 | 0.8902 | 0.9756 |
| CE5 | 0.075 | 0.8357 | 9 | 0.2317 | 0.2195 |
| GH28 | 0.077 | 0.8694 | 9 | 0.2561 | 0.2317 |
| GH53 | 0.079 | 0.6675 | 9 | 0.0732 | 0.1098 |
| GH6 | 0.089 | 0.3824 | 9 | **0.0488** | **0.0244** |
| PL1 | 0.095 | 0.8777 | 9 | 0.2683 | 0.2439 |
| GH51 | 0.097 | 0.7732 | 9 | 0.1098 | 0.1463 |
| GH12 | 0.106 | 0.2846 | 9 | **0.0366** | **0.0244** |
| CE1 | 0.107 | 1.0084 | 9 | 0.5976 | 0.7317 |
| PL4 | 0.111 | 1.0085 | 9 | 0.5976 | 0.7317 |
| CE12 | 0.112 | 1.0000 | 13 | 0.6842 | 0.6585 |
| CE16 | 0.119 | 0.8942 | 13 | 0.2368 | 0.2683 |

| **Entry** | **Test fraction** | ***K* (five starting points)** | **Nearest step (labelled tips)** | ***P* nearest step** | ***P* against the 21-tip null** |
|---|---|---|---|---|---|
| GH5 | 0.142 | 0.9686 | 13 | 0.4605 | 0.3659 |
| GH5_9 | 0.186 | 0.8219 | 17 | 0.1585 | 0.2073 |
| GH5_50 | 0.205 | 0.7652 | 21 | 0.1341 | 0.1341 |
| GH5_12 | 0.235 | 0.8840 | 21 | 0.2439 | 0.2439 |
| GH5.sub | 0.252 | 0.9920 | 26 | 0.5769 | 0.4268 |
| GH5_15 | 0.255 | 0.6507 | 26 | 0.1282 | 0.0976 |

The mapping is a zero-order interpolation and is least stable where the steps are closest together: between the two highest steps the probability of a significant call for an observed *K* of 0.5 moves from 29 to 98% (S5.4). Entries whose test fraction falls between two steps are not interpolated, and we do not claim the procedure is exact, only that it is closer to the design of each entry than a single shared null is.

*S5.8 The fungal null sets in full*

Genes with no reason to be under relaxed selection specifically in ectomycorrhizal lineages, run through the identical pipeline, measure the test's rejection rate on presumed-null genes in this design directly (Figure 2). Two null sets on the same single-copy orthologues differ only in which 21 tips carry the test label: the tips labelled ectomycorrhizal when the codon analyses were fixed (null B; three of the twenty-one were reassigned by the later audit of Results 1, and Methods M5 explains why the labels were not changed), and 21 tips drawn at random from the non-ectomycorrhizal species (null C, the ectomycorrhizal-free null). A third, earlier labelling (null A; Figure 2a) is reported only as a check (Supplementary S5.1).

Labelling the 21 ectomycorrhizal tips on single-copy BUSCO orthologues rejects the null in 51.2% of genes (42 of 82; 60.0% of the 70 that returned an interpretable likelihood-ratio statistic). Replacing those labels, on the same alignments and trees, with 21 random non-ectomycorrhizal tips rejects in 35.8% (29 of 81; 43.9% of 66 with an interpretable statistic). Against a nominal rate of 5%, these are about ten and seven times too high. Whole-set rates keep uninterpretable genes in the denominator as non-rejections and are therefore lower bounds; comparisons between arms use the common interpretable set (Methods M5). On the 76 genes present in both arms the rates are 53.9 against 35.5%, a difference of 18.4 percentage points, with 28 genes rejected under the ectomycorrhizal labelling alone and 14 under the random labelling alone (McNemar exact $P = 0.044$).

Random labels isolate what the branch-partitioning scheme contributes on its own, and roughly two-thirds of the excess is present before any biological signal is available to produce it. The residual gap between the arms is not established as ectomycorrhizal-specific by these data: the random labelling is not structure-preserving (Saputra et al., 2021), though clustering does not predict rejection among the draws (Supplementary S5.1), and three of the twenty-one labelled tips are not ectomycorrhizal, a dilution that narrows the gap rather than producing it. Null B run at one optimiser starting point rather than five rejects 57.0% (49 of 86) instead of 51.2%, a first indication of the irreproducibility documented in the companion manuscript (Kim and Shin, 2026b, Results 1).

In PAML, on the same null set, the two-ratio branch model rejected 58 of 88 genes (65.9%) at $\alpha = 0.05$, and Clade Model C, which allows one of three site classes to differ between branch sets (against M2a_rel; Weadick and Chang, 2012), 40 of 88 (45.5%). On the 81 genes shared with the RELAX run the three reject 54 (66.7%), 37 (45.7%) and 29 (35.8%): accommodating variation among sites left a rejection rate nine times the nominal level. Contrast-FEL (Kosakovsky Pond et al., 2021), which tests the same null site by site without a $K$ parameter and with synonymous rate variation on, rejected at 9.18% of 61,059 sites at its asymptotic stage. The site-level figure is not comparable to the gene-level rates, and the programs do not fail on the same genes (Supplementary S5.3).

#### SUPPLEMENTARY S6 — SUPPORTING DETAIL FOR RESULTS 5 (WHAT THE TEST DOES WHEN THE ANSWER IS KNOWN)

*S6.1 Rescaling the simulation to the divergence of the data*

Supports the statement that the first construction was rebuilt so that each simulated alignment reached the mean pairwise identity of its own template.

The branch-length scale was searched separately at each generating value of $K$, because $\omega_{\text{test}} = \omega_{\text{ref}}^{K}$ accelerates the labelled branches whenever $\omega_{\text{ref}}$ is below one, and a scale fitted at $K = 1$ therefore overshoots elsewhere. The scale factors this required (12.6 to 53.2 at $K = 1$, and roughly half that at $K = 0.5$) are themselves a measure of how far the first construction was from the data.

The rebuilt set reaches realised identities of 0.557 to 0.561 against per-template targets of 0.500 to 0.593.

Under the shallow construction the ranges of total tree length do not overlap: 30.3 to 215.3 on the null orthologues against 3.4 to 17.1 on the simulated replicates. The branch lengths were taken from a gene tree estimated under a nucleotide model and passed to the simulator, which interprets branch lengths under the codon model it is generating from; the fitting program, which re-estimates branch lengths, is insensitive to this, but the simulator is not.

**Replicate counts and completion.** The rebuild kept twenty replicates per template at $K = 1$ and raised the count at each relaxed value from six to eight. Completion was equalised: an earlier pass truncated 22 of 352 fits on observed data against 69 and 70 of 400 on the tip- and internal-labelled simulated arms; re-running all 161 under a twenty-four-hour limit brought the three sets to full completion. In the divergence-matched set the same pass left two of 320 clade-model fits incomplete, 0.6%.

*S6.2 A withdrawn decomposition of the observed rejection rate*

Supports the statement that the simulated rate bounds what the test does when the answer is known but does not partition the observed rate.

An earlier version of this work read the simulated and observed rates as a decomposition, attributing about 22 percentage points to the test itself and about 13 to whatever real alignments carry in addition. We withdraw that reading, for the reasons that follow. The two rates rest on different sets, 94 simulated replicates against 81 observed genes, and the shallow simulated alignments are about an order of magnitude less diverged; we do not claim the simulation accounts for the observed rate.

*S6.3 Non-convergence in the generating model*

Supports the statement that the $\omega$ distributions were inherited from RELAX null fits that had in part not converged.

Across the 81 null genes the largest $\omega$ class exceeds 100 in seventeen genes and sits at the optimiser bound in three.

*S6.4 Rejection rate by template gene*

Supports the statement that the inflation of the error rate is not the property of a single pathological alignment.

In the primary configuration (tip labels, five starting points) rejection rates varied across template genes from 11.1 to 36.8%.

*S6.5 Single-ω cross-check on the direction of the RELAX bias*

Supports the statement that the errors in the primary configuration are asymmetric.

A single-$\omega$ fit gives the same picture independently of $K$: in the primary configuration (tip labels, five starting points) the median ratio of test to reference $\omega$ was 0.957.

*S6.6 Composition and paired transitions under internal labelling*

Supports the statement that moving the test label from the tips to an internal branch changes both the rate and the sign of the error.

On the shallow set, across the 89 replicates fitted under both labellings, the composition of rejections went from 19 relaxations against one intensification under terminal labels to one against eleven under internal labels.

On the divergence-matched set, across the 80 replicates fitted under both labellings, 22 replicates were rejected under internal labels alone against eight the other way (McNemar exact $P = 0.016$).

In the two-ratio comparison on the shallow set, the RELAX internal-label rate of 13.7% is over the 95 replicates returning a statistic under internal labels; over the 89 fitted under both labellings it is 13.5%.

*S6.7 Sign of the two-ratio bias under both labellings*

Supports the statement that the sign of the two-ratio bias, unlike that of the RELAX bias, is unaffected by the labelling.

Foreground $\omega$ was estimated below background $\omega$ in 74 of 100 tip-labelled replicates and 84 of 100 internal-labelled replicates, with median differences of -0.008 and -0.017 against a true difference of zero.

*S6.8 Relation between K and the two-ratio difference*

Supports the statement that the two statistics are less closely related than their joint miscalibration suggests.

Across the 94 tip-labelled replicates analysed by both programs the relationship is not merely weak but runs counter to the mapping the RELAX parameterisation implies: since $\omega_{\text{test}} = \omega_{\text{ref}}^{K}$ and every background estimate here lies below one (0.036 to 0.341), a $K$ below one should raise foreground $\omega$ above background, yet only 28 of the 94 replicates are concordant with that expectation (exact binomial $P = 1.1 \times 10^{-4}$). This follows from the two formulations rather than indicating a fault in either: $K$ compresses a three-class distribution of $\omega$ towards one, whereas the two-ratio model fits a single $\omega$ to each branch set, and the mean of a compressed distribution may move in either direction. Agreement between the two programs on the rejection rate therefore reflects a shared property of the design rather than agreement about the quantity estimated.

*S6.9 Optimiser behaviour of the clade model on the divergence-matched set*

Supports the statement that a residual factor of 1.68 on interpretable fits (2.8 if the two arms' unequal negative-likelihood-ratio fractions are ignored, which Results 5 declines to do) separates the divergence-matched simulation from the observed null genes.

Terminations with a negative likelihood-ratio statistic, which are impossible under convergence, occur in 41.2% of clade-model fits on the divergence-matched set, against 8.0 and 4.0% on the shallow arms and 2.3% on the observed data. Agreement between the two starting values falls in step, from 27 and 30% under M2a_rel and 34 and 40% under Clade Model C on the shallow arms, to 5.0 and 11.2% on the matched set. Deep alignments fitted under a mixture over sites are where this optimiser fails most often, and that combination is more extreme in the simulation than in the data it was built to imitate.

Both headline rates are therefore depressed lower bounds, and depressed to very different degrees, the depression being eighteen times greater on the matched set. Restricted to interpretable fits the rates are 40 of 86 on the observed genes and 13 of 47 on the divergence-matched set, 46.5 against 27.7%.

*S6.10 The 78-gene simulated-set replication of the two-ratio labelling contrast*

Supports the 78-gene figures in Results 5. Genes were classified by their fitted RELAX null $\omega$ distribution before simulation; three boundary-stratum genes whose fitted class sat at the optimiser ceiling of $10^{10}$ could not be simulated (the generating rate matrix cannot be constructed) and are excluded throughout.

| **Stratum (genes)** | **Terminal labels** | **Internal labels** | **Ratio** |
|---|---|---|---|
| Moderate: all $\omega$ < 100, one class > 1 (57) | 25.4% | 34.2% | 1.3 |
| No fitted heterogeneity: highest class = 1 (7) | 14.3% | 21.4% | 1.5 |
| Boundary: any class ≥ 100 (14) | 10.7% | 66.1% | 6.2 |
| All 78 genes, gene-averaged (4 replicates each) | 21.8% (68/312) | 38.8% (121/312) | 1.8 |

Rates are gene-averaged throughout, each gene weighted equally. Every gene contributes exactly four replicates, so pooling the 312 cells in the final row and averaging the 78 per-gene rates give the same value; the final row is the three strata combined. The analysis plan fixed before the batch ran made the moderate stratum the primary comparison and directed that the other two strata be reported separately from it; the combined row is therefore a departure from that rule, recorded as an amendment and given here because the main text quotes it. It should be read with the stratum rows above it, which is where the effect is actually located. The paired McNemar test over the 312 shared alignments gives 98 replicates rejected under internal labels only against 45 under terminal labels only (exact $P = 1.1 \times 10^{-5}$); restricted to the moderate stratum, 58 against 38 ($P = 0.052$). The five original templates reproduce their earlier behaviour on the shared alignments (0 of 20 terminal-labelled replicates rejected against 13 of 20 internal), and every negative likelihood-ratio statistic and every starting-point disagreement in either arm falls in the boundary stratum. The template in that stratum carries a fitted class at $\omega$ = 145, so the difference between the five-template and 78-gene results lies in the template genes, not the test; and because the boundary stratum holds every convergence failure, the moderate-stratum rates are free of the failures documented in the companion manuscript. Stratum definitions and the gene-averaged summary were fixed in that plan before the batch ran, whereas the paired McNemar test was not specified in it; the batch regenerates from recorded per-replicate seeds.

*S6.11 The simulated sets and the RELAX results in full*

*Design*

The parameters of the simulation were read from the data rather than chosen. For each of five template genes spanning 239 to 1,132 codons, the three-class $\omega$ distribution and its weights came from that gene's own RELAX null fit, the transition/transversion ratio from its GTR fit, the alignment length from its alignment, and the topology, branch lengths and {Test} labels from its own gene tree. Only $K$ was set by us: twenty replicates at $K$ = 1 and six each at $K$ = 0.7 and $K$ = 0.5, giving 160 alignments, re-analysed with the same RELAX invocation used throughout.

That first construction was far shallower than the data it stood in for (median total tree length 12.1 against 116.3 on the null orthologues, fitting the same model to both). The cause, identified only later, is in part a mismatch of units: the gene-tree branch lengths are expected substitutions per nucleotide site, whereas the codon simulator reads a branch length as substitutions per codon. The set was therefore rebuilt, the branch-length scale of each template being found by bisection so that the simulated alignment reached the mean pairwise identity of that template's own alignment. One template could not reach its target and was dropped, leaving 80 replicates at $K$ =

1 and 32 each at $K = 0.7$ and $K = 0.5$ (Supplementary S6.1). Every figure below is labelled with the set it comes from. Two limitations remain: the $\omega$ distributions were inherited from RELAX null fits that had in part not converged (Supplementary S6.3), and the templates were selected because they had returned $K \approx 1$ in the observed analysis, which is selection on the outcome of the test under study.

Recovering the divergence of the observed alignments also recovered the pathology that the shallow set could not produce: returns of $K = 1.0000$ exactly rose from 1.1 to 28.8% of replicates, against 22.2% on the observed null genes. The shallow set was, in this sense, a negative control that could not reproduce the failure it was built to characterise.

The labelling contrasts below rest on codeml rather than on RELAX. Refitted at a second optimiser setting, none of 27 enzyme entries reproduced its own RELAX log-likelihood, and 46 of 410 fits ended in a negative likelihood-ratio statistic, which convergence makes impossible, and every one of those runs terminated normally (Kim and Shin, 2026b, Results 1 and 2); a RELAX rejection rate therefore carries optimiser variance on top of whatever the design contributes. The two-ratio model returns the same fit on repetition (Kim and Shin, 2026b, Results 1), so a difference between its two labellings on the same alignment is not optimiser noise.

*The nominal error rate is not attained*

Under the primary configuration (tip labels, five starting points; Figure S1), 21 of 94 true-null replicates were rejected at $P < 0.05$, or 22.3%, four and a half times the nominal rate; the one-starting-point arm rejected 16 of 100, or 16.0%. These alignments contain no alignment error, no rate variation across sites, no multinucleotide substitution and no branch-to-branch heterogeneity, and were generated under the model being fitted, so the excess cannot be attributed to any property of real sequence data. Every template gene rejects above nominal (Supplementary S6.4). The simulated rate bounds what the test does when the answer is known; it does not partition the observed rate of 35.8% (Supplementary S6.2). The matched set rejects less, 14 of 80 replicates or 17.5%, against 29 of 81 observed genes (Fisher exact $P$ = 0.012), while the shallow set's 22.3% is not distinguishable from the observed rate ($P$ = 0.065): recovering the divergence of the real alignments recovers the pathology documented in the companion manuscript (Results 2) without recovering the rejection rate.

*The errors are not symmetric, and their direction is not predictable*

Of the 21 rejections in the primary configuration, 20 were in the direction of relaxation; the median estimate of $K$ was 0.9409 against a true value of one, and 78 of 94 replicates fell below one (two-sided binomial $P = 5.8 \times 10^{-11}$; Supplementary S6.5). Moving the test label from the tips to an internal branch changes both the rate and the sign. On the shallow set, across 89 replicates fitted under both labellings, the rejection rate fell from 22.5 to 13.5% and the fraction of estimates below one from 83.1 to 20.2%. On the divergence-matched set the contrast reversed: across 80 replicates, terminal labelling rejected 17.5% and internal labelling 35.0% (McNemar exact $P$ = 0.016), with 31.2 against 43.8% of estimates below one and a median $K$ of exactly 1.0000 under both. On shallow replicates terminal labelling produced relaxation in 19 of 20 rejections; at the divergence of the observed alignments it produced intensification in 10 of 14. An analyst who knows only where the labels sit knows neither how often the test will reject nor on which side of $K = 1$ it will do so.

*An independent implementation responds differently to the same change of design*

We repeated both labellings in PAML with the two-ratio branch model on the same five-template alignments and labelled trees; all 800 fits completed. Under tip labels the two-ratio test rejected

10.0% of the true-null replicates, twice the nominal level (one-sided exact binomial $P = 0.028$) but less than half the RELAX rate. Moving the label to internal branches reversed the ordering: the two-ratio rate rose to 47.0% while that of RELAX fell from 22.5 to 13.5% on the 89 replicates fitted under both labellings, with 41 replicates rejected under internal labels but not tip labels against four the other way ($P = 9.3 \times 10^{-9}$; Supplementary S6.6). Both implementations are miscalibrated and respond to the same change of design in opposite directions; no labelling scheme yields nominal behaviour in both. These five-template figures generalise in neither direction: on the replicates matched to the observed alignments the two-ratio test under tip labels rejects 37 of 80 (46.2%) against 17.5% for RELAX, reversing which is the more inflated.

The sign of the two-ratio bias is unaffected by the labelling, foreground $\omega$ being estimated below background $\omega$ under both (Supplementary S6.7), so the interpretation that a tip-labelled design generates apparent relaxation applies to RELAX specifically. Nor are the two statistics closely related: across the 94 tip-labelled replicates analysed by both programs, $K$ and the two-ratio difference between foreground and background $\omega$ show no rank association (Spearman $\rho = 0.04$, $P = 0.69$; Supplementary S6.8). These alignments were simulated under the three-class $\omega$ mixture that RELAX fits, which favours RELAX over a single-$\omega$ model; on the observed null genes the ordering is reversed (Results 2).

Clade Model C, which also accommodates variation among sites (tested against M2a_rel; Weadick and Chang, 2012), rejected 3.0% of tip-labelled and 11.0% of internal-labelled shallow replicates and 16.2% of the divergence-matched set, against 45.5% of the observed null genes. Negative likelihood-ratio terminations, which cannot reject, are eighteen times more frequent on the matched set; restricted to interpretable fits the residual factor is 1.68 rather than 2.8 (Supplementary S6.9). With truncation removed as an explanation by re-running the truncated fits to completion (two of 320 remained incomplete on the divergence-matched set; Supplementary S6.1), and depth by the divergence-matched set, it is that residual, not the whole of the 45.5%, that implicates a property of real coding sequence the simulation does not reproduce; alignment error, gene-tree misspecification and among-lineage rate variation remain the candidates, and we have not distinguished among them.

*The magnitude of the labelling contrast tracks the fitted ω configuration*

Extended to the full null C gene set (Methods M5; 312 alignments from 78 genes, each fitted under both labellings), the direction replicates, 98 replicates being rejected under internal labels only against 45 under tip labels only (McNemar exact $P = 1.1 \times 10^{-5}$; 21.8% under tip labels, 68 of 312, against 38.8%, a pooled comparison recorded as an amendment to the plan), but the 4.7-fold separation of the five-template rates does not; the five templates themselves reproduce their earlier behaviour, none of 20 tip-labelled replicates rejecting against 13 of 20 internal. In the moderate stratum, primary in the plan, the rates are 25.4 and 34.2% (58 against 38 discordant replicates, $P = 0.052$); the contrast concentrates in the boundary stratum, at 10.7 against 66.1%, which holds one of the five templates and every negative likelihood-ratio statistic and starting-point disagreement in either arm (Supplementary S6.10). The tip-labelled rate of 10.0% above is therefore a five-template value; on the 78 gene-derived parameter sets even the least inflated labelling, terminal, rejected at 21.8%, more than four times the nominal level, and the size of the labelling effect is a property of the fitted $\omega$ configuration, largest exactly where the generating fit is least trustworthy.

*Detection is adequate before the correction and not after it; the estimates are poor throughout*

Genuine relaxation was detected at 82.8% for $K = 0.5$ and 86.2% for $K = 0.7$ on the shallow set, lower for the larger true effect and therefore inconsistent with the expected behaviour of the test.

On the divergence-matched set the asymptotic ($\chi^2$) figures are 71.9 and 65.6%, in the expected order. The estimates are worse than the calls. At five starting points on the shallow set, a true *K* of 0.7 gives recovered values spanning 0.014 to 1.025 and a true *K* of 1.0 gives 0.411 to 2.411, wide enough to contain every effect size the observed analysis reported. This justifies reporting profile intervals rather than point estimates.

*S6.12 Figure S1*

Supports Results 5.

**Figure S1.** Estimates of *K* when the generating value is known. Both panels show the shallow simulated set, not the divergence-matched set. (a) RELAX estimates of the selection-intensity parameter *K* under terminal labels with five optimiser starting points, against the generating value. Each point is one replicate, red where $P < 0.05$; teal bars mark the true *K*, the dashed line $K = 1$. $n = 29$ at $K = 0.5$, 29 at $K = 0.7$ and 94 at $K = 1.0$; annotated medians 0.620, 0.734 and 0.941, and 21 of the 94 true-null replicates were rejected. (b) Percentage rejected at true $K = 1$ among the 89 replicates from (a) that were also fitted under internal labels, at five starting points, split by direction: tip labels 22.5% (19 relaxed, $K < 1$; 1 intensified, $K \geq 1$), internal branches 13.5% (1 relaxed, 11 intensified). Dashed line, nominal 5%. **Both features of (b) reverse once the replicates are matched to the divergence of the observed alignments, and this figure does not show that set**: there, across the 80 replicates fitted under both labellings, terminal labelling rejects 17.5% against internal labelling's 35.0% (McNemar exact $P = 0.016$), and terminal labelling's rejections fall mostly on the intensified side, 10 of 14 (Results 5).

#### SUPPLEMENTARY S7 — SUPPORTING DETAIL FOR THE TWO PANELS REGISTERED IN ADVANCE (RESULTS 2 TO 4 AND RESULTS 5)

*S7.1 The two panels*

Supports Results 2 ("Two further panels, registered in advance").

The first is 50 yeasts of the order Pichiales with the standard nuclear genetic code, where the trait is failure to grow on glycerol (Opulente et al., 2024); the second is 50 lactobacilli from the smallest clade of the genome taxonomy tree holding enough host-adapted and free-living species, where the trait is host adaptation against free-living and nomadic lifestyles (Duar et al., 2017).

Four sets of amendments made after the lock, each recorded before the fits it affects, are listed with their reasons in Supplementary S3; three were made after the first two yeast verdicts were known, and two selection choices after the resulting trait had been seen.

Both panels are about half as deep as the fungal one (median one-ratio tree length 52.5 and 61.4 against 116.3). The registered threshold for narrowing the clade, twice the fungal value, was not reached in either. The placement of the trait also differs. The 18 host-adapted lactobacilli do not form a single clade on any of the 80 gene trees, falling mostly into three groups.

*S7.2 The second and third predictions in more detail*

Supports Results 5 ("Moving the label from terminal to internal branches" and "A reference built under one labelling does not calibrate another").

The size of the terminal-to-internal shift in rejection rate under the second prediction differs among the three panels, about 1.3-fold in the fungi, 2.5-fold in yeast and 1.75-fold in lactobacilli, and we do not interpret that difference.

The third prediction asks the question that pooled nulls had left open. The reverse referral, terminal-labelled replicates against the internal-labelled reference, rejected 5.2, 1.7 and 2.4%, and referring each replicate to its own design returned 4.0% in every panel, as the construction requires. The excess is concentrated: seven of the 20 fungal genes, three of the 20 yeast genes and nine of the 20 lactobacillus genes had rejection rates of 5% or less, and a few genes carry most of the mean.

*S7.3 The fourth prediction: incomplete genes and stratum stability*

Supports Results 3 ("As registered").

The reference removed about a quarter of the excess over 5% in yeast and an eighth in lactobacilli. The median likelihood-ratio statistics quoted in the main text are medians over genes in yeast and over all replicates in lactobacilli.

Twelve yeast genes and 13 lactobacillus genes have no generator, because the RELAX null fit ended in an optimiser error (ten in each panel) or returned a class at or near the optimiser ceiling (at $10^{10}$ in one yeast and three lactobacillus genes, and at $6.1 \times 10^{7}$ in one yeast gene, where the generator failed), and they are counted as incomplete rather than removed; the share of planned fits left incomplete, 15.0 and 16.2%, exceeds the registered 10% flag in the stages that depend on the generator, and the verdicts reported in Results 3 are computed on the completed fits as registered. Scored over all 80 genes with those without a generator counted as not rejected, the rates are 36 of 80 in yeast, 45.0% (33.8–56.5), and 51 of 80 in lactobacilli, 63.8% (52.2–74.2), and the conclusion is unchanged. One further caution applies to the stratum assignments. Refitting the yeast genes with a different seed changed the stratum of 20 of 65, the instability documented for RELAX in the companion manuscript (Kim and Shin, 2026b); any $K = 1$ generator remains a valid null, so the direction of the second and third results is unaffected, but the per-stratum rates carry that instability.

*S7.4 The exploratory follow-up in more detail*

Supports Results 3 ("An exploratory follow-up: a generator at the depth of the data, and a branch-site test").

A second generator, the RELAX null model with its branch lengths in the correct unit, was planned but dropped before the full run began: those branch lengths are inflated by the large-$\omega$ classes of the RELAX fits, to a median of 11.4 times the one-ratio tree length in yeast and 32.9 times in lactobacilli.

In the branch-site analysis the refit used five starting values per model in all. The one rejection, which had arisen from a null fit stopped at a boundary, disappeared. Even with five starting values the fits of the alternative model differed by up to 119 log-likelihood units on one yeast gene, so an optimum missed by all five cannot be excluded. No gene was rejected at the 50:50 mixture critical value of 2.71 either.

The power of the branch-site test was measured on ten genes per panel drawn by a fixed seed from the 80, under a plan and a calibration (one gene, three replicates) fixed by checksum before any power fit. Each gene was simulated with pyvolve under codeml's branch-site model A with the parameters of its best branch-site null fit to the real alignment, its codon frequencies, its codon count and its missing data, with the random terminal labels of the real analysis as foreground; branch lengths followed codeml's definition for model A on each branch type, which a calibration on 3,000-codon replicates recovered to within 3% (tree length, foreground length, $\kappa$, $\omega_0$) where the alternative scaling inflated the foreground length 2.8-fold. Four cells were run, ten replicates per gene each: no effect (the null fit unchanged), $\omega_2$ = 3 or 8 at 10% of sites, and $\omega_2$ = 20 at 5%. Each replicate was fitted with the control file of the real genes, from four null and three alternative starting values, refitted from the other fit's branch lengths and $\kappa$ where the best alternative ended more than 0.001 below the best null or the test rejected, and rejected at $\chi^2_1 > 3.841$. Power was averaged over genes with a percentile bootstrap over genes. With no effect, 5 of 100 yeast replicates (95% bootstrap interval 1.0–9.0%) and 3 of 100 lactobacillus replicates

(0.0–7.0%) were rejected, so the size is not shown to exceed 5%, though it is not shown to be controlled at 5% either. With an effect, 100 of 100 yeast replicates were rejected in each cell, and 95, 96 and 97 of 100 lactobacillus replicates (bootstrap lower bounds 85, 88 and 91%); pooled, 97.5, 98.0 and 98.5%. Every lactobacillus miss came from one gene with the longest tree in either panel, on which most first-stage starts stopped at about half the generating tree length; in those replicates the null escaped on refitting and the alternative did not, so the misses are failures of the optimiser and every evaluable replicate of that gene was rejected. Counting them as not evaluable, a post hoc reading, gives 100% in each cell. In two yeast null replicates the alternative collapsed below an interior null point; counting both as rejected gives a yeast size of 7.0% (3.0–11.0%) and a pooled size of 5.0% (2.5–8.0%), with no lower bound above 5%. With power measured, the zero on the 80 real genes of each panel bounds the share of them carrying an effect of a cell's size on every labelled branch at 3.7% in yeast and 4.0–4.3% in lactobacilli, and at 1.9–2.0% pooled (one-sided, joint coverage at least about 92.5%). The bound holds only for these genes, these labels and these effects, on data drawn from the fitted model, so it is optimistic for real data; the gene-level bootstrap rests on ten genes, and the exact binomial lower bound for ten of ten high-power genes, 0.74 (0.86 for twenty), would widen it to 5.0% per panel and about 2.2% pooled. Twenty-one of 6,227 fits stopped at the 24-hour limit; each was a single starting value and none could have changed a decision.

In the gap check the chosen genes had gaps at 5.1 to 12.3% of characters in yeast and 1.0 to 2.7% in lactobacilli. Against the replicates as first run, 6 of 10 yeast and 8 of 10 lactobacillus genes were rejected; against the same replicates with the real gaps, the same 6 and 8, with no gene rejected under one reference only (exact McNemar $P$ = 1.000). Imposing the gaps moved the replicates' one-ratio tree length by less than 0.5% on every gene. The check can move only genes near the threshold: 13 of the 14 rejected genes had no replicate reaching the real statistic under either reference. It treats a gap as missing data, as codeml does under cleandata = 0, and so does not reproduce alignment error next to gaps.

In the per-branch analysis the instrument check passed after its revision: the median ratios of one-ratio tree length and $\omega$ were 1.111 and 1.068 for G3 in yeast and 1.152 and 1.007 in lactobacilli, 1.026 and 0.979 for G3n in yeast and 0.998 and 0.980 in lactobacilli, and between 0.97 and 1.06 for the check's single-$\omega$ generator, and the median round-trip slope of G3 was 0.945 and 0.894. All 240 HyPhy fits and all 12,800 replicate fits completed. Every one of the 40 real alignments departed from a single $\omega$ more than its five comparators; the real statistic was a median 6.48 times the median comparator in yeast (range 2.66–19.33) and 3.82 times in lactobacilli (1.54–15.88). Against G3, 0 of 20 yeast genes and 1 of 20 lactobacillus genes were rejected (one-sided 95% upper bounds 13.9 and 21.6%); against G3n, 13 and 15; against the first analysis's replicates, 14 and 18. One gene was rejected under G3 only and 28 under G3n only. The single G3 rejection is marginal: one of its 40 replicates reached the real statistic (8.32 against 7.34). Taking a one-ratio replicate's own random-label statistic as if it were real, none of the 40 genes was rejected against G3n. The G3 replicates were rejected at $\chi^2_1$ in 72.4 and 81.2% of fits under fresh labels, close to the real genes' 14 and 18 of 20, and the G3n replicates in 19.2 and 18.9%. In the fitted pattern of every real gene the median $\log_{10} \omega$ of internal branches exceeded that of terminal branches, by a median 0.53 (range 0.19–0.78), against 0.03 in the per-branch fits to one-ratio replicates; because random labels fall on terminal branches only, this contrast alone sets a random terminal set apart from the rest, and the analysis does not separate it from variation among terminal lineages. That G3 is wider than the truth follows from its construction; the data do not measure by how much.

*S7.5 Positive controls, reported without a verdict*

At $K = 0.5$ and 0.7 on the test branches, four replicates per gene and value under terminal labels, the two-ratio test detected the departure in 73.2 and 54.4% of 272 yeast replicates at $\chi^2_1$ and in 65.3 and 42.9% of 268 lactobacillus replicates. Referred instead to the 60 $K = 1$ replicates of the same gene, detection fell to 61.4 and 42.3% in yeast and to 56.0 and 28.4% in lactobacilli. These rates are measured on the shallow simulated alignments of the fourth prediction, a sixth to an eighth as diverged as the real ones, and are not an estimate of power on real data.

*S7.6 Label position and length under correct specification (exploratory)*

The plan (newpanel/P4L/PLAN.md), its scripts, the label sets and a pre-run review were fixed by checksum before any fit. Replicates r001 to r020 of the depth-matched one-ratio generator (Results 3) were fitted under four label sets of $k$ branches ($k = 19$ in yeast, 18 in lactobacilli), each with the two-ratio model from $\omega = 0.5$ and 1.5 against the one-ratio fits of the same replicate (likelihood-ratio test on one degree of freedom, rejection at 3.841): T_rand, the random terminal labels of the registered analysis; T_long, the $k$ longest terminal branches; I_long, the $k$ longest internal branches; and I_match, internal branches whose summed generating length lies within 5% of that of T_rand, chosen per gene (62 yeast and 76 lactobacillus genes reached that band; for the others I_match equals I_long where even the longest internal branches fall short (15 yeast and 2 lactobacillus genes) and is otherwise the closest draw (3 and 2), and the primary contrast is restricted to the in-band genes). Intervals are gene-clustered bootstrap percentiles (10,000 resamples); a contrast is *absent* when its 95% bootstrap interval includes 0 and its 90% bootstrap interval lies within ±5 percentage points, *present-small* when its 95% bootstrap interval excludes 0 and the point estimate is below 5. All 19,200 new fits completed; no flag was triggered (incomplete 0%; replicates whose two starting values differ by more than 0.01 log-likelihood units at most 0.1% in any arm).

| Panel | Arm | Rejected | % | 95% CI (Clopper–Pearson) |
|---|---|---|---|---|
| Yeast | T_rand | 90 / 1,600 | 5.6 | 4.5–6.9 |
| Yeast | T_long | 122 / 1,600 | 7.6 | 6.4–9.0 |
| Yeast | I_long | 101 / 1,600 | 6.3 | 5.2–7.6 |
| Yeast | I_match | 95 / 1,600 | 5.9 | 4.8–7.2 |
| Lactobacilli | T_rand | 114 / 1,600 | 7.1 | 5.9–8.5 |
| Lactobacilli | T_long | 122 / 1,600 | 7.6 | 6.4–9.0 |
| Lactobacilli | I_long | 125 / 1,600 | 7.8 | 6.5–9.2 |
| Lactobacilli | I_match | 95 / 1,600 | 5.9 | 4.8–7.2 |

| Panel | Contrast | Pairs (genes) | Difference, pp | 95% | 90% | State | McNemar |
|---|---|---|---|---|---|---|---|
| Yeast | I_match - T_rand (in band) | 1,240 (62) | +0.2 | -1.5 to 1.9 | -1.2 to 1.7 | absent | 65 / 62 |
| Yeast | T_long - T_rand | 1,600 (80) | +2.0 | 0.3 to 3.8 | 0.6 to 3.5 | present-small | 116 / 84 |
| Yeast | I_long - I_match (in band) | 1,240 (62) | +0.6 | -1.0 to 2.3 | -0.7 to 2.1 | absent | 59 / 51 |
| Yeast | I_long - T_rand | 1,600 (80) | +0.7 | -0.9 to 2.3 | -0.6 to 2.1 | absent | 91 / 80 |
| Lactobacilli | I_match - T_rand (in band) | 1,520 (76) | -1.2 | -2.9 to 0.5 | -2.7 to 0.1 | absent | 77 / 96 |
| Lactobacilli | T_long - T_rand | 1,600 (80) | +0.5 | -1.2 to 2.2 | -0.9 to 1.9 | absent | 101 / 93 |
| Lactobacilli | I_long - I_match (in band) | 1,520 (76) | +1.8 | 0.2 to 3.5 | 0.5 to 3.2 | present-small | 94 / 66 |
| Lactobacilli | I_long - T_rand | 1,600 (80) | +0.7 | -1.2 to 2.6 | -0.9 to 2.2 | absent | 110 / 99 |

The size of T_rand on r001 to r060 is 261 of 4,800 (5.4%; bootstrap 4.9–6.0%) in yeast and 327 of 4,800 (6.8%; 6.2–7.5%) in lactobacilli. The lactobacillus upper bound equals the 7.5% limit of the range the plan calls nominal (360 of 4,800); under other bootstrap seeds it lies just above it. The plan reads I_match - T_rand with T_long - T_rand and requires both panels to fall in the same row of its outcome table; yeast falls in the row with the first absent and the second present-small, lactobacilli in the row with both absent, so the panels disagree and no clause is written.

I_long - T_rand and I_long - I_match carry no rule. An "absent" state on a nominal size of 5% still allows about twice that size. The verdict script stopped on a numerical overflow in its exact binomial interval; the amended copy changes only that computation and reproduces every line the frozen script printed before stopping (deposit, newpanel/P4L/AMENDMENT_1.md). Every count above was reproduced by an independently written parser, and every contrast state held under 21 further bootstrap seeds.

*S7.7 Class-separated branch models on the real null genes (exploratory)*

The plan (newpanel/P4T/PLAN.md), its scripts and a pre-run review were fixed by checksum before any fit of this design existed; the one-ratio and two-ratio fits of the same genes were already known, and only the new fits were blind. The branches of each real null gene were divided into four classes: the randomly labelled terminals of the registered analysis (R; 21 tips in fungi, null C; 19 in yeast; 18 in lactobacilli), the other terminals of reference species (O), the terminals of the trait species (X; the 21 ectomycorrhizal tips in fungi and the registered trait tips in yeast and lactobacilli) and the internal branches (I). Three codeml models (model = 2, NSsites = 0, the control file of the existing two-ratio fit otherwise unchanged) were fitted to each of the 88, 80 and 80 genes from $\omega = 0.5$ and 1.5, keeping the higher log-likelihood, 1,488 fits in all: TI, with one $\omega$ for internal and one for terminal branches; RX, with $\omega$ for I, for R and O together, and for X; and R4, with $\omega$ for I, R, O and X. The primary test was R4 against RX on one degree of freedom at 3.841; TI against the one-ratio fit tested the premise that internal and terminal branches differ. The states were fixed in advance: consistent with nominal (one-sided exact binomial $P \geq 0.05$ against 5% and a Clopper–Pearson upper bound of at least 5%), conservative (upper bound below 5%), above nominal ($P < 0.05$), or unresolved if more than 5% of genes were incomplete, more than 10% had starts differing by more than 0.01 log-likelihood units, or more than 5% had a statistic below -0.01. No flag was raised: every gene was complete, one fungal and one lactobacillus gene had starts differing by more than 0.01 units (at most 0.17), and no statistic was negative. Every count was reproduced by an independently written parser, which also confirmed the class marks on all 744 trees.

| **Panel (genes)** | **TI against one-ratio** | **R4 against RX (primary)** | **95% CI (%)** | **Two-ratio, existing** | **Internal > terminal $\omega$ in TI** |
|---|---|---|---|---|---|
| Fungi (88) | 48 (54.5%) | 35 (39.8%) | 29.5–50.8 | 58 (65.9%) | 67 |
| Yeasts (80) | 78 (97.5%) | 35 (43.8%) | 32.7–55.3 | 55 (68.8%) | 80 |
| Lactobacilli (80) | 79 (98.8%) | 23 (28.8%) | 19.2–40.0 | 71 (88.8%) | 80 |
| Pooled (248) | | 93 (37.5%) | 31.5–43.8 | | |

The primary test was above the nominal 5% of the $\chi^2_1$ reference in every panel and pooled (one-sided exact binomial $P \leq 5 \times 10^{-12}$), the outcome for which the plan allows only the statement that the excess persists among reference terminals and that its source is not identified here. Paired with the two-ratio test on the same genes, it rejected less often in each panel (McNemar exact $P$ = 0.0014, 0.00453 and $6.12 \times 10^{-13}$). The median class $\omega$ in R4 was 0.0380 for R and 0.0347 for O in fungi, 0.0582 and 0.0602 in yeast and 0.0317 and 0.0305 in lactobacilli, against 0.0622, 0.1119 and 0.1082 for internal branches. The analysis is not a calibration of the test under a correct null, and it does not identify why the reference terminals differ.

#### SUPPLEMENTARY S8 — SUPPORTING DETAIL FOR DISCUSSION 2 (WHAT TO DO INSTEAD)

*S8.2 Cross-program discordance under relabelling*

Supports the statement that the three programs failed under different conditions and on different genes.

Moving the label from terminal to internal branches took the error rate of one test from 22.5 to 13.5% (89 replicates fitted under both labellings) and that of the other from 10.0 to 47.0%, so no labelling scheme yielded nominal behaviour in both. Of the genes flagged by either RELAX or the two-ratio test under comparable criteria, only about a quarter were flagged by both. Agreement on how often these tests reject is not agreement on which genes they reject, and neither is evidence that they are measuring one thing.

*S8.3 Sampling error in the bootstrap threshold*

Supports the statement that an undersampled null can be less conservative than no correction at all.

Estimated from ten replicates the ninety-fifth percentile of the shallow set's null statistic ranged from 2.45 to 8.64 across resamples, and at the lower end of that range it falls below the $\chi^2_1$ critical value; estimated from eighty replicates the same percentile ranged only from 6.63 to 7.43. All figures in this subsection are for the shallow set; the divergence-matched set's percentile is 152.6 and is not comparable with them.

*S8.4 Choice of gene set for the reference distribution*

Supports the statement that the reference distribution should match the design, and that we cannot say whether it must also match the gene set.

Null distributions did not differ among the five genes used for simulation (Kruskal–Wallis $P$ = 0.31), nor detectably between the single-copy orthologues and forty enzyme families on a different gene set (35.8% of 81 against 35.0% of 40, a comparison whose 95% Wilson interval on the latter runs from 22.1 to 50.5% and so could not have resolved a difference smaller than the design effect), but differed substantially with the number of labelled tips (31.9 to 47.8%).

*S8.5 Compute cost of a design-matched reference distribution*

Supports the statement that the correction is affordable for a targeted analysis and not at present affordable at genome scale.

On our panel of 51 taxa a single RELAX fit took a median of 64 minutes, with the 81 null orthologues consuming 93 processor-hours in total. Fifty replicates for one design is therefore about 53 processor-hours, and the 539 required to reach a Bonferroni-corrected level across 27 tests about 570. Genome scale is far worse than a first estimate suggests, because the replicate count is set by the corrected significance level and not by the number of genes. Testing on the order of ten thousand genes puts the Bonferroni-corrected level at $5 \times 10^{-6}$, and $R \geq 1/\alpha - 1$ then demands about 200,000 replicates for each distinct labelling design, some 24 processor-years. Ten thousand replicates, the round number the scale of the analysis invites, would license only about five hundred tests. Deeper alignments are worse: the enzyme-family control, at 49 to 342 sequences, took a median of 168 minutes per fit.

A study whose tested families span several test fractions requires a separate null for each; ours span six, and each null is a complete re-analysis of the null gene set.

*S8.6 What is and is not new*

Supports the statement in Discussion 2 that the device is old and that the contribution is the dependence on the design.

What is and is not new should be stated exactly, because the device itself is old. That a likelihood-ratio test whose asymptotic distribution cannot be trusted should be referred to a parametric bootstrap is standard, and Halabi et al. (2021) applied precisely that to a test of this family, reporting type I error rates of 15 to 21% and correcting them by simulating under a fitted model. Three things separate what we report. Their inflation is measured on simulated data and attributed to the $\chi^2$ approximation and to uncertainty in the trait history; ours is measured on real genes under random labellings of varying size, and moves with the number of labelled tips (31.9 to 47.8%) while the gene set is held fixed. Their reference distribution is generated from a fitted model; the one we recommend must additionally reproduce the labelling design, which is the axis their construction holds constant and ours varies, and on real genes a reference generated from the fitted model did not restore the nominal rate even when it reproduced both the design and the depth of the data (Results 3). And their result concerns one statistic, where we find the same failure in two codebases (three implementations, of which two were run under both labellings); on the shallow set those two responded to the same change of design in opposite directions, while RELAX on the divergence-matched set and the two-ratio model across the 78 gene-fitted models each rejected more under internal labels. The contribution is not the bootstrap. It is that the quantity the bootstrap must condition on is the design, and that this had not been shown to matter.

Two further lines of work bear on the same tests and measure a different quantity. Hughes and Friedman (2008) showed that a branch-by-branch codon test returns apparent positive selection far more often on short branches, a property of the individual branch and the evidence it carries; the rate we report is a property of the set an analyst selects, with the alignment held fixed, though terminal and internal labellings also differ in the length of the labelled branches, so the branch-length dependence they describe may contribute to that contrast; we did not separate the two. Kowalczyk et al. (2021) showed that a branch-site result on an arbitrarily chosen foreground can reflect selection across the whole tree, and proposed removing the foreground as a check; that check cannot distinguish terminal from internal labelling, which is the axis along which our effect is measured. Both are complementary to what we report rather than prior instances of it.

*S8.7 Design-matched references, permutations and structure-preserving relabelling*

Supports the statement that matching the design is not the same as permuting the labels.

The recommendation needs stating precisely, because two constructions are easily conflated. What we evaluated in simulation is a reference distribution built by re-running the analysis on data generated under the same labelling design, with the labels held where the design puts them; on real genes that construction, simulated from the fitted model, did not restore the nominal rate even at the depth of the data, so matching the design is necessary but not sufficient (Results 3). Permuting the labels is a different construction, and Results 2 gives a reason not to substitute one for the other. A permuted labelling is not structure-preserving, and the pre-audit ectomycorrhizal set is measurably more clustered than a random draw. Structure-preserving relabelling is the intermediate case, and it is not the same construction either. Permulations (Saputra et al., 2021) draw labellings that retain the phylogenetic clustering of the observed one, and a permulation-style null of that kind (a Brownian trait labelled at its highest tips, whose clustering spans the observed value) is what we use on the count axis, where the question is whether the statistic is extreme among the labellings the tree could have produced. The selection axis puts a different quantity at fault. What we showed moves the null distribution there is how many branches are labelled, and the rejection rate tracks the number labelled with the gene set held fixed; which tips carry the label also mattered (null B against null C), but we cannot separate signal from structure in that contrast. A relabelling null varies the assignment while holding the design; the null we recommend holds the assignment and reproduces the design. Only the second conditions on the quantity that sets the error rate, which is why the two cannot be exchanged here even though the count axis is well served by the first.

**The forms of the correction, in full.** The correction buys size, not power, and in simulation only. A larger critical value moves the operating point along a fixed receiver-operating characteristic. On the divergence-matched replicates that loss is severe and on the shallower ones slight (Results 5, Figure 4b). On our real null genes at the same test fraction the ninety-fifth percentile of the null statistic is 33.2, against 152.6 on those matched alignments, so the losses we measured are the worst of them rather than the typical ones. A study should decide in advance whether an analysis with that operating characteristic is worth running, and the decision has a shape. The cost is worth paying when a false positive would be expensive to retract and the expected effect is large, where a study intends to name particular genes as relaxed and to build on that claim, as ours did. It is not worth paying when the analysis is exploratory and the output is a ranked list to be followed up experimentally, since there the uncorrected test is a screen and its error rate is absorbed by the follow-up. What is never defensible is the third case, and it is where we were: running the uncorrected test, reporting the survivors as findings, and leaving the reader to assume the nominal rate applies. The choice is between correcting and not claiming, not between correcting and claiming anyway.

Two forms of the correction are in circulation and they do not carry the same requirements, a distinction we were slow to draw ourselves. Referring the observed statistic to a percentile of the null distribution, the form the simulated evaluation of Results 5 assesses, has no lower bound on the *P* value it can support, but an undersampled null can be less conservative than no correction at all: estimated from ten replicates, the ninety-fifth percentile of our shallow null statistic spanned 2.45 to 8.64 across resamples and fell below the $\chi^2_1$ critical value in part of that range (Supplementary S8.3). Reporting an empirical *P* value does have a bound, $1/(R + 1)$ from $R$ replicates, so $R \geq 1/\alpha - 1$: nineteen replicates for a single test at 0.05 and 539 for 27 tests at a

Bonferroni-corrected level. The reference distribution should match the design, and we cannot say whether it must also match the gene set: null distributions differed substantially with the number of labelled tips (31.9 to 47.8%) while a forty-family control on a different gene set returned a rate we could not distinguish from null C, though with forty families that control could not have detected a gene-set effect much smaller than the design effect itself (Supplementary S8.4).

We are nonetheless not proposing a device of our own. Contrast-FEL performs a label-permutation test on significant sites by default, and on our null genes that step took the site-level rejection rate from 9.18 to 0.478%, below the nominal level rather than above it (S5.3), evidence that referring the statistic to a reference distribution generated from the data rather than to $\chi^2$ is practicable, in a program that ships with it, although a permutation is not the construction we evaluated. A step of this kind establishes that a finding is not an artefact of the asymptotic approximation; at the resolution Contrast-FEL ships with, it cannot license a claim at a corrected threshold (S5.3). The recommendation here is that the two programs lacking any such step acquire one. We do not rank the three, since one rate is measured on sites and two on genes.

*S8.8 A worked example: four published RELAX tests read against same-shaped labels*

Supports the worked example in Discussion 2.

Zahnow et al. (2025) tested the four epithelial sodium channel subunit genes (*SCNN1A*, *B*, *G* and *D*) of up to 91 mammals for relaxed selection in marine lineages with RELAX, and deposited their alignments, labelled trees, command and 23 RELAX outputs (Source Data 6, Zenodo 10.5281/zenodo.15255678). We read only tests whose test set is a single terminal branch or a two-tip clade, the only shapes with more than one alternative of the same shape among the reference branches of these trees, and of those only the four the published output calls significant:

*SCNN1A* and *SCNN1G* in the otter (one terminal branch) and *SCNN1A* and *SCNN1D* in the sirenians (a two-tip clade). Selection on published significance is by design, since the question is how a published positive reads against the same tree's other lineages; no rate over published tests is estimated. The plan, the scripts and the list of 131 runs were fixed by checksum before any run whose result is read, and the inputs were byte-identical to the deposit. Each run used the deposited alignment and tree and the published command in HyPhy 2.5.101 with a fixed seed and one thread, except that the reference set was named where this release refuses the published argument; the test, reference and nuisance sets of every run that completed were checked by branch name against the published output. A published label counted as reproduced when at least two of three seeds gave $P \leq 0.05$ with $K$ on the published side of 1, and only then were its alternatives read, one seed each. An alternative moves the test label to another lineage of the same shape whose branches are all reference branches, leaving the published test branches outside both sets.

*SCNN1A*, otter (published $P = 0.0037$, intensification, $K = 50$, the upper bound): not reproduced, with $P = 0.36$–$0.45$ and $K = 1.30$–$1.31$ under the three seeds. The published alternative fit has a log-likelihood of -22282.24 against -22287.80 to -22288.21 in ours, on the same data and parameter count, so our runs missed the published optimum; our null fits (-22288.21 to -22288.50) are also about 2 units below the published null (-22286.45), which points to a difference of optimiser or HyPhy release. This is not evidence against the published result.

*SCNN1G*, otter (published $P = 0.0323$, intensification, $K = 8.90$): not reproduced in direction, with $P = 0.027$–$0.029$ and $K = 0.06$–$0.08$ under all three seeds. The published output contains, after the fit with $K = 8.90$ (log-likelihood -19895.85), a fit in the other direction of $K$ with a higher log-likelihood (-19895.66, $K = 0.05$); the reported $P$ value and direction correspond to the first, after RELAX's check for a flat likelihood surface. Our seeds agree with the better fit in direction, at log-likelihoods (-19896.12 to -19896.41) below both published alternative fits.

*SCNN1A*, sirenians (published *P* = 0.0072, relaxation, *K* = 0.64): reproduced under all three seeds (*P* = 0.0072–0.0078, *K* = 0.63–0.65). Its likelihood-ratio statistic, 7.23 with each model taken at its highest log-likelihood over the three seeds, exceeded that of each of the 12 same-shaped alternatives (largest 1.02), rank 1 of 13, and none of the alternatives would be called significant at $\chi^2_1$. With 13 labels no rank-based *P* value below 1/13 is attainable. Eleven of the 12 alternatives are shorter than the published lineage (total length 0.051 against 0.001–0.102 in the tree's units), and the two within a factor of two of its length are both below it.

*SCNN1D*, sirenians (published *P* = 0.0118, relaxation): not evaluable. HyPhy rejected the deposited alignment, in which all 68 sequences carry in-frame stop codons; the published runs used an edited alignment of 704 codons, against 723 in the deposited one, which is not in the deposit.

An independent recount reproduced the statistics, *P* values, *K* values, branch sets and ranks from the run outputs, and an audit checked the log-likelihoods against the published outputs.

#### SUPPLEMENTARY TABLE S9

**The 51 genomes carrying the codon alignments used in every selection analysis.** These are the species with coding sequences recoverable in a form suitable for codon-aware alignment; they are a subset of the 183-genome annotation panel and of the 182-tip phylogenetic panel, and they are not a random subset of either (Discussion 4). "Lifestyle (used here)" is the assignment under which every selection analysis and every null set in this paper was built. "Lifestyle (corrected)" is filled only for the three genomes the annotation audit of Results 1 reassigned, and shows what they would be under the corrected assignment; the selection analyses were not re-run on it, and Methods M5 states why. Under the assignment used here the test set is 21 ectomycorrhizal tips of 51; under the corrected one it would be 18. "Public" records whether the genome's portal page carried an associated publication at the time of analysis: 35 of the 51 were, and 16 were not. Portal codes are the stable JGI MycoCosm identifiers and are sufficient to retrieve each genome.

| **Portal** | **Species** | **Lifestyle (used here)** | **Lifestyle (corrected)** | **Public** |
|---|---|---|---|---|
| Altal1 | *Alternaria alternata* SRC1lrK2f v1.0 | Pathogen | — | yes |
| Amagr1 | *Amanita* aff. *grandis* KIS10 v1.0 | Ectomycorrhizal | — | **no** |
| Amamu1 | *Amanita muscaria* Koide v1.0 | Ectomycorrhizal | — | yes |
| Amapyr1 | *Amanita* aff. *conicoverrucosa* KIS12 v1.0 | Ectomycorrhizal | — | **no** |
| Amath1 | *Amanita thiersii* Skay4041 v1.0 | Saprotroph | — | yes |
| Amore1 | *Amorphotheca resinae* v1.0 | Saprotroph | — | yes |
| Ascim1 | *Ascobolus immersus* RN42 v1.0 | Saprotroph | — | yes |
| Aurde3_1 | *Auricularia subglabra* v2.0 | Wood decayer | — | yes |
| Aurpu_var_pul1 | *Aureobasidium pullulans* var. *pullulans* EXF-150 | Saprotroph | — | yes |
| Bissp1 | *Bisporella* sp. PMI_857 v1.0 | Saprotroph | — | yes |
| Bolcoc1 | *Boletus coccyginus* 2016PMI039 v1.0 | Ectomycorrhizal | — | **no** |
| Botbo1 | *Botryobasidium botryosum* v1.0 | Saprotroph | — | yes |
| Cananz1 | *Cantharellus anzutake* C23 v1.0 | Ectomycorrhizal | — | yes |
| Cenge3 | *Cenococcum geophilum* 1.58 v2.0 | Ectomycorrhizal | — | yes |
| Chove1 | *Choiromyces venosus* 120613-1 v1.0 | Ectomycorrhizal | — | yes |
| ClaPMI390 | *Clavulina* sp. PMI_390 v1.0 | Saprotroph | — | yes |

| Portal | Species | Lifestyle (used here) | Lifestyle (corrected) | Public |
| --- | --- | --- | --- | --- |
| Conpu1 | *Coniophora puteana* v1.0 | Wood decayer | — | yes |
| Dacsp1 | *Dacryopinax primogenitus* DJM 731 SSP1 v1.0 | Wood decayer | — | yes |
| ElagrMar1 | *Elaphomyces granulatus* Maridalen v1.0 | Ectomycorrhizal | — | **no** |
| Fibsp1 | *Fibulorhizoctonia* sp. CBS 109695 v1.0 | Saprotroph | — | yes |
| Gloin1 | *Rhizophagus irregularis* DAOM 181602 v1.0 | Arbuscular mycorrhizal | — | yes |
| Gyman1 | *Gymnopus androsaceus* JB14 v1.0 | Saprotroph | — | yes |
| Hebcy2 | *Hebeloma cylindrosporum* h7 v2.0 | Ectomycorrhizal | — | yes |
| Hetan2 | *Heterobasidion annosum* v2.0 | Pathogen | — | yes |
| Hyafin1 | *Hyaloscypha finlandica* PMI_746 v1.0 | Ericoid mycorrhizal | — | **no** |
| Hyavar1 | *Hyaloscypha variabilis* MD1 v1.0 | Ericoid mycorrhizal | — | **no** |
| Hypsu1 | *Hypholoma sublateritium* v1.0 | Wood decayer | — | yes |
| Hyssto1 | *Hysterangium stoloniferum* HS.BST v1.0 | Ectomycorrhizal | — | yes |
| Jaaar1 | *Jaapia argillacea* v1.0 | Wood decayer | — | yes |
| Kurarg1 | *Kurtia argillacea* OMC1749 v1.0 | Ectomycorrhizal | Ericoid mycorrhizal | **no** |
| Lacaka1 | *Lactarius akahatsu* QP v1.0 | Ectomycorrhizal | — | yes |
| Lacbi2 | *Laccaria bicolor* v2.0 | Ectomycorrhizal | — | yes |
| Matter1 | *Mattirolomyces terfezioides* MAT_tt4AIII v1.0 | Ectomycorrhizal | — | **no** |
| MelPMI1271_1 | *Meliniomyces* sp. PMI_1271 v1.0 | Ericoid mycorrhizal | — | **no** |
| Melbi2 | *Meliniomyces bicolor* E v2.0 | Ericoid mycorrhizal | — | yes |
| Morco1 | *Morchella importuna* CCBAS932 v1.0 | Ectomycorrhizal | Saprotroph | yes |
| Neole1 | *Neolentinus lepideus* | Wood decayer | — | yes |

| **Portal** | **Species** | **Lifestyle (used here)** | **Lifestyle (corrected)** | **Public** |
|---|---|---|---|---|
| | v1.0 | | | |
| Neucr_trp3_1 | *Neurospora crassa* FGSC 73 trp-3 v1.0 | Saprotroph | — | yes |
| Paxin1 | *Paxillus involutus* ATCC 200175 v1.0 | Ectomycorrhizal | — | yes |
| Penswi1 | *Penicillium swiecickii* 182_6C1 v1.0 | Ectomycorrhizal | Saprotroph | **no** |
| Phchr2 | *Phanerochaete chrysosporium* RP-78 v2.2 | Wood decayer | — | yes |
| Phlbr1 | *Phlebia brevispora* HHB-7030 SS6 v1.0 | Wood decayer | — | yes |
| Ruscya1 | *Russula cyanoxantha* BavarianF04 v1.0 | Ectomycorrhizal | — | **no** |
| Ruseme1 | *Russula emetica* Přilba v1.0 | Ectomycorrhizal | — | **no** |
| Rusoch1 | *Russula ochroleuca* Přilba v1.0 | Ectomycorrhizal | — | **no** |
| Sebve1 | *Sebacina vermifera* MAFF 305830 v1.0 | Orchid mycorrhizal | — | yes |
| Ser400_1 | *Serendipita* sp. 400 v1.0 | Orchid mycorrhizal | — | **no** |
| Ser405_1 | *Serendipita* sp. 405 v1.0 | Orchid mycorrhizal | — | **no** |
| Serend1 | *Serendipita* sp. 407 v1.0 | Orchid mycorrhizal | — | **no** |
| Thacu1 | *Rhizoctonia solani* AG-1 IB | Pathogen | — | yes |
| Theter1 | *Thelephora terrestris* UH-Tt-Lm1 v1.0 | Ectomycorrhizal | — | yes |

Penswi1 was recorded at the time of download as *Penicillium* sp. GLBRC 242 v1.0, which is a different assembly (31.35 Mb, 11,517 gene models); the files analysed here are the 34.13 Mb, 12,802-model assembly of *Penicillium swiecickii* 182_6C1 v1.0, and the name in this table is corrected accordingly. Its pre-audit lifestyle label, ectomycorrhizal, was inherited from that other assembly's portal record rather than read from the record of Penswi1 itself, so it is not evidence that the portal assigns *P. swiecickii* to ectomycorrhizal fungi. The lifestyle audit and every count are unaffected, since they were made on the files.

The count analyses do not depend on the unpublished genomes: restricting the primary contrast to public genomes gives 0.411 against 0.367 (Results 1). The selection analyses do depend on them, and cannot be reproduced from public data alone.

#### Supplementary figures

##### Figure S1

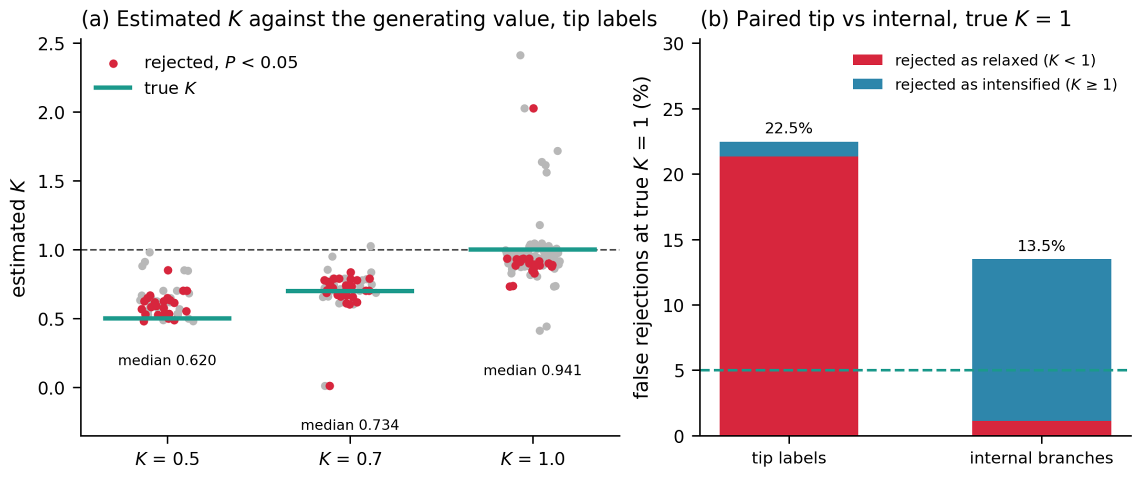
